# REZCUE: 3D spatial multi-omics profiling from archived FFPE tissue

**DOI:** 10.64898/2026.10.06.757132

**Authors:** Lichun Zhang, William Chi-Shing Cho, Joshua Jing Xi Li, Brian Pak Shing Pang, Ben Tin Yan Wong, Eldric Pui Lam Tsoi, Yuqi Tan, Hei-Ming Lai

## Abstract

Archived formalin-fixed, paraffin-embedded (FFPE) tissue represents a vast clinical resource, yet three-dimensional imaging and omics-wide molecular profiling remain largely incompatible because current antigen retrieval and clearing can degrade nucleic acids. We developed REZCUE, a depth-invariant antigen retrieval chemistry that enables 3D multiplexed immunofluorescence imaging of intact FFPE tissue while quantitatively preserving DNA, RNA, and protein for multi-omics analyses. REZCUE restores antigenicity in tissue archived for up to 30 years, supports 160 commercial whole-IgG antibodies across tissue types, and enables imaging to 6 mm depth. Imaging before destructive molecular profiling establishes a 3D spatial reference for registering molecular assays to intact tissue architecture. REZCUE enabled multi-modal mapping of spatial niches of the mouse brain and identified a human breast tumor compartment not distinguishable by conventional section-based spatial-omics, revealed after integrating 3D image-guided segmentation with spatial transcriptomics, mass spectrometry, miRNA sequencing, and whole-genome sequencing. In short, REZCUE transforms archived FFPE specimens into spatially registered, molecularly resolved, 3D substrates for sequential spatial multi-omic interrogation.

**Graphical abstract:** 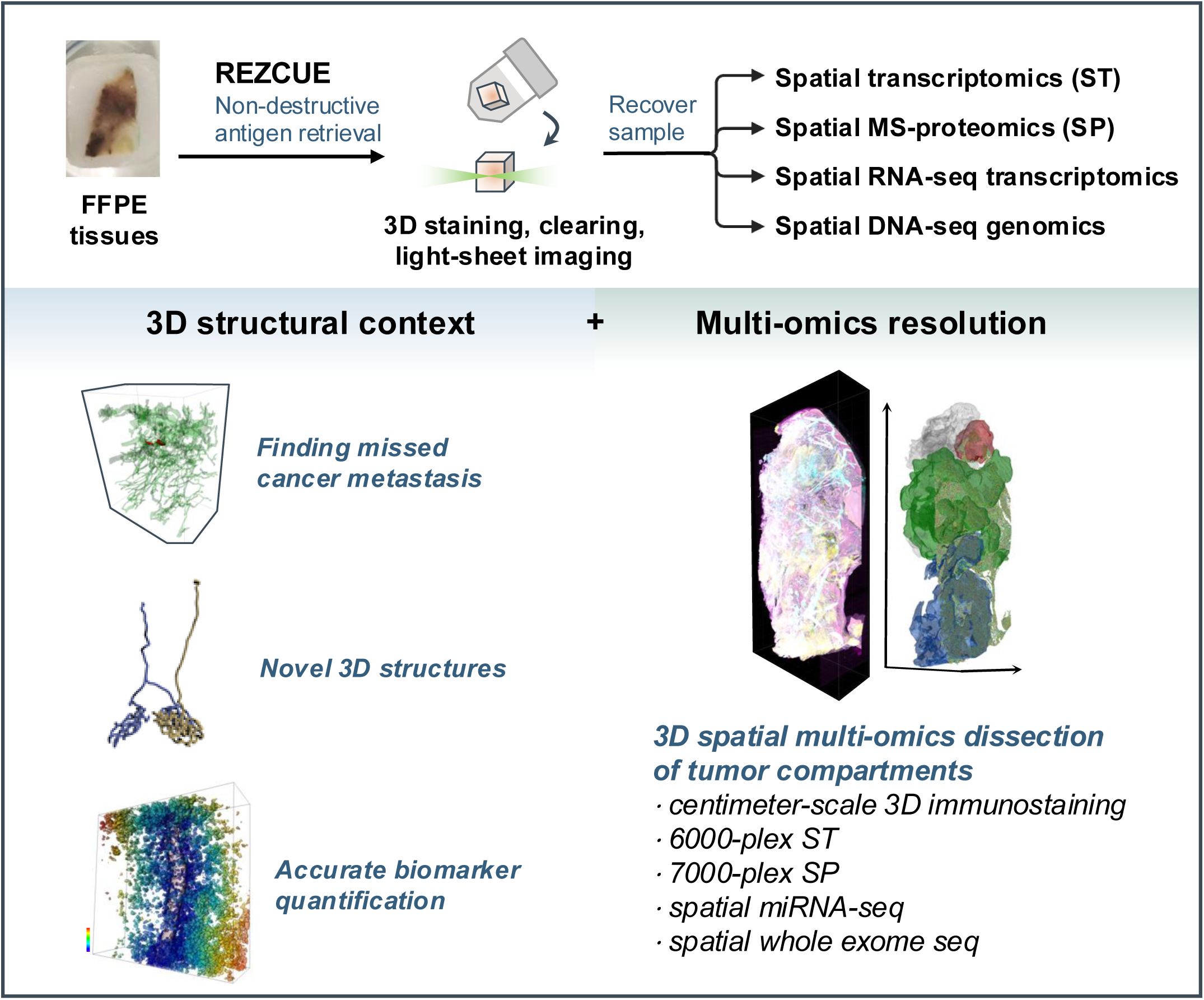

## Introduction

Understanding the mechanisms underlying complex diseases requires statistically powered, systems-level molecular profiling from large patient cohorts with data on long-term clinical outcomes. Archived clinical specimens are uniquely suited to this goal ^1^. These archived tissue collections exist mostly in formalin-fixed, paraffin-embedded (FFPE) format, and many of these pathology archival tissue blocks are linked to decades of clinical outcome data from trials and cohort studies ^2^. They allow retrospective cohorts with known treatment responses and outcomes to be assembled in months rather than the years a prospective study requires, and they capture disease directly in patients. Each block is a volumetric snapshot of human pathology but exists in finite quantity. Extracting the maximum molecular information from every specimen, while retaining its 3D morphological context, is therefore central to make full use of these archives and decode multifactorial diseases.

However, working with archived FFPE specimens presents two fundamental challenges. First, conventional assays such as traditional H&E, immunohistochemistry (IHC) require physical sectioning of the FFPE blocks, which consumes the tissue and discards its 3D context. A single 4-5 µm section samples well under 1% of a typical 5 mm-thick block, so rare events, such as micrometastases, lymphovascular invasion or focal biomarker expression, are easily missed ^3^. Continuous structures, including vessels, ducts and invasion fronts, could cross multiple sections; cell counts, volumes and spatial relationships measured in one plane can misrepresent the specimen as a whole ^4^. Stereology and serial sectioning with computational reconstruction can recover some of this information, but they remain limited by sectioning artefacts, registration error and throughput ^3,5–8^. In contrast, 3D imaging of intact, cleared specimens bypasses sectioning altogether, enabling complete sampling of tissue architecture, provided that the involved chemistry does not significantly destroy tissue biomolecules.

Because each FFPE specimen is finite, the value of 3D imaging is greatest when the tissue remains usable for subsequent molecular analysis; otherwise, volumetric imaging comes at the cost of every other assay the specimen could have supported. At present, researchers must choose one or the other: high-parameter molecular assays require thin tissue sections, while volumetric imaging has not been compatible with subsequent molecular assays.

To our knowledge, no method has combined 3D immunostaining of FFPE tissues with downstream multi-omics, single-cell or spatially resolved molecular assays ^9–11^. Most established clearing and deep-immunostaining methods were developed for paraformaldehyde (PFA)-fixed, non-embedded tissues and do not directly transfer to FFPE tissues ^12,13^. Methods adapted or developed for FFPE tissues, including DIPCO ^14^, HIF-clear ^15^, aDISCO ^9^, OPTIClear ^10^, FIDELITY ^16^, and miDISCO+ ^17^, achieved volumetric immunostaining but rely on heat-induced retrieval, strong detergents, strong acids or bases. None has demonstrated compatibility of downstream molecular assays such as single-cell or spatial transcriptomics, and genomic sequencing. Conversely, DISCO-seq and DISCO-MS combine 3D immunostaining with downstream transcriptomics and mass spectrometry, respectively, but were developed for fresh-frozen tissue ^18,19^ Only DISCO-MS is compatible with anti-GFP nanobody immunostaining, but not with the whole IgG antibody immunostaining of endogenous antigens ^18^. A recent 3D pathology-guided microdissection (3DPM) approach shows that whole-genome sequencing can be performed on archival FFPE tissue after volumetric imaging, by physically milling out annotated sub-volumes for bulk sequencing, but the imaging step relies on small molecular dyes rather than antibody-based imaging ^20^. Extending the compatibility of antibody-based 3D immunostaining with downstream multi-omic assays to archival FFPE specimens, which represent the largest repository of clinically annotated human tissue, is therefore critical to unlocking their untapped biological and informational potential.

Here we present REZCUE (for <u>R</u>escuing <u>E</u>pitopes with Zwitterionic detergent and super<u>C</u>haotropes for <u>U</u>tilizing <u>E</u>mbedded tissues), a non-destructive chemistry that achieves deparaffinization, deep and uniform 3D antigen retrieval, immunostaining and clearing of intact FFPE tissue, while preserving DNA, RNA, and other biomolecules for downstream multimodal omics-wide profiling (**Figure 1A-B**). REZCUE replaces Heat-Induced Epitope Retrieval (HIER) with sequential, mild solution incubations, and achieves penetration for millimeter-to-centimeter-scale FFPE blocks. REZCUE also restores antigenicity in FFPE tissues archived for up to 30 years, and preserves tissues for spatial transcriptomics, mass-spectrometry-based proteomics, RNA sequencing, and whole-genome sequencing on the same tissue block after 3D imaging. Applied to mouse brain and a 25-year-old human breast tumor, REZCUE enabled multimodal mapping of spatial niches and resolved a luminal B tumor compartment that was not distinguishable by conventional section-based analysis. REZCUE thus enables 3D-resolved, multi-omic datasets from archived specimens, allowing retrospective studies and reanalysis of tissues as new questions and technologies emerge.

**Figure 1.**
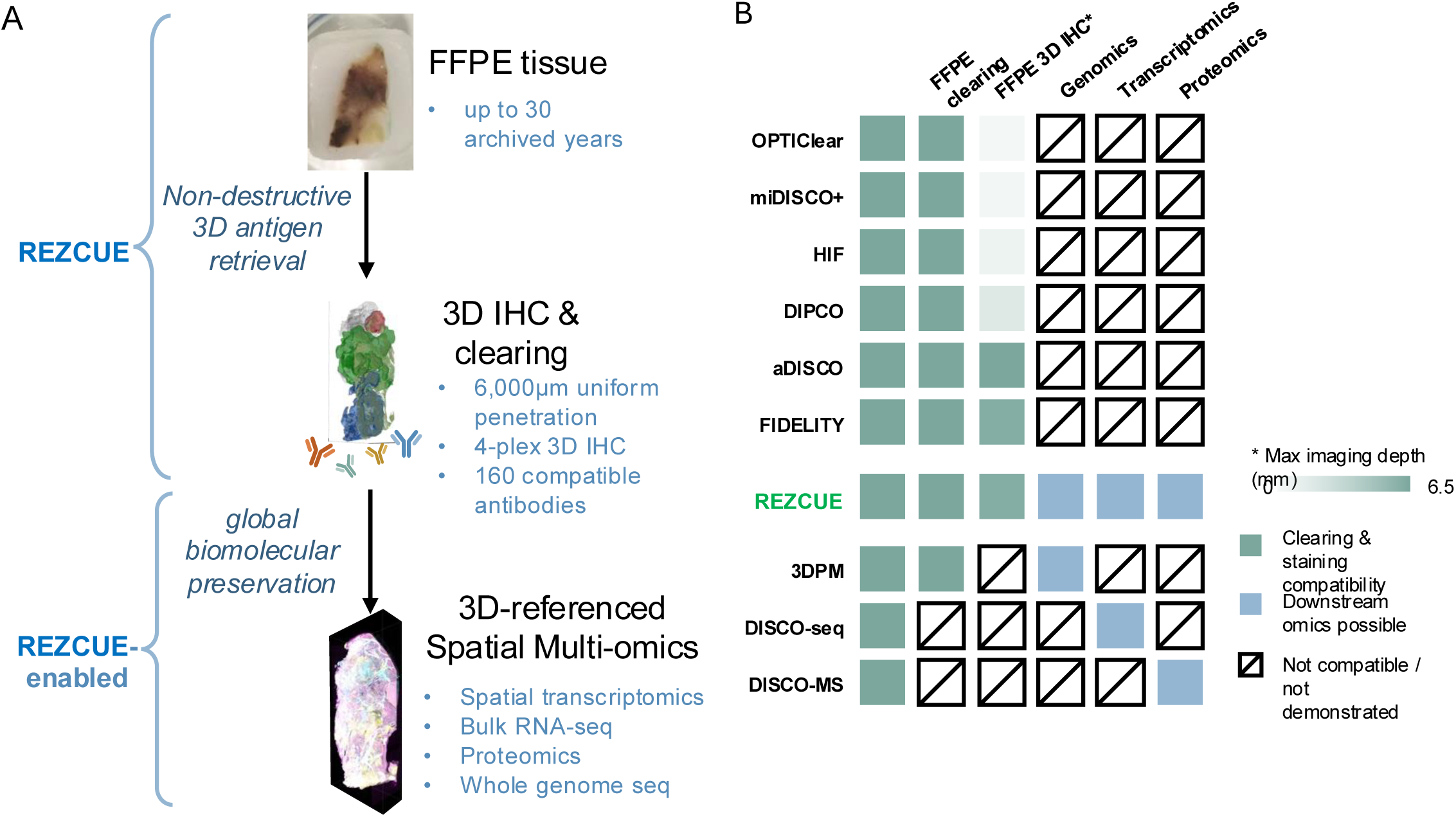
REZCUE methodology. **A**. Overview of REZCUE, which enables deeply and uniformly penetrating 3D immunostaining in FFPE samples archived for up to 30 years, while preserving biomolecules for downstream spatially resolved multi-omics. **B.** The unique technological gap addressed by REZCUE compared with methods demonstrating FFPE 3D immunostaining, and omics profiling methods after tissue clearing but not immunostaining.

## Results

### REZCUE retrieves antigens in archival 3D FFPE blocks in a non-destructive way

Traditional FFPE retrieval involves deparaffinization with xylene, followed by various digestive conditions involving heat (95 - 121 °C), denaturing detergents, protease digestion, or strong acid (pH < 1) or alkaline (pH > 14) treatment (**Figure 2A**) ^21^. Harsh chemical treatment was deemed necessary for antigen retrieval because excessive formaldehyde crosslinking was believed to be the primary cause of epitope masking; however, these treatments aimed at breaking covalent crosslinks also cause biomolecule loss ^22^. It is unclear whether harsh retrieval conditions are necessary. Fixed tissue that was not paraffin-embedded stains well with many antibodies, whereas lightly fixed tissue stains poorly once embedded in paraffin ^23^. This suggests that crosslinking is not the main driver of epitope masking.

**Figure 2.**
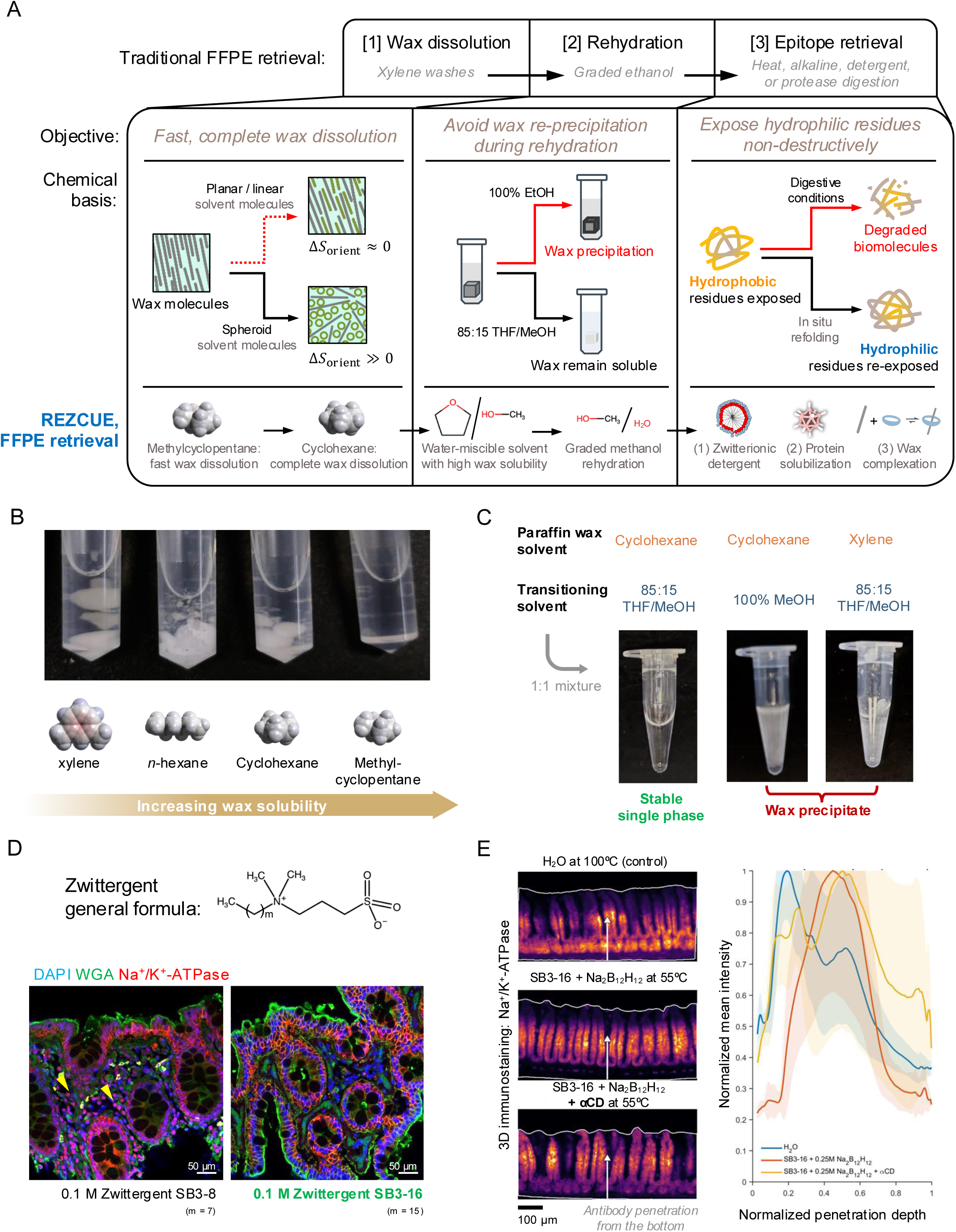
Principles of REZCUE chemistry. **A**. Complete paraffin wax extraction, avoiding wax reprecipitation in tissue matrix, and non-destructive protein refolding are crucial aspects in the REZCUE development. A comparison between traditional antigen retrieval and REZCUE antigen retrieval is provided, along with their chemical basis. **B**. Solubility of 1g paraffin wax in various 500 µl organic solvents within 30 minutes at 20-25 °C. Methylcyclopentane is fastest in dissolving paraffin wax. **C.** Paraffin wax-saturated solvents (cyclohexane or xylene) were mixed in 1:1 ratio with a transitional solvent. Only a solvent blend of 85:15 v/v THF/MeOH mixed with wax-saturated cyclohexane did not result in wax reprecipitation. **D**. Structural optimization of sulfobetaine-based aliphatic zwitterionic detergent (SB3-X, where X is the aliphatic chain length). The use of 0.1 M SB3-8 with an *n*-octyl tail as antigen retrieval reagent resulted in the appearances of non- specific nuclear staining (yellow arrows) for anti-Na^+^/K^+^-ATPase antibodies (in red), whereas lengthening the aliphatic chain to a palmitoyl tail eliminated it for the case of SB3-16. **E**. 5 mm-long FFPE mouse colon 3D immunostaining with anti-Na^+^/K^+^-ATPase antibodies after various antigen retrieval treatments. The addition of alpha-cyclodextrin (αCD) led to full-thickness penetration. Left: representative images of a z-slice (white arrow: antibody penetration direction from serosa towards mucosa); right: normalized mean intensities (lines) with S.D.s (shaded regions) plotted against normalized distance transformation coordinates from surface at 0 to the deepest region of 1.

To develop a new non-destructive chemistry, we hypothesized that proteins are folded “inside-out” when embedded in a highly hydrophobic medium under heat, burying hydrophilic epitopes while exposing hydrophobic amino acid residues on the surface. Traditional FFPE antigen retrieval may therefore work by facilitating re-exposure of hydrophilic epitopes once the tissue returns to an aqueous environment, where the breakage of covalent bonds can introduce more degrees of freedom within the macromolecules for epitope re-exposure (see further discussions in **Supplementary Note 1**). If this hypothesis is true, we reasoned that by facilitating protein refolding through alternative means, antigen retrieval can be achieved without destroying biomolecules.

In contrast to the essentially 2D case of tissue sections, complete deparaffinization of intact FFPE tissue volumes is much more difficult and any residual wax can thermodynamically favor the masking of hydrophilic epitopes. Our first step was to develop a method for mild tissue deparaffinization. For decades, tissue deparaffinization relied on xylene or other alkenes (such as limonene or terpenes) to dissolve wax. We found xylene to be a poor solvent for paraffin wax, possibly because the π-π-interactions between xylene molecules made them enthalpically unfavorable to dissolve linear aliphatic alkanes in paraffin wax (**Supplementary Figure 1A**). Interestingly, changing xylene to the linearly shaped *n*-hexane also performed poorly despite favorable enthalpic interactions with paraffin wax molecules and lower molecular weight for diffusive infiltration. We reasoned that *n*-hexane provided limited orientational entropy gain during wax dissolution, which can be mitigated by using a more compact alkane such as cyclohexane and methylcyclopentane (**Supplementary Notes 1-2**). Treatment of FFPE tissue with methylcyclopentane and cyclohexane resulted in rapid, complete tissue deparaffinization (**Figure 2B, Supplementary Figure 1A, Supplementary Notes 1-2**). Since paraffin wax is poorly soluble even in anhydrous alcohols, we also recognized the need to minimize paraffin wax reprecipitation as the tissue is transferred into washing alcohol solvents in preparation for rehydration. After extensive screening, a transitional solvent washing step with an 85%/15% v/v tetrahydrofuran/methanol solvent blend was used to minimize paraffin wax reprecipitation as the tissue transitions from the deparaffinization solvent to methanol (**Figure 2C, Supplementary Figure 1B, Supplementary Note 1**).

Next, we identified conditions that promote in situ protein refolding. After exploring diverse conditions (**Supplementary Figure 2**), we discovered incubating FFPE tissues in an aqueous solution containing detergents and superchaotropes is a promising candidate for non-covalent antigen retrieval. The superchaotropic dodecahydro-*closo*-dodecaborate ion ([B_12_H_12_]^2-^) is known to reduce non-specific binding by preventing re-aggregation of proteins by temporarily coating macromolecular surfaces ^24^. We screened for the most suitable detergent for their ability to recover expected immunostaining patterns in the presence of [B_12_H_12_]^2-^ (**Supplementary Figure 3A**). We found that zwitterionic detergents, especially alkyl zwitterionic detergents, were especially promising and proceeded to its systematic structural optimization. We found the palmitoyl variant, *n*-hexadecyl-*N*,*N*- dimethyl-3-ammonio-1-propanesulfonate (Zwittergent SB3-16), resulted in the best signal-to-background ratio (**Figure 2D, Supplementary Figure 3B**). Meanwhile, we confirmed that the presence of [B_12_H_12_]^2-^ was crucial for reduction of non-specific staining (**Supplementary Figure 4A**). Even when treated with both [B_12_H_12_]^2-^ and Zwittergent SB3-16, FFPE tissue retains small amounts of paraffin that reduce antibody penetration uniformity. Therefore, we added α-cyclodextrin (αCD) to further increase the staining penetration depth; αCD creates water-soluble supramolecular complexes with paraffin wax molecules to minimize residual hydrophobic regions (**Figure 2E**) ^25^, as well as including an immediate 50% v/v DMSO post-wash (**Supplementary Figure 4B**). Combining these specifically designed chemical approaches, we arrived at REZCUE, a mild antigen retrieval method that contrasts sharply with traditional antigen retrieval, which uses digestive conditions to break covalent bonds and re-expose epitopes (**Figure 2A**). REZCUE presents a new non-destructive chemistry that fills an important technological gap where the same physical archived specimen can be 3D-imaged with multiple molecular markers using deep, uniformly penetrating immunostaining, followed by downstream multi-omics analysis of the same tissue.

### REZCUE enables large-scale 3D imaging of long archival FFPE specimens

To evaluate REZCUE’s ability to recover high-quality 3D immunostaining images from FFPE samples, we stained and imaged human colon, liver, and appendix FFPE tissue blocks archived for 1, 10, or 30 years (**Figure 3A-B, Supplementary Figure 5-6**). After immunostaining, staining quality was assessed using confocal microscopy (40×) at 50-120 µm depths from the tissue surface. Tissues retrieved using the standard 2D FFPE retrieval process (xylene deparaffinization followed by HIER) showed weak or no staining compared with fresh-fixed, non-paraffin-embedded controls (**Figure 3A, Supplementary Figure 5A, B**). However, REZCUE yielded significantly higher signal-to-background ratios in the same blocks (**Figure 3A, B**), and the staining quality was comparable to images from fresh-fixed, non-paraffin-embedded controls (**Supplementary Figure 5A, C**). To test the generality of REZCUE, we also validated 160 commercially available, whole IgG antibodies on 11 types of REZCUE-processed tissues (**Supplementary Figure 7, Supplementary Table 1**) and cross-referenced it with their expected staining patterns in the specified tissues with the Human Protein Atlas ^26^.

**Figure 3.**
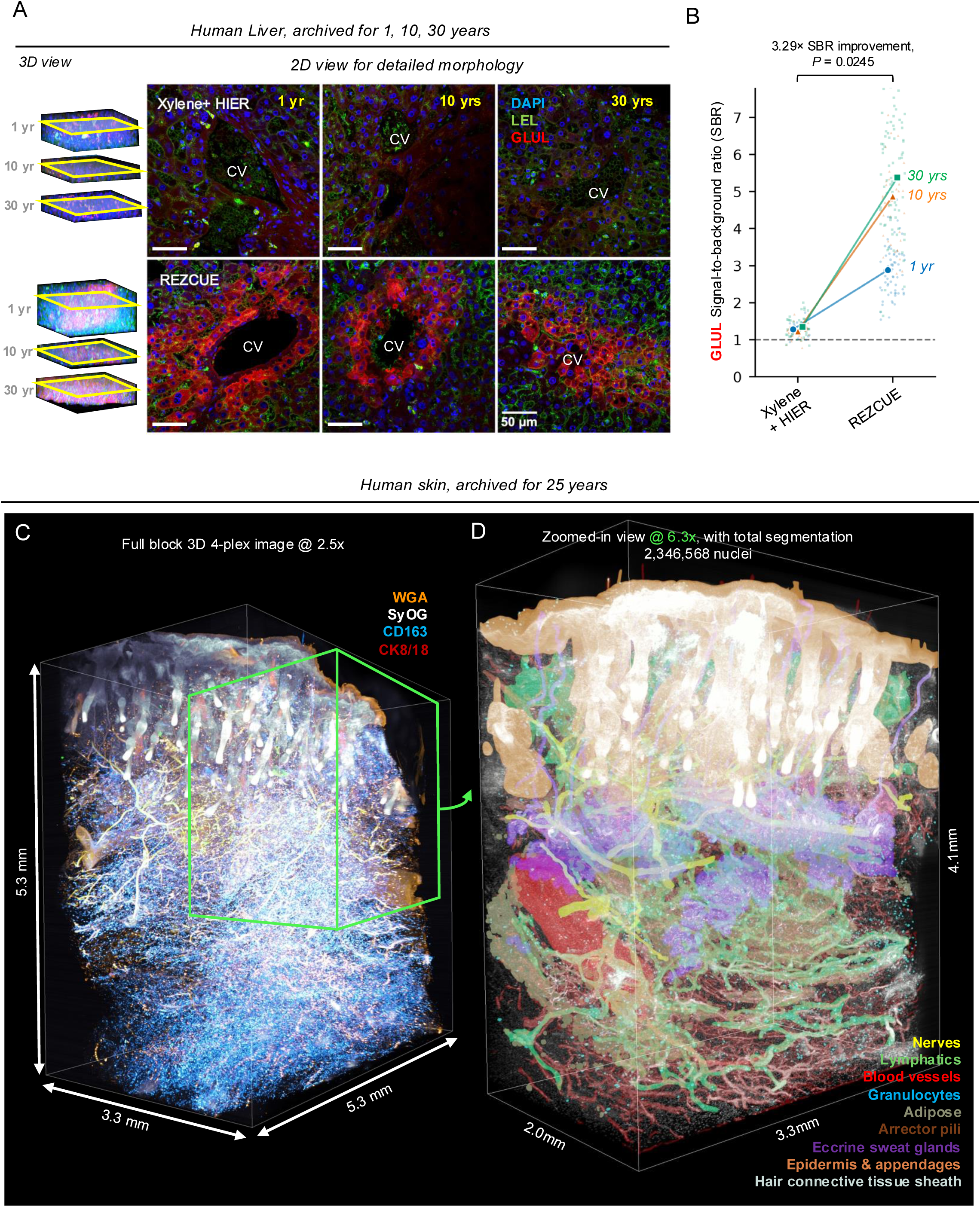
REZCUE enables high-quality 3D histology in challenging samples. **A.** Comparison of immunostaining quality at 50-120 µm deep for samples archived for 1, 10, and 30 years with traditional antigen retrieval (xylene deparaffinization plus HIER; upper panel) versus REZCUE (lower panel). **B.** Quantified GLUL immunostaining signal-to-background ratio at 100-120 µm depth of imaging and at single-cell level. Each small dot represents the data of a segmented peri-central vein hepatocyte GLUL. Larger dots with connected lines represent the paired specimens’ means. Across all paired specimens, a 3.29× signal-to-background ratio improvement was noted with REZCUE compared to conventional xylene + HIER antigen retrieval. *P*-value: geometric mean comparison of all 3 samples in each category using two-tailed, two-sample *t*-test. **C.** LSM image of a 5.3 mm × 5.3 mm × 3.3 mm human skin tissue sample at 2.5× magnification stained with wheat germ agglutinin (WGA), SYTOX Orange (SyOG), anti-CD163 antibodies and anti-CK8/18 antibodies. **D.** Zoomed-in view of panel **C** at 6.3× magnification with machine learning-based segmentation of major mesoscopic structure in the skin, identified by differential color coding. See **Supplementary Video 1** for 3D rendering.

To further assess the quality of 3D multiplexed immunohistochemistry staining after REZCUE treatment, we imaged human gastric, colon, appendix, lung, liver, pancreas, tongue, and urinary bladder tissues by light-sheet fluorescence microscopy (LSFM) (**Supplementary Figures 8-11**). In these tissues, we observed expected features and successful deep multiplexed 3D immunostaining. In a 1.5 mm × 5.5 mm × 3.5 mm skin squamous cell carcinoma, pan-cytokeratin positive keratin pearls can be found with pleomorphic cell morphologies (**Supplementary Figure 8A**). In a 2.5 mm × 8 mm × 5 mm human lung adenocarcinoma, we observed the densely cellular tumor compressing and filling up the normal alveoli (**Supplementary Figure 8B**), highlighted by 3D CAV1 immunostaining. In a human urinary bladder tumor tissue block (3 mm × 4 mm × 2 mm), we observed lymphoid follicles adjacent to the tumor invasive front, a common feature of urothelial carcinomas (**Supplementary Figure 9**).

The largest imaged sample obtainable from our archives was a 10 mm × 6 mm × 6 mm gastric biopsy tissue, where two lectins and the densely expressed antigen VIM were successfully labeled throughout the entire tissue thickness (**Supplementary Figure 10A**).

Challenging tissues such as the human tongue (3.5 mm × 4 mm × 3 mm) was also successfully stained and cleared (**Supplementary Figure 10B**). We also imaged a human appendix tissue block (10 × 3 × 3 mm) and visualized the circumferential GAP43+ nerve plexus encircling the full lumen, along with its 3D relationship with multiple lymphoid follicles where *Ricinus communis* agglutinin I was used to visualize the follicular dendritic cells (**Supplementary Figure 10C**). Across an 11 mm × 6 mm × 2.5 mm pancreas block, we imaged hundreds of islets of Langerhans using anti-insulin antibodies (**Supplementary Figure 10D**). In an early stage (1a) colon cancer sample (6 mm × 9 mm × 4.5 mm), the 3D image clearly demonstrated the spatial transition from normal epithelia to the cancerous region. Cancerous cellular morphological and architectural disturbances were highlighted by 3D EpCAM immunostaining. Interestingly, we observed a vascular metastasis situated closer to the normal epithelial than to the cancerous tissue volume (104.0 µm versus 140.0 µm, respectively; **Supplementary Figure 11**). Lymphovascular metastases are grim risk factors for early-stage colon cancers that can indicate the necessity of oncological resection ^27^. The combination of EpCAM immunostaining, which highlights both epithelial cells and altered cellular arrangements, greatly facilitated detection in this sample.

### REZCUE enables highly detailed imaging of archived clinical samples

In particular, human skin tissues are difficult to clear and deep image due to their dense collagenous and complex compositions. We applied REZCUE to a human skin tissue resection FFPE block (5.3 mm × 5.3 mm × 3.3 mm) stored for 25 years. The sample was labeled as normal skin; it was obtained as a normal margin sample from a sinonasal carcinoma case. After REZCUE, we stained the tissue with two antibodies (anti-CD163, anti-CK8/18), fluorescently labeled wheat germ agglutinin (WGA) and with SYTOX Orange nuclear stain. Uniform deep penetration enabled single-cell-resolution segmentation of the 3D skin tissue structure, resulting in a complete reconstruction of all mesoscale skin structures as a microanatomical model with 2,346,568 nuclei (**Figure 3C**). From the single-cell resolution segmentation, we computed basic statistics such as cell distribution per compartment, 3D intercellular Euclidean distances, and 3D cell-to-structure distances (**Supplementary Figure 12**). The WGA histochemistry combined with nuclear staining also allowed us to restyle the image stack as brightfield 3D H&E staining through color transformation (**Supplementary Video 1**). This accurate reconstruction of the human skin structure reveals all major epidermal and dermal structures (**Figure 3D, Supplementary Figure 12, Supplementary Video 1**).

Although labeled as normal skin, we identified a metastatic lesion lodged deep within the dermal lymphatic network (**Figure 4A-B, Supplementary Video 2**), a finding missed by the original clinical analysis. Concurrent cytokeratin immunostaining supported the metastatic carcinoma nature of the lesion within the lymphatic channel (**Figure 4A**), and additional histopathology specialists unanimously confirmed the lymphovascular invasion as cancerous in a post-hoc blinded review. Lymphovascular metastasis of this kind is associated with an increased risk of distant metastasis across multiple cancer types yet is often missed because diagnosis is typically made from a single section^27,28^. The lesion in this archival sample was detected in only 3.36% of virtual sections along the Z-axis across the full 3.3 mm thickness of the block, underscoring how spatially restricted events can be missed without volumetric information.

**Figure 4.**
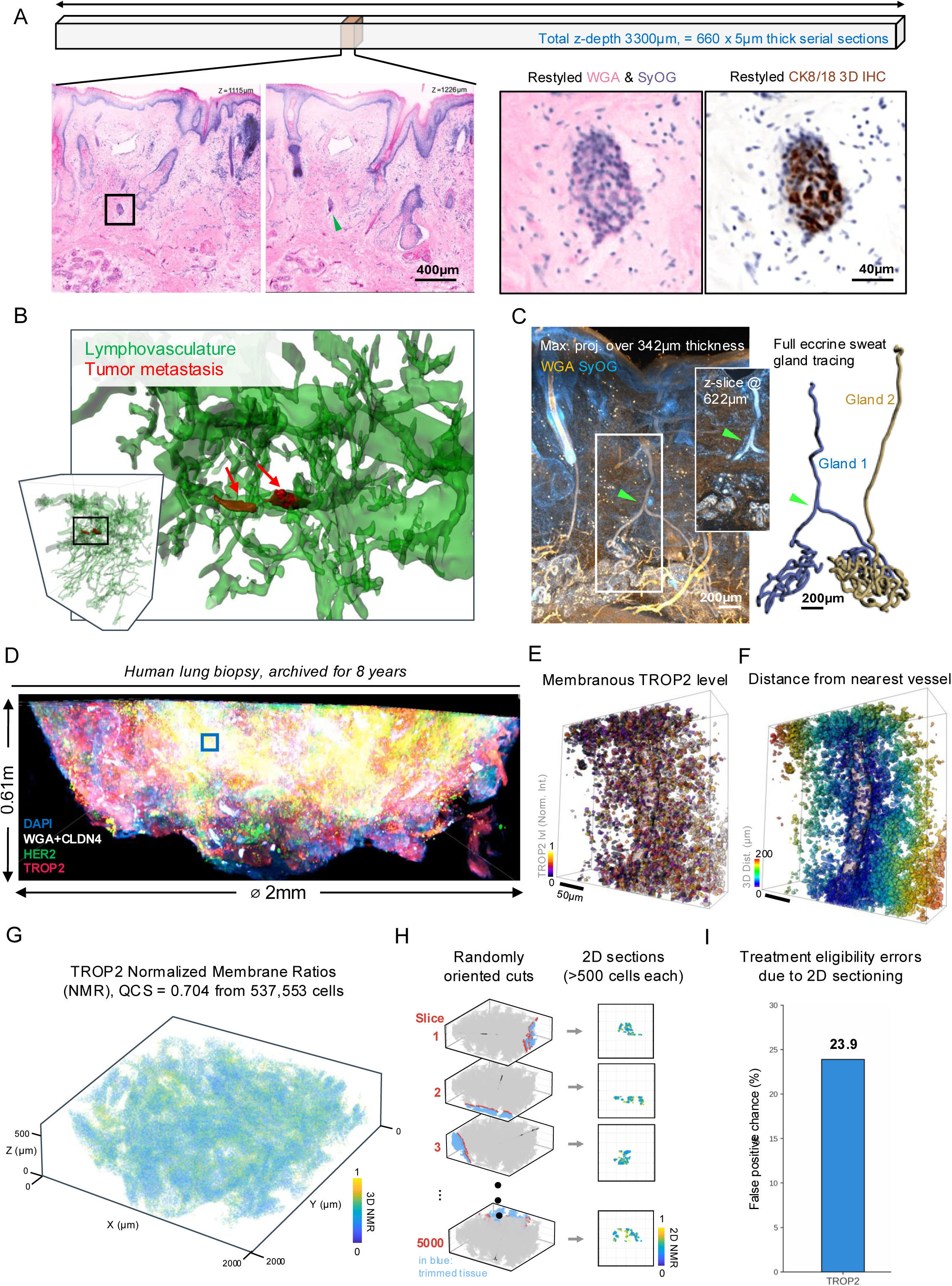
Clinical applications of REZCUE. **A.** Left: Images of the region of the human skin tissue highlighting a lymphovascular metastasis in z = 1115 µm (boxed) and z = 1226 µm (green arrowhead). This metastatic lesion was only present in sections from z = 1115-1226 µm within the 3.3 mm thickness imaged. Right: Restyled WGA and SYTOX Orange (SyOG) image and Cytokeratin 8/18 (CK8/18) immunostained image confirming epithelial origin. See also **Supplementary Video 2**. **B.** 3D rendering of the lymphovasculature throughout the human skin tissue (in green) and the segmented tumor metastasis (in red). **C.** Left: Maximum intensity projection image of region of the human skin sample with branched eccrine sweat gland duct (boxed, with branch indicated by green arrowhead). Right: Diagrams of two fully traced eccrine sweat glands from the skin sample. **D-I.** A human lung cancer biopsy was subjected to REZCUE and stained for actionable tumor markers (TROP2 and HER2) and single-cell segmentation (WGA, and CLDN4). **D)** Image of select region. Boxed area colored in dark blue. **E.** Single-cell segmentation with quantified TROP2 expression level. **F.** Distance of TROP2^+^/HER2^+^ cells from the nearest vessel. **G.** 3D scatter plot of 537,553 cancer cells color-coded by 3D TROP2 NMR. **H.** Illustration of simulated randomly oriented biopsy trimming and cutting process reminiscent of current clinical practice. Cells on the qualified section are in red; trimmed cells before obtaining the qualified section are in blue; all other cells are in grey. The en face view of the obtained section cells is plotted in 2D squares, color-coded by 2D NMR. See also **Supplementary Video 3. I.** False-positive probability of treatment-response classification. Distributions were generated from 5,000 random 2D sections simulated independently for TROP2 and HER2. The 3D ground truth for TROP2 predicted the patient to be non-responsive.

We also discovered a bifurcating eccrine sweat gland duct in this sample (**Figure 4C**). This observation could explain a decades-old anatomical discrepancy. Textbooks traditionally describe the human eccrine gland as a single coiled secretory tubule that connects to an excretory duct, which opens onto the skin surface through a sweat pore ^29^. However, the fact that fewer eccrine sweat gland skin pores than excretory ducts occur in the dermis has long puzzled anatomists. In 1968, Spearman performed dye injection into cadaveric skin pores and observed the dyes in distinct location spots under the skin, which led to the hypothesis that eccrine sweat glands bifurcate ^30^. To our knowledge, our image provides the first image-based anatomical evidence for eccrine sweat gland bifurcation in human skin.

### REZCUE reduces scoring error for patient eligibility for TROP2-targeted therapy

Next, leveraging the single-cell resolution of REZCUE-enabled 3D multiplexed immunohistochemistry, we showed how 3D quantification of morphological features can inform clinical treatment. It remains difficult to predict patient responses to antibody-drug conjugate (ADC) treatment ^31–33^. A recent analysis of the phase III TROPION-Lung01 trial demonstrated that patients with >75% of tumor cells exhibiting a normalized membrane ratio (NMR) ≤0.56 had improved responses to TROP2 ADCs ^34^. NMR is defined as the ratio of TROP2 membrane staining intensity to the combined membrane and cytoplasmic staining intensity. The fraction of tumor cells with an NMR ≤0.56 is referred to as quantitative continuous scoring (QCS). This score was subsequently used to determine patient eligibility for TROP2-targeted therapy ^34^.

The QCS score in the phase III TROPION-Lung01 trial was calculated from a single tissue section per patient. We therefore sought to investigate the extent to which random sectioning bias could affect QCS estimates. To develop a 3D analogue of QCS, we performed REZCUE on a TROP2^+^/HER2^+^ human FFPE lung cancer biopsy sample (2 mm × 2 mm × 0.6 mm), which had been archived for 8 years. We stained the sample with WGA and three antibodies (anti-CLDN4, anti-TROP2, and anti-HER2) (**Figure 4D**). WGA and CLDN4 were used for single-cell segmentation and TROP2 and HER2 were used to identify ADC targets. A UNet-based segmentation model was used for cancer area detection, and single-cell segmentation was performed with Cellpose-SAM ^35^ (**Method Figure 1**). NMRs and distances to nearest blood vessels were also computed for all the cancer cells (537,553 cells) from the sample (**Figure 4E-G**).

To quantify the discrepancy between 3D and 2D QCS on the same sample, we performed in silico sectioning of the 3D tissue volume. Sectioning was performed 5,000 times at different orientations, with each resulting section required to contain at least 500 cells to qualify for analysis (**Figure 4H, Supplementary Figure 13A, Supplementary Video 3**). The 3D QCS value was 0.704, suggesting the patient is ineligible for TROP2 ADC treatment, but 23.9% of the 2D sections had QCS values that would have falsely suggested treatment eligibility (**Figure 4I, Supplementary Figure 13B**). The intratumoral heterogeneity in TROP2 NMR may explain the lack of statistical significance reported in a TROP2 QCS study of a small cohort ^36^. Assuming 3D QCS represents the ground truth as compared to the 2D QCS, by removing the sampling bias that tumor heterogeneity introduces into single-section assessment, 3D QCS reduced false-positive calls by approximately 25% relative to conventional 2D scoring. This approach could reduce the need for repeat biopsies, additional tissue consumption, and the treatment delays that follow from an inaccurate initial biomarker call. Importantly, REZCUE supports robust 3D NMR-based quantification of phenotypic biomarkers while preserving the same tissue block for subsequent genotype-based profiling to guide targeted therapy selection.

### REZCUE-treated specimens are compatible with downstream multi-omics molecular profiling

Although 3D multiplexed immunohistochemistry volumetric imaging provides critical information on tissue architecture and spatial organization, downstream orthogonal assays are equally important for characterizing the molecular state of individual cells. To show that REZCUE does not interfere with multimodal molecular profiling, we sagittally bisected a wild-type mouse brain and subjected both hemispheres to FFPE processing. One hemisphere was retained as an untreated control, whereas the other underwent REZCUE processing, immunostaining with anti-NeuN (neurons), anti-AQP4 (astrocytes), and anti-CNP1 (oligodendrocytes), 3D imaging, and re-embedding in paraffin wax (**Figure 5A-C**). Both hemispheres were then sectioned in parallel and subjected to a matched panel of downstream multimodal assays, including spatial transcriptomics using the CosMx Spatial Molecular Imager platform, bulk RNA sequencing, and proteomics by mass spectrometry. REZCUE-processed and control samples had comparable transcript counts (**Figure 5D-E**). To evaluate concordance at the cellular level, we performed standard preprocessing of the CosMx data and annotated cell types using established cell type-specific markers (**Supplementary Figure 14A-B**). The REZCUE-processed and control samples had similar cellular compositions with comparable proportions of the major neuronal cell types (**Figure 5F-H**). The largest differences in cell-type composition were observed for oligodendrocytes and glutamatergic neurons with ∼10% and ∼6% higher abundances, respectively, in the REZCUE sample than the control (**Figure 5H**). These differences may reflect intrinsic variability rather than effects of REZCUE processing. We next mapped the annotated cell types back to their spatial coordinates to assess whether the underlying tissue architecture was preserved after REZCUE treatment of the tissue. The spatial distributions of neuronal cell types were highly consistent with the anatomical organization of the control brain (**Figure 5I**), and differences in cell-type fractions were minimal (**Figure 5J**), indicating that REZCUE processing preserves transcript molecular integrity, cellular composition and spatial organization and is compatible with downstream spatial transcriptomic profiling.

**Figure 5.**
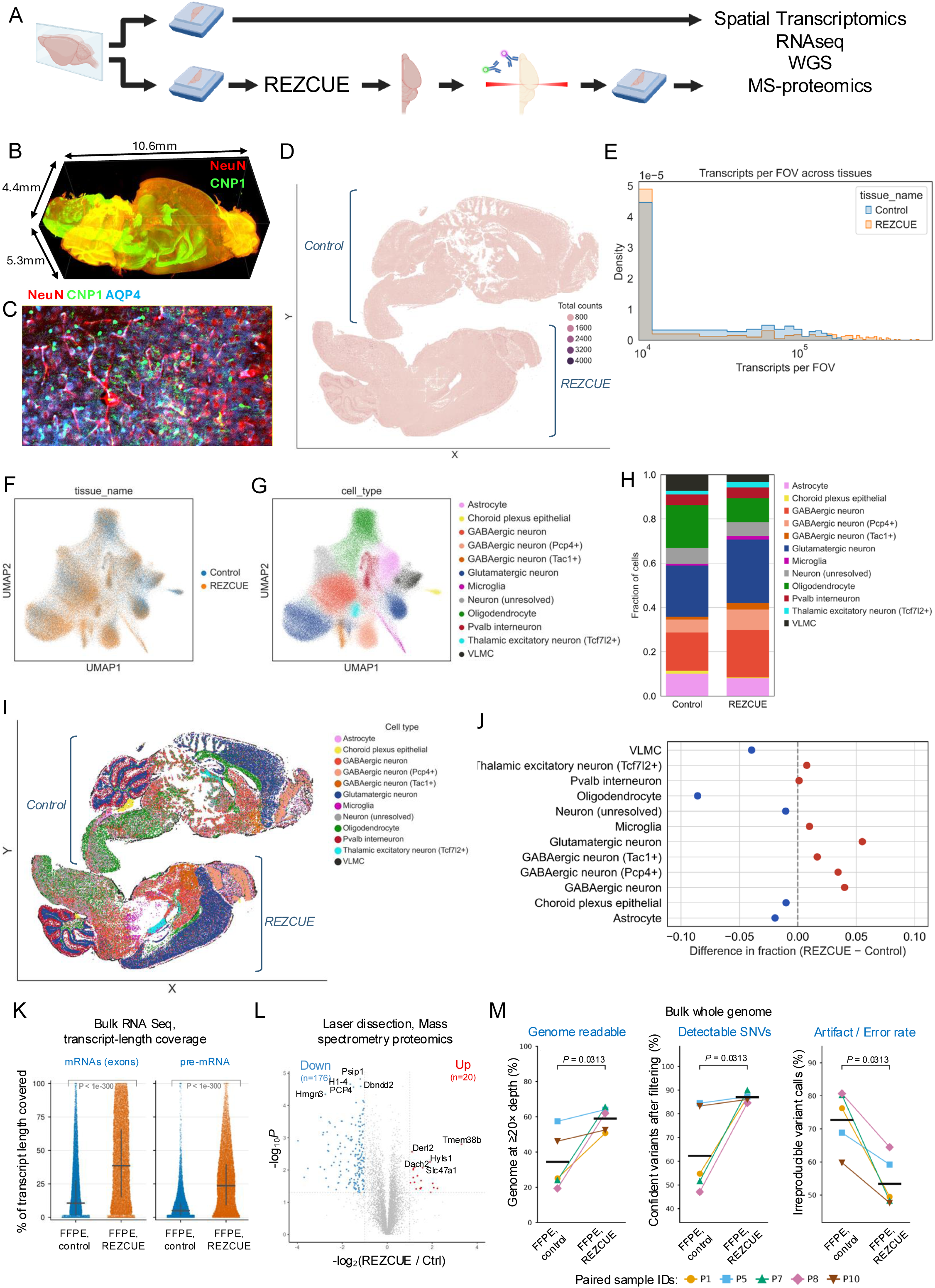
REZCUE preserves molecular and cellular fidelity. **A.** Schematic of the experimental workflow comparing standard processing of FFPE tissue (top) with the REZCUE workflow (bottom). REZCUE-treated tissue was de-crosslinked, re-embedded, and re-sectioned prior to downstream spatial transcriptomics, bulk RNA sequencing, whole-genome sequencing, and mass spectrometry-based proteomics. **B.** 3D LSFM reconstruction of a REZCUE-processed mouse brain hemisphere immunolabeled for NeuN (neurons, red) and CNP1 (oligodendrocytes, green). **C.** High-magnification image of REZCUE-processed tissue immunolabeled for NeuN (red), CNP1 (green), and AQP4 (astrocytes, blue). **D.** Spatial transcriptomic sections from matched control (top) and REZCUE-processed (bottom) mouse brain hemispheres, colored by total transcript counts per field of view (FOV). **E.** Density distribution of transcripts detected per FOV in control (blue) and REZCUE-processed (orange) tissues. **F.** UMAP embedding of single-cell spatial transcriptomic profiles from control and REZCUE-processed tissue colored by tissue of origin. **G.** UMAP embedding colored by annotated cell type. **H.** Plots of cell-type compositions (fractions of cells) in control versus REZCUE-processed tissues. **I.** Spatial sections from control and REZCUE-processed brains colored by cell type. **J.** Difference in cell-type fraction (REZCUE minus control) for each annotated cell type; dashed line indicates no difference between conditions. **K.** BulkRNA sequencing quality metrics for FFPE control and REZCUE-treated samples. The *p* values from paired Wilcoxon signed-rank tests are indicated. **L.** Volcano plot of protein abundances in REZCUE-treated versus control tissue; significantly up (red) and downregulated (blue) proteins are indicated. **M.** Volcano plot of bulk RNA-seq differential expression (REZCUE versus control); significantly up (magenta) and downregulated (green) genes are indicated relative to fold-change and significance thresholds (dashed lines).

We also performed bulk RNA-sequencing of the REZCUE-processed mouse hemibrain and the contralateral control (**Supplementary Table 2**). This sequence information complements the in-situ hybridization-based RNA identification of CosMx. We found RNA yield and library qualities were lower from the control than from the REZCUE-processed sample: For 18 out of 20 predefined QC metrics, the REZCUE-processed sample equaled or exceeded that of control (**Supplementary Figure 15A**) and sequencing accuracy was similar (**Supplementary Figure 15B**). REZCUE provided broader coverage across the full transcript lengths for both mRNAs and pre-mRNAs (**Figure 5K**), with 93.1% of mRNAs and 96.0% pre-mRNAs having improved transcript coverage compared to the control. Based on intron-to-exon read depth per gene, we ruled out the possibility of DNA contamination in both REZCUE-processed and control samples (**Supplementary Figure 15C**). REZCUE processing resulted in higher per-gene transcript count (**Supplementary Figure 15D**), and the control dataset was dominated by a structural RNA artifact with 46.1% of mapped reads concentrated in two signal recognition particle loci versus 4.0% in REZCUE. Compared with the control, REZCUE had more effective gene detection (12252 vs 9292 genes at ≥5 reads). Known brain transcripts (e.g., *Mbp*, *Snhg14*, *Cdr1*) were among the top 60 most abundant expressed genes in the REZCUE-processed sample, whereas the most abundant transcripts in the control sample were non-coding RNAs. Consistent with this, most of the mapped reads in the control correspond to structural RNAs, whereas in the REZCUE-processed sample, most reads were from mRNAs (**Supplementary Figure 15E**). The superchaotropic boron cluster reagent used in the REZCUE process may help preserve library complexity due to its broad nuclease inhibitory effects ^37^, although this cannot be tested statistically with the data generated.

To test if REZCUE is also compatible with proteomics by mass spectrometry (MS), we compared three sections from REZCUE-treated samples against three control sections. We laser-microdissected anatomically equivalent areas (the mouse cerebellum) from these sections and extracted total proteins for MS. Of the proteins detected in both REZCUE-processed and control samples, only 2.4% were significantly reduced in abundance in REZCUE-treated samples (**Figure 5L**). However, there was a slight but statistically significant decrease in total protein counts in the REZCUE-processed sections versus the control on average (7335.6 vs. 7402.3 proteins, *p* = 0.039, unpaired two-sided *t*-test; **Supplementary Figure 16A**). Nonetheless, pairwise comparisons between individual REZCUE-treated samples and control samples showed high levels of concordance of at least 0.96 between any given samples (**Supplementary Figure 16B-D**). Complementing MS-based proteomics, we also show that REZCUE treatment does not affect downstream section-based chromogenic immunohistochemistry (**Supplementary Figure 17**). We thus conclude that REZCUE processing preserves proteins relatively well, enabling highly accurate spatial proteomics analysis of the tissue.

Finally, we tested whether REZCUE-treated samples are compatible with downstream whole genome sequencing (**Figure 5M**). We obtained 10 FFPE human lung cancer blocks archived for 5-10 years. To ensure that the comparisons were based on materials of the same genetic composition, we homogenized the FFPE tissues and divided each sample into portions of equal weight to form a matched pair. Although DNA was substantially fragmented, as expected for FFPE tissue, in matched comparisons, similar amounts of DNA were extracted from pairs of REZCUE-processed and control samples (4.528 μg vs. 5.439 μg; exact one-sided Wilcoxon signed-rank test, *p* = 0.024 against a non-inferiority margin of 20% from control) and comparable whole-genome sequencing quality metrics were obtained (**Supplementary Figure 18A–B, Supplementary Tables 3–4**). Five matched pairs passed quality control and proceeded to sequencing. When an identical variant-calling filter was applied to REZCUE-processed and control samples within a common sequencing-depth window, REZCUE-treated samples had a higher fraction of reproducible variants than paired controls (86.95 ± 1.96% vs. 64.21 ± 18.16%; favorable in 5 of 5 pairs; exact one-sided Wilcoxon signed-rank *p* = 0.031 against a 10% non-inferiority margin) (**Supplementary Figure 18C**), and a larger fraction of the genome was callable at ≥20× depth (by 24.6%; favorable in 5 of 5 pairs; exact one-sided Wilcoxon signed-rank test, *p* = 0.031 against a −5% non-inferiority margin) in the REZCUE-processed samples (**Figure 5M**). Despite detecting more reproducible variants, REZCUE did not increase the rate of irreproducible calls: The rate was lower than in controls (53.81 ± 7.59% vs. 73.15 ± 8.86%; one-sided Wilcoxon signed-rank *p* = 0.031) (**Figure 5M, Supplementary Figure 18C**). Together, these results indicate that the approximately 29% reduction in total single-nucleotide variant (SNV) calls after REZCUE treatment reflects the removal of artifactual calls rather than a loss of true variant detection (**Supplementary Figure 18C**). Overall, REZCUE achieved SNV and mutation recovery from archival FFPE tissue comparable to that of paired standard-processing controls. Thus, REZCUE-processed tissue is compatible with downstream spatial transcriptomics, RNA sequencing, spatial mass-spectrometry-based proteomics, and whole genome sequencing.

### REZCUE enables volumetrically precise, single-cell level, multi-omic analysis of unique tumor compartments

To demonstrate the power of integrating 3D volumetric imaging with downstream multimodal molecular profiling on the same FFPE tissue block, we applied REZCUE to a human breast cancer FFPE tissue specimen (25 mm × 12 mm × 4 mm) that had been archived for 25 years. Because breast tumors are molecularly heterogeneous, resolving clonal subtypes is critical for treatment decisions ^38^. 3D imaging can map the spatial architecture of distinct phenotypic compartments across a tumor, but mapping that architecture is only a first step: Interpreting its biological and clinical significance requires molecular characterization of the spatially defined populations. By allowing 3D imaging to be combined with downstream molecular assays on the same tissue, the REZCUE methodology enables linking each compartment’s 3D organization to its molecular state and microenvironment for a multiscale view of tumor heterogeneity.

We first treated the tissue block with REZCUE, then immunostained it with a three-antibody panel (anti-TROP2, anti-HER2, anti-HER3) plus a nuclear marker and imaged it with LSFM (**Figure 6A**). Following 3D imaging, the tissue was re-embedded and processed for CosMx spatial transcriptomics, enabling molecular characterization at single-cell resolution. We then selected three spatially adjacent regions for proteomics, two spatially adjacent regions for miRNA sequencing, and subsequently punch biopsies for whole exome sequencing analyses, thereby enabling integrated analysis of tissue architecture, transcriptomic state, proteomic composition, and genomic variation within the same archived specimen (**Figure 6A**). Subsequently, we segmented individual tumor compartments (**Figure 6B**) and identified six spatially discontiguous compartments (**Supplementary Figure 19**). Since the six tumor compartments were not present in the same plane, a plane containing three distinct tumor compartments was used for the CosMx spatial transcriptomics experiment.

**Figure 6.**
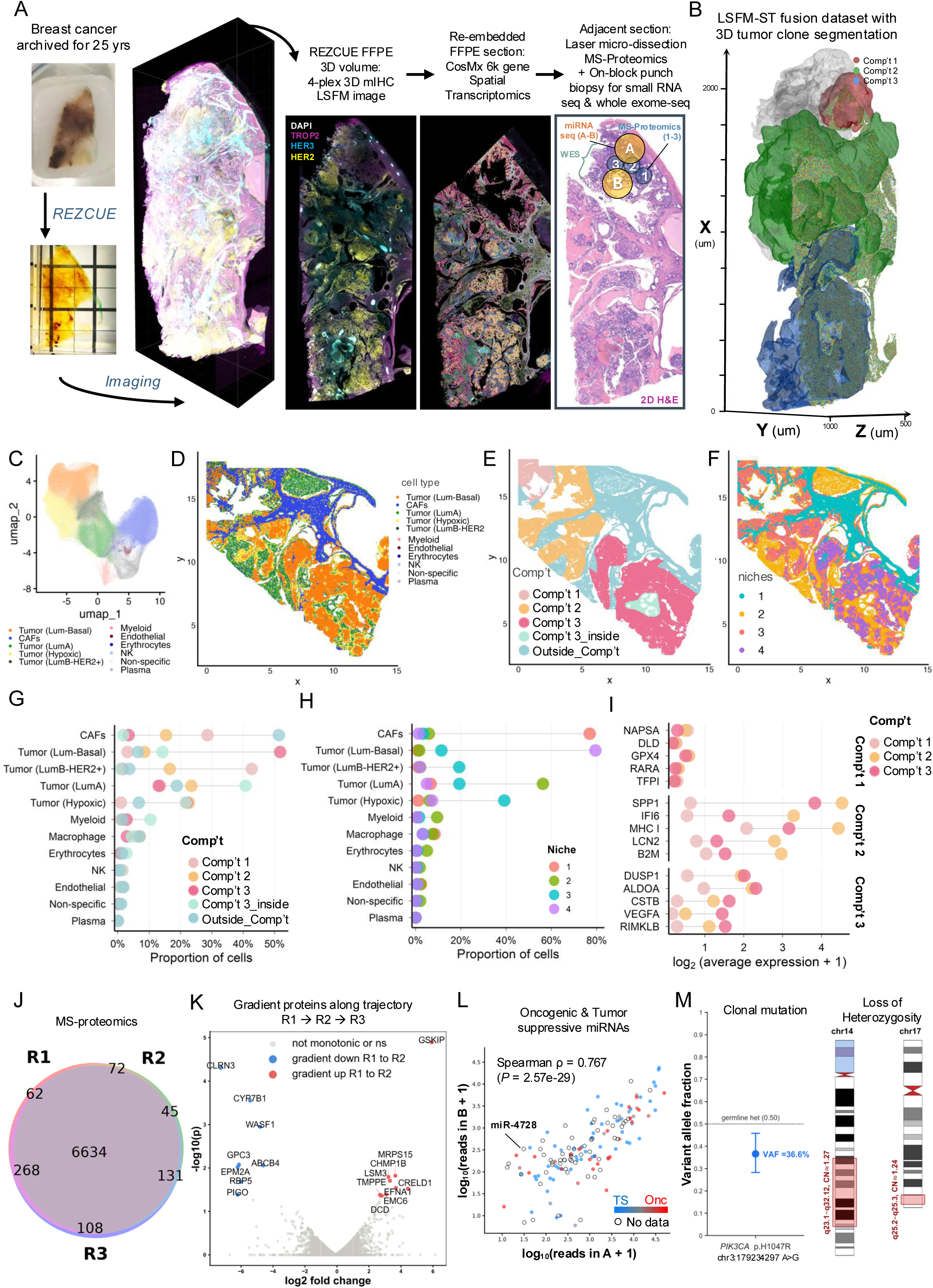
Spatial multi-omic profiling of breast tumor tissue processed with REZCUE reveals a molecularly distinct, spatially buried tumor compartment. **A**. Workflow for 3D spatial multi-omic profiling of a breast tumor archived as FFPE for 25 years. The tissue block was processed with REZCUE and imaged by four-plex 3D immunohistochemistry LSFM to generate a whole-block volumetric reconstruction, then re-embedded and sectioned for CosMx 6,000-gene spatial transcriptomics. An adjacent section was used for laser microdissection-coupled mass spectrometry proteomics and on-block punch biopsy for small RNA sequencing, and was subsequently extracted for whole-genome sequencing. Inset: H&E-stained section showing on-slide laser microdissected regions (1, 2, and 3) analyzed for protein composition and punch-biopsy sites analyzed for miRNAs (A and B) within the compartment 1 region. See **Supplementary Video 4**. **B.** 3D reconstruction of the LSFM and spatial transcriptomics merged the dataset with tumor compartments (compartment 1, red; compartment 2, green; compartment 3, blue) overlaid on the whole tumor volume (gray). **C.** UMAP of single-cell spatial transcriptomic profiles colored by annotated cell type, including four molecularly distinct tumor populations (Luminal-Basal, Luminal A, Luminal B-HER2^+^, and Hypoxic) and stromal and immune cell types. **D-F.** Spatial maps of a representative tissue section colored by **D.** cell type, **E.** 3D-defined tumor compartment assignment. A distinct enclosed cavity within compartment 3 was identified in the 3D reconstruction and designated as *compartment 3_inside*. The remaining tissue regions not belonging to compartments 1, 2, or 3 were collectively designated as *Outside_compartments*. **F.** colored by niche assignment as defined by local cell-type composition. **G.** Proportion of each cell type within each 3D-defined tumor compartment. **H.** Proportion of each cell type within each spatial niche. **I.** Average expression of indicated genes in compartments 1, 2, and 3. **J.** Venn diagram of proteins detected across three regions sampled within compartment 1. **K.** Volcano plot of protein abundance along the region 1→2→3 trajectory within compartment 1. Proteins with significant, monotonically decreasing (blue) or increasing (red) abundance across the trajectory are indicated; non-monotonic or non-significant proteins are shown in gray. Representative gradient proteins are labeled. **L.** Scattered plot of miRNAs detected from punch biopsy sites A and B in compartment 1. Oncogenic and tumor-suppressive miRNAs are colored red and blue, respectively, whereas miRNAs with non-polarized or conflicting roles in breast cancer are colored grey. Those with no literature experimental data were plotted as open circles. **M.** Whole-genome sequencing profiling of the remaining biopsy on compartment 1 revealed the *PI3KCA* p.H1047R mutation and loss of heterozygosity in chr14q23.1-q31.12. VAF, variant allele frequency. Based on 3D LSFM imaging, tumor cells account for 70-80% of the sample.

We identified 441,383 cells in the CosMx data. We developed a custom 2D-to-3D registration pipeline to nonlinearly align the CosMx section to the LSFM 3D image volume (**Method Figure 2, Supplementary Video 4**). Using established cell-type markers, we identified four molecularly distinct tumor subtypes and five immune and stromal cell populations within the CosMx section (**Figure 6C**; **Supplementary Figure 20A–B**). The spatial distributions of these annotated cell types were consistent with previously described breast tumor architecture (**Figure 6D**). By alignment of the 3D tumor compartment masks derived from volumetric imaging with the corresponding CosMx spatial transcriptomic data, we were able to perform molecular and cellular characterization of individual tumor compartments within their 3D-defined boundaries (**Figure 6E**). This integration revealed information that is not captured by conventional local niche analysis, in which cells are grouped according to their immediate microenvironment ^39^. Whereas niche analysis provides a localized description of cellular neighborhoods (**Figure 6F; Supplementary Figure 20C**), the 3D compartment boundaries provide a higher-order, volumetric representation of tumor organization, allowing tumor compartments to be assessed in the context of their true 3D continuity and partitioning. By comparing cell-type compositions and molecular signatures across 3D-defined tumor compartments and their corresponding local niches (**Figure 6G–H**), we determined that compartment 1 is distinct from other identified compartments. Compartment 1 tumor cells have elevated levels of HER2, CXCL14, OXR1 (a key regulator of oxidative-stress protection), and LEFTY1 (a Nodal/TGF-β antagonist associated with developmental and luminal differentiation programs) (**Figure 6I**). Reactivation of LEFTY1 in tumor cells is associated with more differentiated, hormone-responsive states rather than aggressive phenotypes ^40^. These features are indicative of a distinct Luminal B-HER2^+^ identity of compartment 1 relative to the other tumor compartments. Collectively, the molecular profiles suggest distinct functional states across compartments: Compartment 1 appears to be metabolically and oxidatively buffered, compartment 2 has a more interferon-active and immune-engaging phenotype, whereas compartment 3 has the strongest invasive and metabolic signature.

To further understand the tumor heterogeneity, we collected three adjacent regions within the compartment 1 area for proteomic analysis (annotated 1, 2, and 3 in **Figure 6A**) as we observed prominent immune cell aggregates around the compartment on the H&E-stained section. We first assessed protein detection overlap across the three regions and found high overall proteomic similarity, with 6,691 of 7,391 total proteins (∼91%) detected in all three regions (**Figure 6J, Supplementary Figure 20D**). Despite this overall similarity, trajectory analysis identified a discrete subset of proteins that increased monotonically in abundance from region to region, including SAAL1, SOAT2, CYP3A7, and GZMK (**Figure 6K**). This suggests that there is a continuous gradient in inflammatory, hormone-metabolic, and immune-associated programs across the tumor consistent with positioning of the first region closer to a stromal/immune-rich interface and the third closer to a more tumor cell-rich microenvironment.

To further demonstrate the compatibility of REZCUE processing with sequencing-based assays, we obtained two punch biopsies from the compartment 1 region (annotated A and B in **Figure 6A**) for small RNA sequencing. Small RNA species such as miRNAs are poorly recovered from archival samples, particularly those stored for decades ^41^. Under stringent criteria, we identified 145 miRNAs: 29 were classified as oncogenic, 56 as tumor-suppressive, and 60 as ambiguous (**Supplementary Table 5, Figure 6L**) ^42,43^. The abundances of these miRNAs were highly concordant in the two regions sampled (Spearman *ρ* = 0.767, *p* = 2.57 × 10^−29^), indicative of reliable detection. The *ERBB2*-intronic miR-4728-3p was detected in both regions, consistent with HER2 expression in this tumor sample.

We next characterized genomic mutations by whole exome sequencing on a final punch biopsy on compartment 1 (**Supplementary Figure 21A–B**). Whole exome sequencing revealed a clonal, non-germline mutation in *PIK3CA* that resulted in an H1047R substitution in this protein (variant allele fraction = 0.366; with 70-80% cells in the sample as tumor cells by histology; **Figure 6M**). This is a well-known driver mutation in breast cancer that is targetable by PI3K inhibitors such as alpelisib ^44^. *TP53*, *PTEN*, *AKT1*, *ESR1*, *ERBB2*, *BRAF*, and *ERBB2* were negative for mutations (**Supplementary Figure 21C**), and none of these loci showed focal gene amplification (**Supplementary Figure 21D**). The sample also showed a slight dominant subclonal loss of heterozygosity at 17q25.2– q25.3 (folded B-allele frequency (BAF) = 0.242 [95% CI 0.224–0.250]; log2 ratio = −0.484; family-wise error rate *p* value (*P*_FWER_) = 0.0559); this region is frequently altered in breast cancer ^45,46^. There was also an additional loss of heterozygosity at 14q23.1–q32.12 (folded BAF = 0.233 [95% CI 0.200–0.263]; log2 ratio = −0.464; *P*_FWER_ = 0.0719) (**Figure 6M, Supplementary Figure 21E**), another region known to be altered in breast cancer ^47^. Although HER2 was detected by immunohistochemistry in this sample, the gene encoding HER2, *ERBB2* was not amplified (upper-bound copy number estimate = 3.0) (**Supplementary Figure 21D**). HER2 upregulation can occur independently of ERBB2 amplification ^48^.

Together, these results show that integrating 3D imaging with spatial transcriptomics, proteomics, and sequencing-based assays on the same archival tissue block resolves both the spatial organization of individual tumor compartments and their underlying molecular and genomic identity. In this specimen, that integration linked the 3D architecture of compartment 1 to a driver mutation in *PIK3CA*, subclonal loss of heterozygosity at 17q25.2–q25.3 and 14q23.1–q32.12, and a HER2-positive phenotype occurring independently of *ERBB2* amplification. REZCUE processing also enabled analyses of small RNA species such as miRNAs that are typically difficult to obtain from decades-old FFPE tissue. Although individual measurements, such as the *PIK3CA H1047R* mutation, could be obtained by sequencing alone, assigning these molecular and genomic features to a specific, spatially buried tumor compartment required that compartment to be defined in 3D and then sampled from the same preserved tissue block. Preserving one tissue volume for multiple spatially registered assays thus allowed us to link 3D structural context to the transcriptomic, proteomic and genomic state of individual tumor compartments, which would be difficult to achieve from imaging or molecular profiling alone, or from a single arbitrarily oriented 2D section.

## Discussion

The development of REZCUE enables a framework for transforming long-term archived human tissue into an accessible resource for integrated 3D and cross-modal molecular analysis. Critically, by redesigning antigen-retrieval chemical approaches, REZCUE removes the trade-off between 3D imaging and molecular profiling, allowing antigen retrieval without degrading biomolecules. We discovered methylcyclopentane/cyclohexane allows highly efficient tissue deparaffinization, designed a transitional solvent blend to avoid wax reprecipitation and developed an aqueous cocktail to expose epitopes. Using this method, we showed that multi-millimeter-thick specimens archived for up to 30 years can be 3D-immunostained with deep, uniform probe penetration, without losing the molecular information required for downstream spatial genomic, transcriptomic, and proteomic analyses. Such dual compatibility sets it apart from other 3D immunostaining techniques ^9,10,14,15,17^ and stems from the careful design choice of REZCUE chemistry around the physicochemical properties of tissue biomolecules. Unlike other post-tissue clearing, single modality omics profiling methods, which have relied on small molecule dyes or a single anti-GFP nanobody for staining, REZCUE supports 3D immunostaining with a panel of 160 commercially available, whole-IgG antibodies, while preserving the specimen for multi-omics profiling ^18–20^. The performances of DNA and RNA assays were slightly better than controls. This was not unexpected given a recent finding that the boron cluster [B_12_H_12_]^2-^ utilized in the REZCUE workflow possessed broad nuclease inhibitory activity ^37^. Together, these results establish a framework for maximizing archival tissue utility and future-proofing clinical trial specimens. By staining and imaging REZCUE-processed samples first by 3D LSFM and subsequently by 2D spatial multi-omics, researchers can now link macroscopic tissue architecture with cellular and molecular states, effectively generating a digital preview of an entire specimen before triaging them for fine-scale, high-resolution spatial multi-omics profiling.

We acknowledge several limitations in our study. As a methodological development study, we demonstrated the utility of 3D tissue imaging compared with conventional 2D imaging in a small number of samples with replicates, but future studies implementing REZCUE at scale will be required to assess the biological and clinical impact of 3D tissue imaging. Meanwhile, although REZCUE removed the chemical barrier to combining 3D imaging with molecular profiling, several challenges remain — most reflecting the limits of current imaging and computational technologies rather than sample processing issues. First, the number of markers that can be resolved simultaneously in 3D (4-plex in this study) remains far below the multiplexing achievable in 2D. For example, spatial transcriptomics methods effectively enable genome-wide coverage ^49^. This gap stems from constraints in volumetric fluorescence imaging, including limited channel numbers, spectral overlap, and cyclic-staining throughput, rather than a ceiling imposed by the retrieval chemistry. Advances in 3D-compatible cyclic staining technologies and imaging instrumentation will be necessary to increase the number of markers that can be imaged in 3D, and REZCUE’s non-destructive tissue preparation will support this ^50,51^. Second, once a 3D image is acquired, choice of the plane to carry forward for 2D spatial multi-omics is a computational problem that will be solved by new image-registration and sampling strategies rather than new chemistry ^11,52,53^. Finally, methods for spatially resolved imaging of the whole exome and proteome on 2D sections have not yet been developed. We anticipate that REZCUE will provide the chemistry backbone for true 3D spatially resolved multi-omics analyses.

---

**Supplementary Figure 1.**
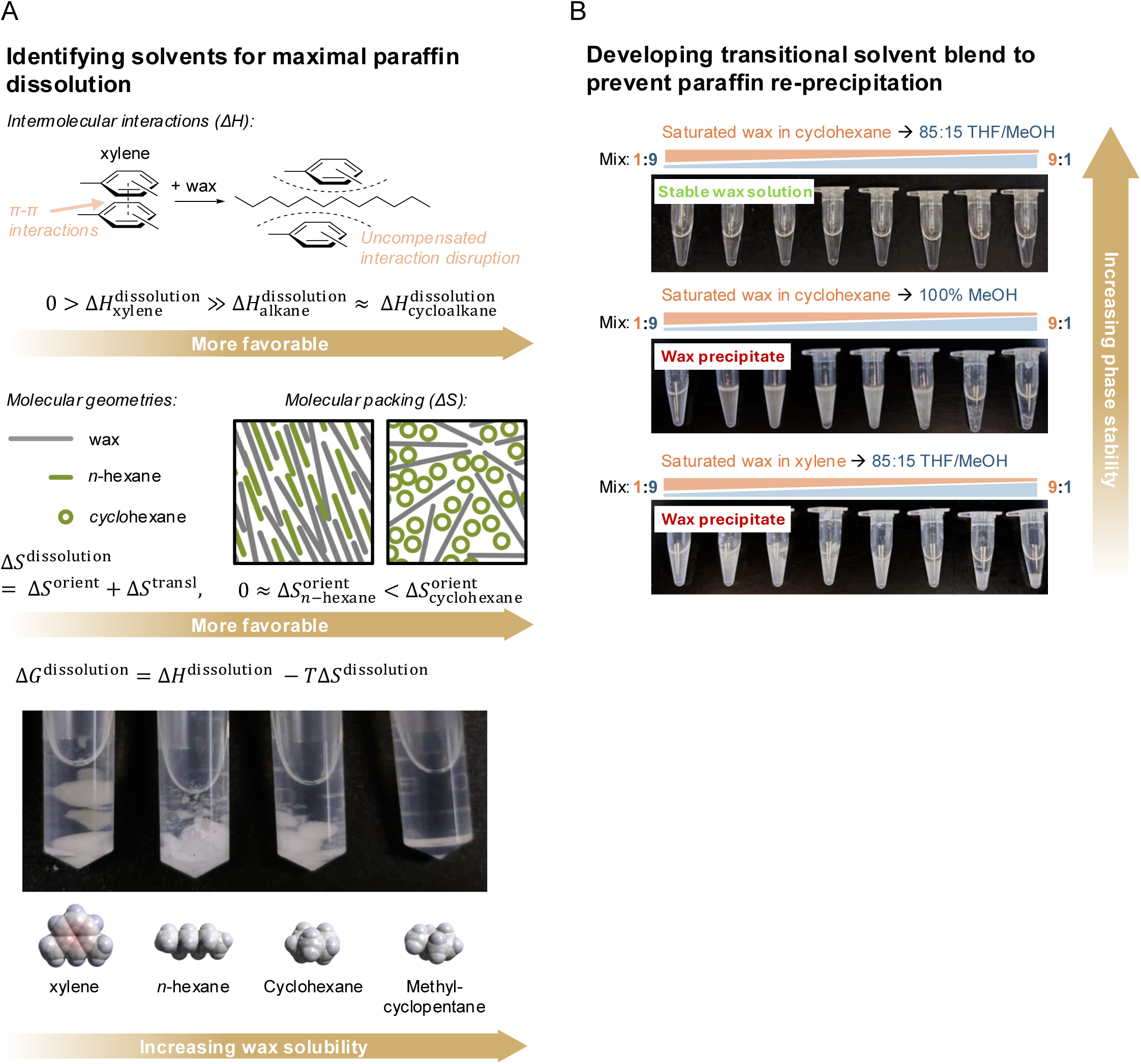
REZCUE chemistry overview. **A**. Solvent choice rationale for paraffin wax dissolution. Upper panel: The intercalation of aliphatic paraffin wax molecules between xylene molecules during deparaffinization disrupts their π-π interactions, rendering dissolution enthalpically unfavourable. Middle panel: Switching to *n*-hexane, which is also linear, promotes aligned molecular arrangements to maximize translational entropy (Δ*S*^transl^) but sacrificing orientational entropy (Δ*S*^orient^) for the long-chained paraffin wax molecules. This does not happen with spheroid-shaped cyclohexane which further gains orientational entropy in paraffin wax dissolution. Lower panel: Experimental comparison by dissolving 1g paraffin wax in 500 µl organic solvent at 20-25 °C for 30 minutes. Methylcyclopentane provided the fastest wax dissolution. **B**. Developing transitioning solvent washes after tissue deparaffinization by cyclohexane and before moving to anhydrous alcohol washes. Wax-saturated solvent (cyclohexane or xylene) was mixed with two solvent blends (85:15 v/v THF/MeOH or 100% MeOH) at 1:9 to 9:1 v/v ratios and photographed after 30 minutes to note for any paraffin wax re-precipitation. The cyclohexane-to-THF/MeOH transition did not lead to wax precipitation, while both cyclohexane-to-MeOH, or xylene-to-THF/MeOH led to wax re-precipitation. This suggests that wax solubility is highly dependent on the resulting ternary solvent system composition.

**Supplementary Figure 2.**
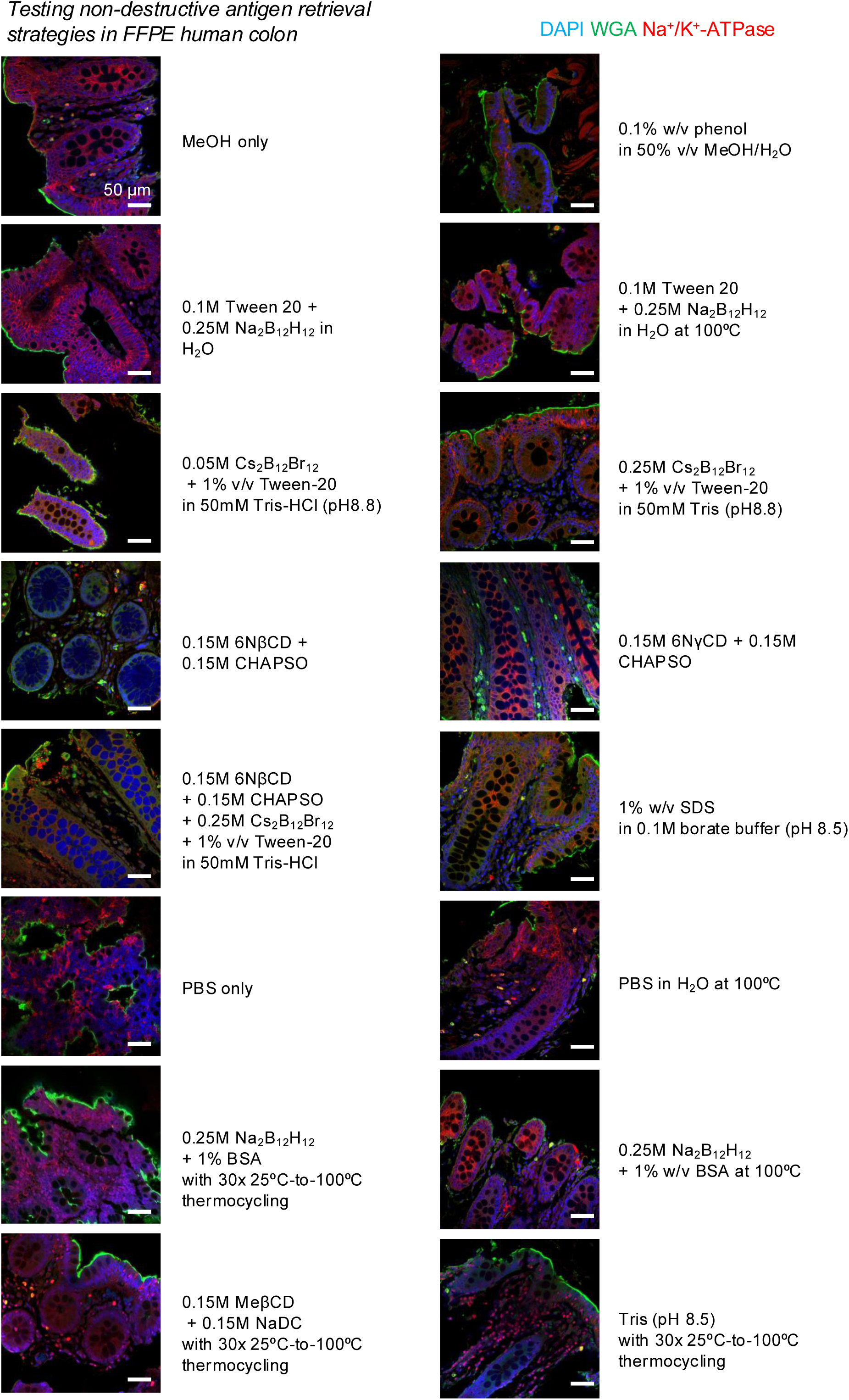
Exploration of aqueous-based antigen retrieval strategies. FFPE human colon tissues were deparaffinized using our REZCUE method and were then incubated at 55 °C (unless otherwise specified) for 24 hours in various aqueous chemical cocktails, as labelled in the figure. The tissue was then stained with DAPI, WGA and anti-Na^+^/K^+^-ATPase antibodies using the INSIHGT2 protocol and imaged at 100-120 µm depths using a confocal microscope. These results established an aqueous solution consisting of the superchaotrope [B_12_H_12_]^2-^ and a detergent to be the most promising for volumetric antigen retrieval.

**Supplementary Figure 3.**
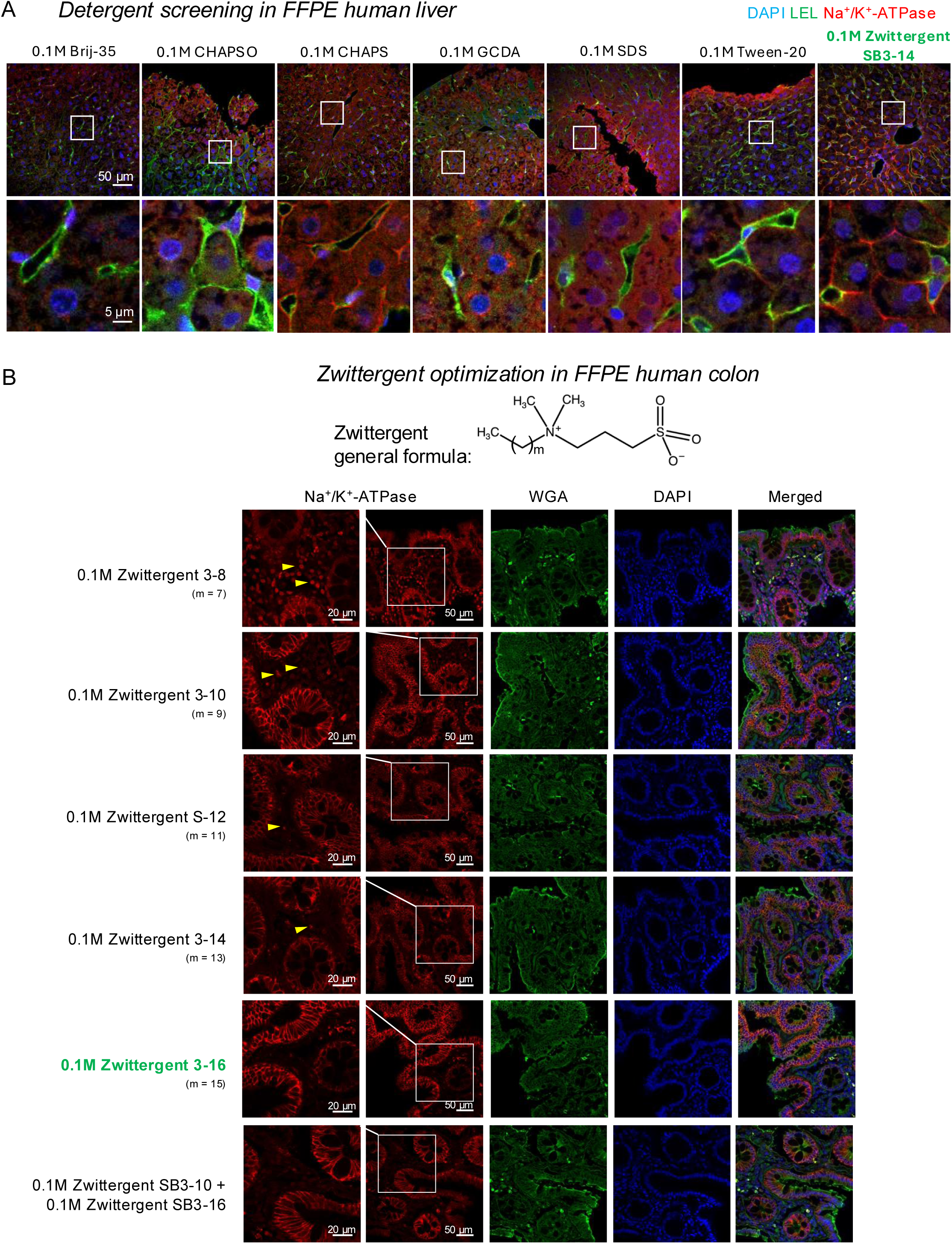
Detergent screening and zwittergent structure optimization. **A**. Detergent screening in FFPE human liver tissues, which were deparaffinized using our REZCUE method, incubated in aqueous solutions containing 0.25 M Na_2_B_12_H_12_ and 0.1 M indicated detergents at 55 °C, and stained using the INSIHGT2 protocol for DAPI, LEL (tomato lectin) and anti-Na^+^/K^+^-ATPase antibodies, which highlights hepatocyte membranes. **B**. Structural optimization of aliphatic zwittergent, for which the general formula is shown, with screening experiment performed as in **A**. The longer the aliphatic chain length, the less the non-specific nuclear staining in the resulting images (indicated by yellow triangles). Such structural optimization did not compromise the staining quality of lectins and nuclear dyes.

**Supplementary Figure 4.**
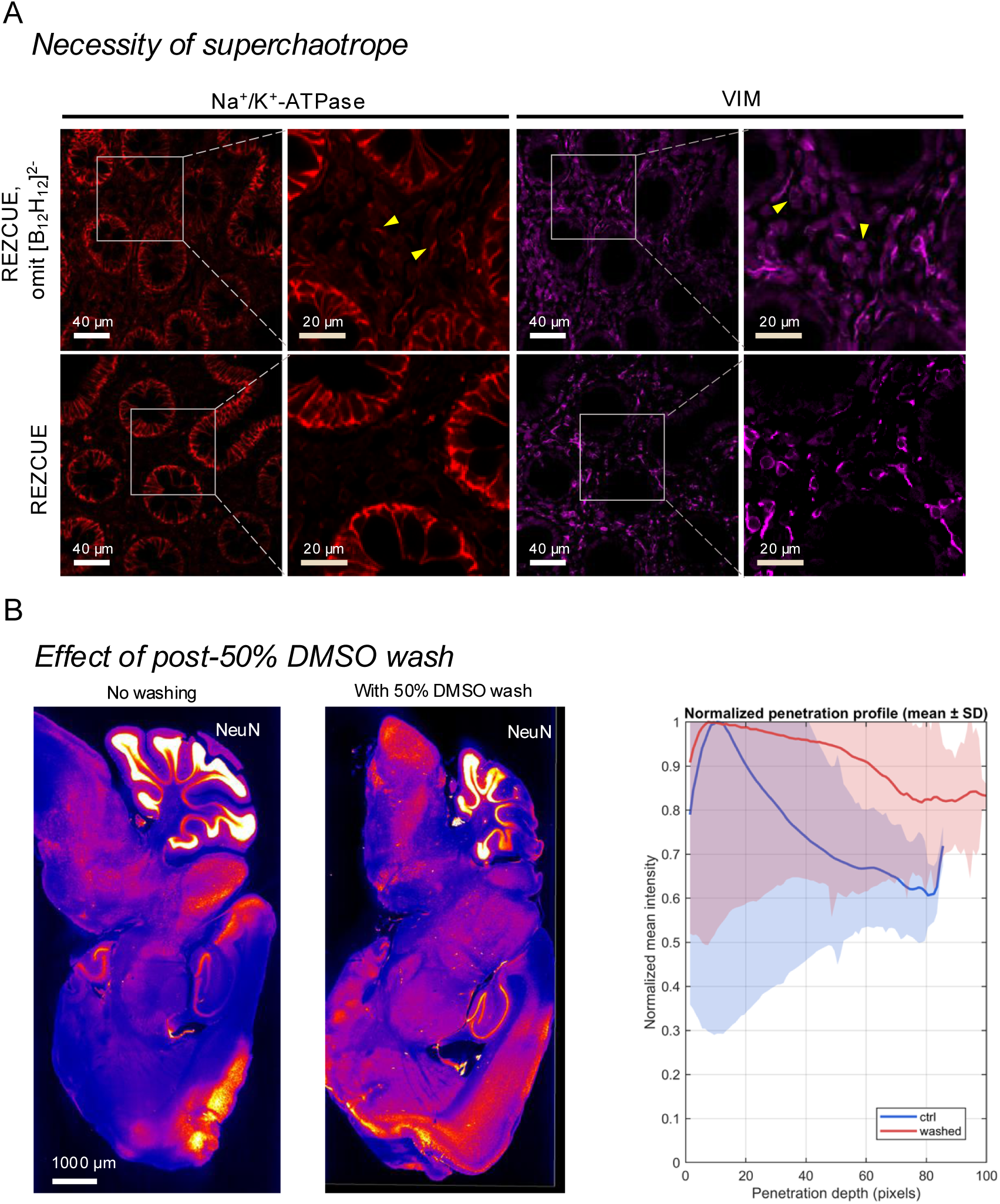
Refinement of aqueous antigen retrieval solution components and protocol. The Omission of Na_2_B_12_H_12_ in REZCUE antigen retrieval led to cytoplasmic non-specific staining. Tissue used: FFPE human colon, stained with anti-Na^+^/K^+^-ATPase (in red) and anti-VIM antibodies (in magenta). **B**. Addition of a 50% DMSO wash immediately after REZCUE antigen retrieval in the aqueous cocktail solution of Zwittergent SB3-16, Na_2_B_12_H_12_, and αCD further improved antibody penetration. Tissue used: half adult mouse brains stained with anti-NeuN antibodies. Left: representative mid-Z-slice of the image volume; right: quantified NeuN immunostaining intensity (means: solid lines, S.D.s: shaded regions) across all tissue pixels ordered by distance transformation from the nearest tissue surface.

**Supplementary Figure 5.**
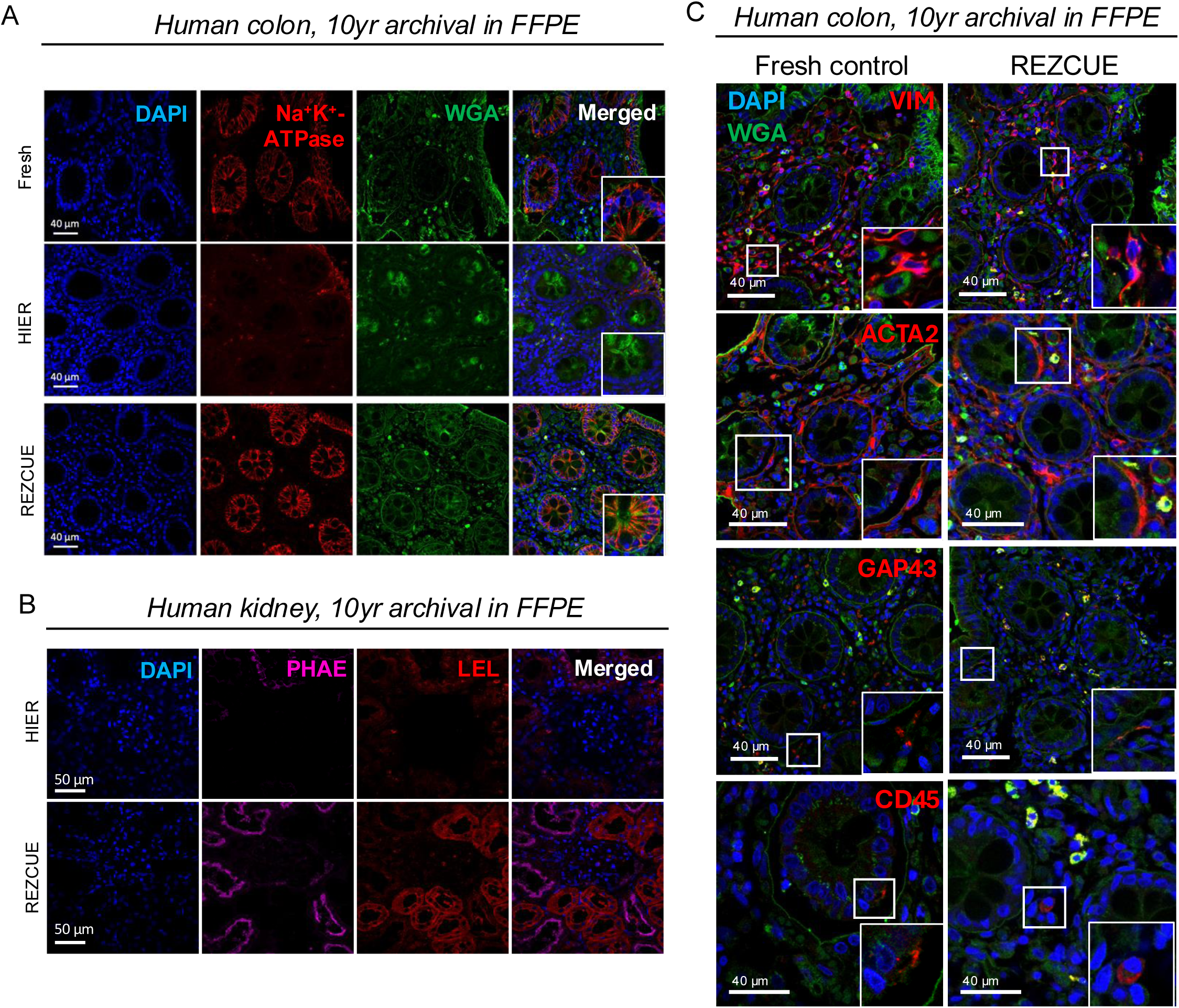
REZCUE enables high-quality 3D staining of human FFPE tissue samples that had been archived for years. Only representative z-slices at 50 µm imaging depth were shown for clarity. **A.** Assessment of 3D immunostaining quality of fresh-fixed, non-FFPE human colon tissue and human colon tissue that had been archived for 10 years after REZCUE processing. **B.** Images of human kidney tissue that had been archived for 10 years and were processed with conventional HIER or REZCUE. Freshly fixed human kidney tissues were not available during the study period. PHAE: Phaseolus vulgaris erythroagglutinin. **C.** Images of fresh-fixed, non-FFPE human colon tissue and human colon tissue that had been archived for 10 years that was processed with conventional HIER or REZCUE.

**Supplementary Figure 6.**
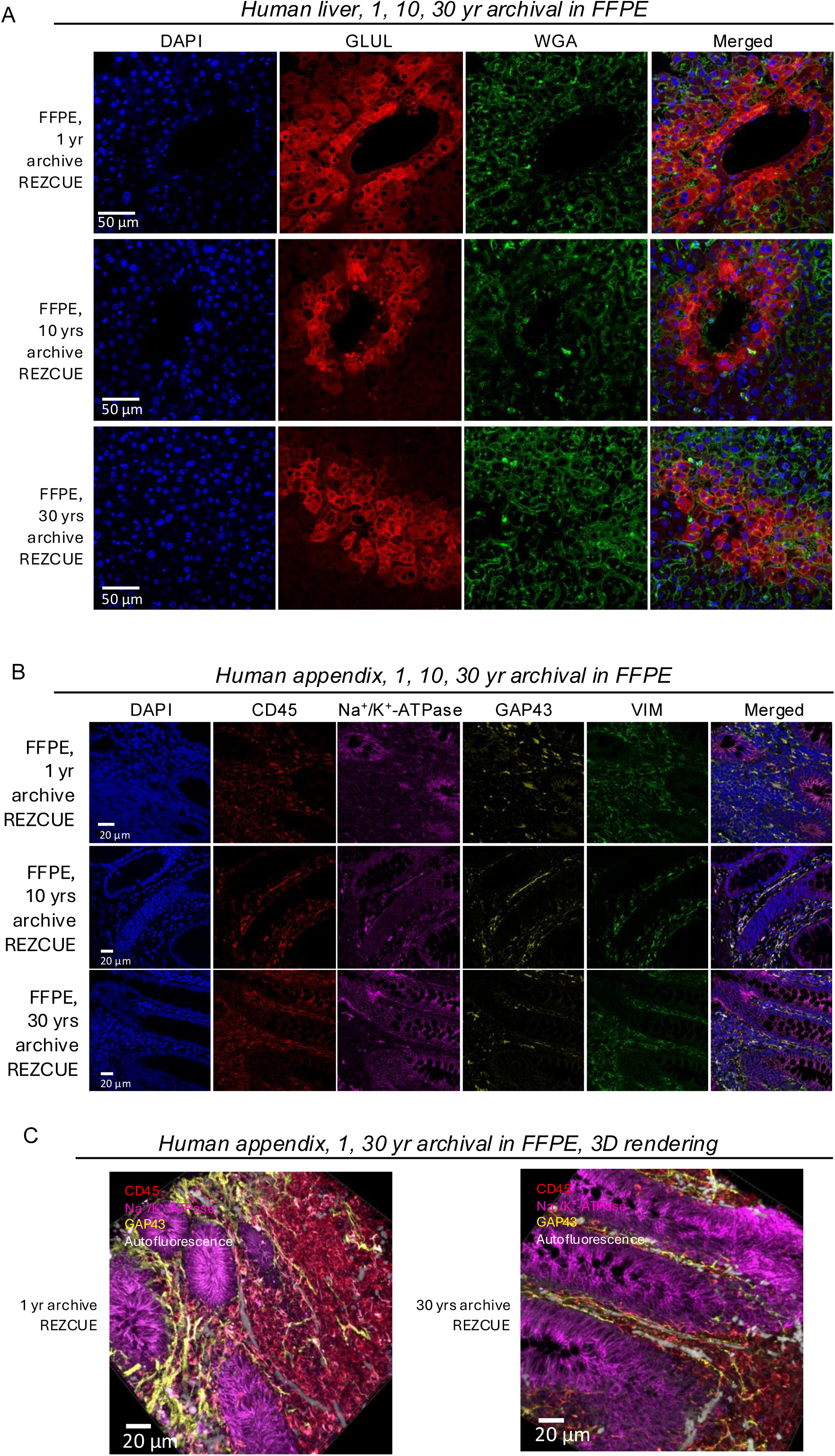
Additional images for 1, 10, and 30-year archived FFPE human samples with REZCUE. **A** and **B**. Assessment of staining quality on samples processed by REZCUE with various duration of FFPE tissue archival. **A**. On the human liver. **B.** On human appendix. **C**. 3D rendering of the images in **B** for the 1 year and 30 year-archived sample.

**Supplementary Figure 7.**
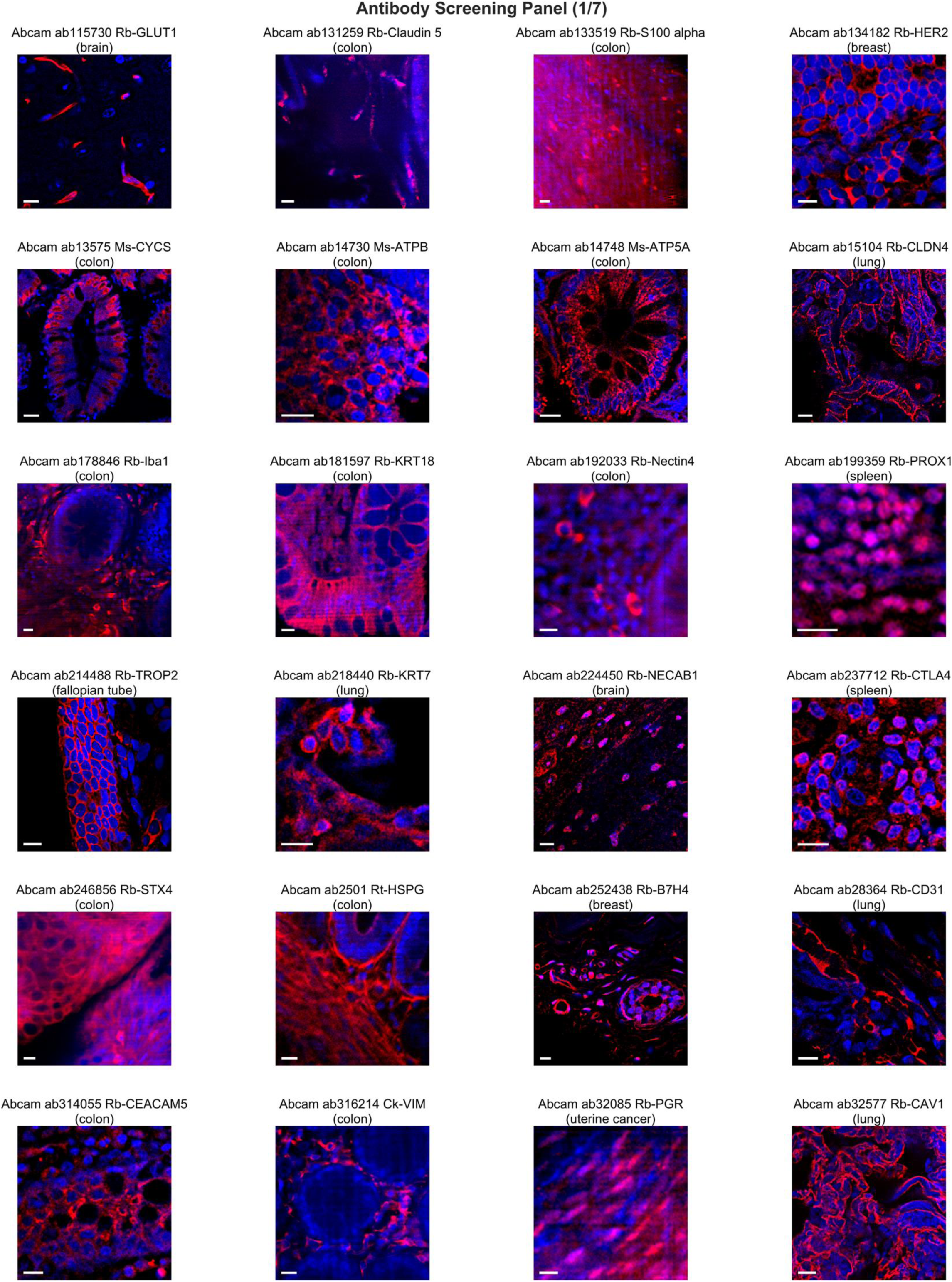
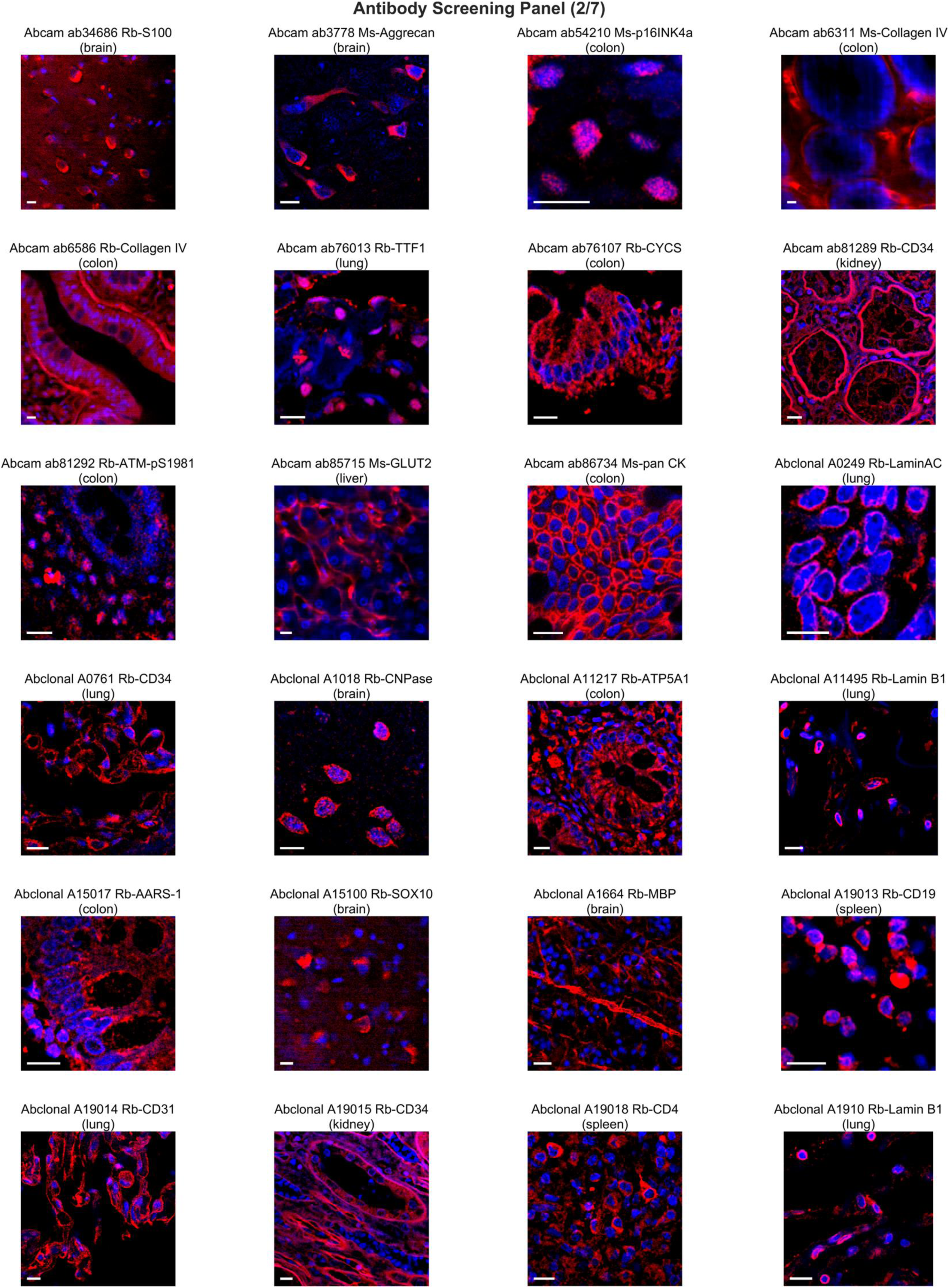
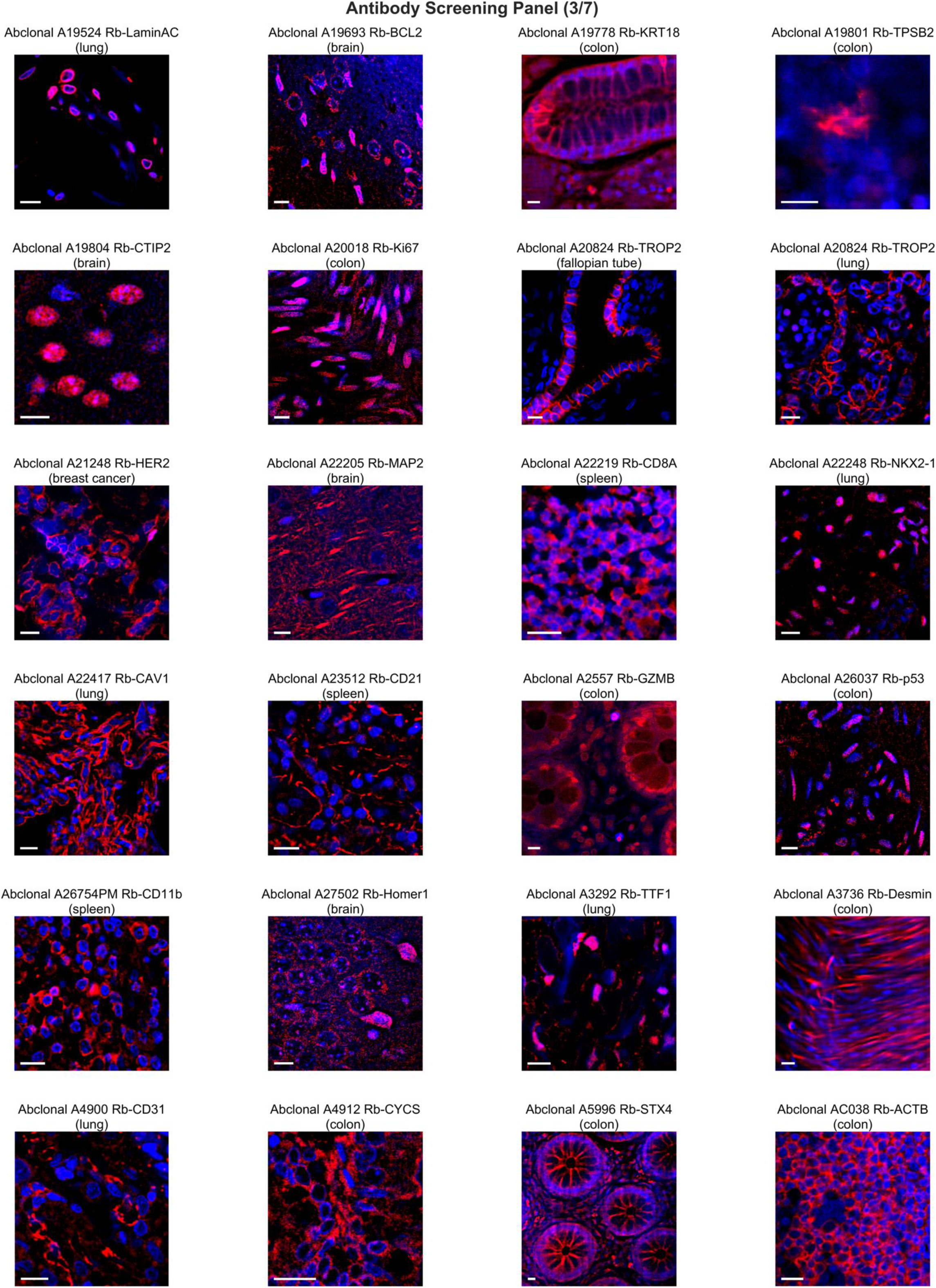
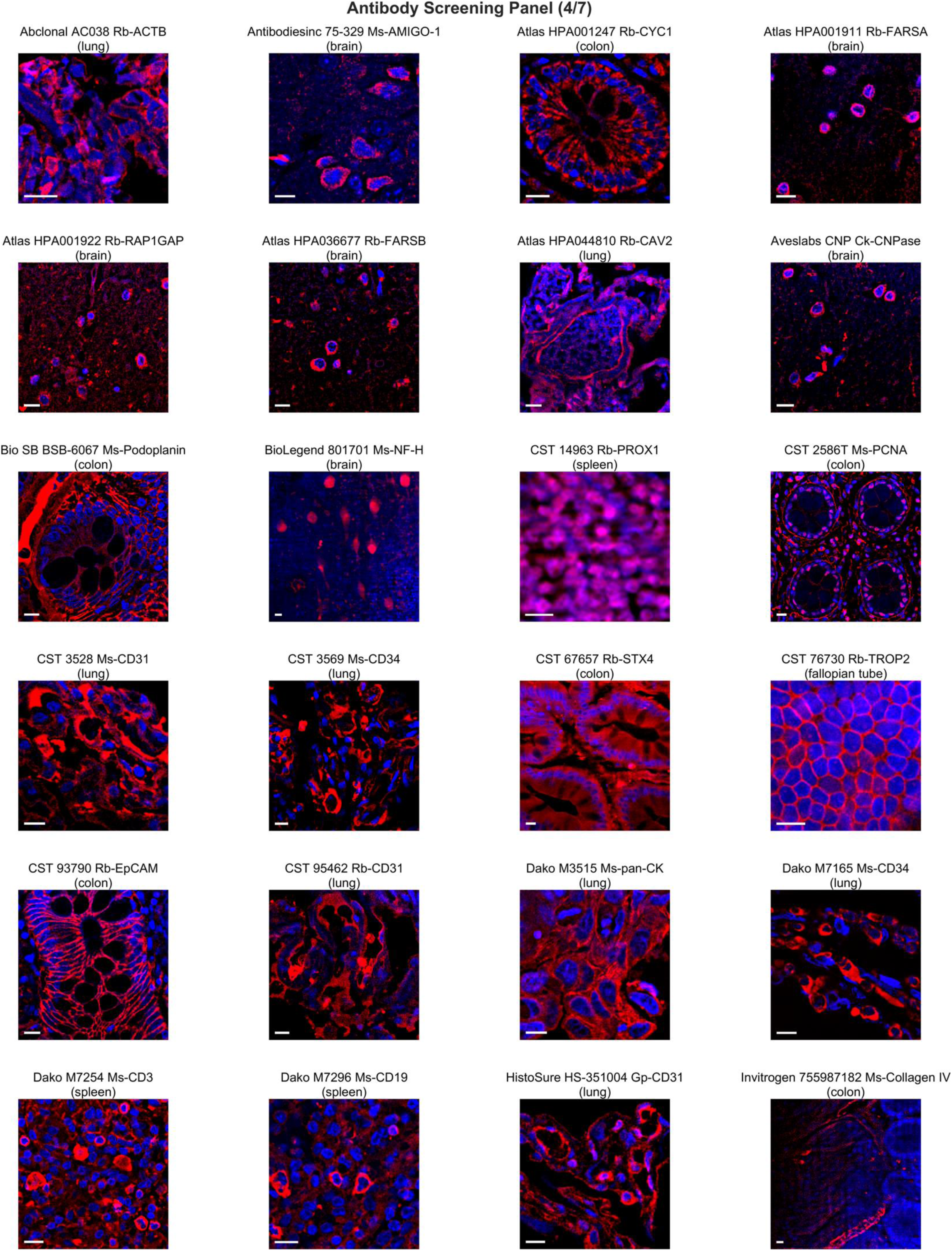
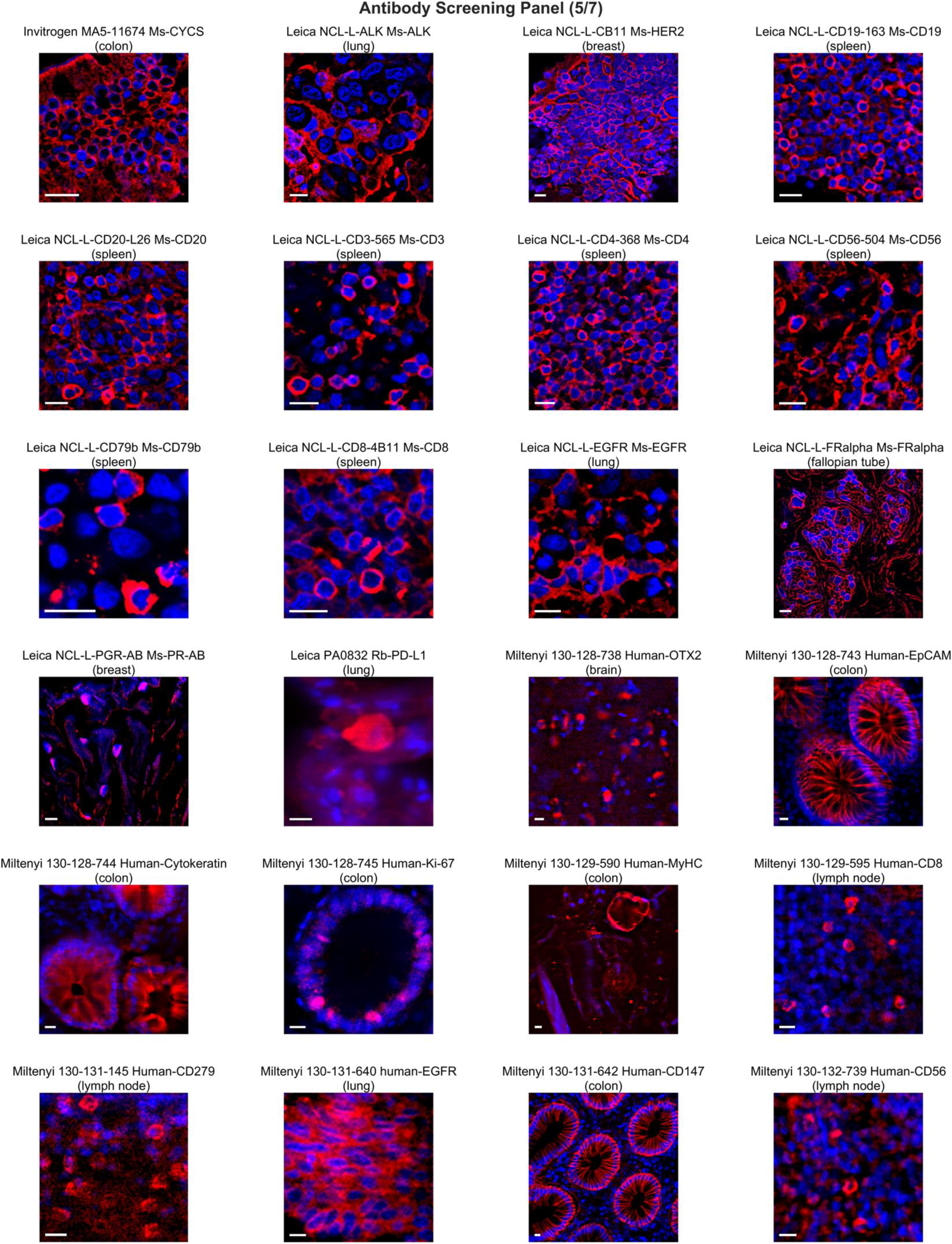
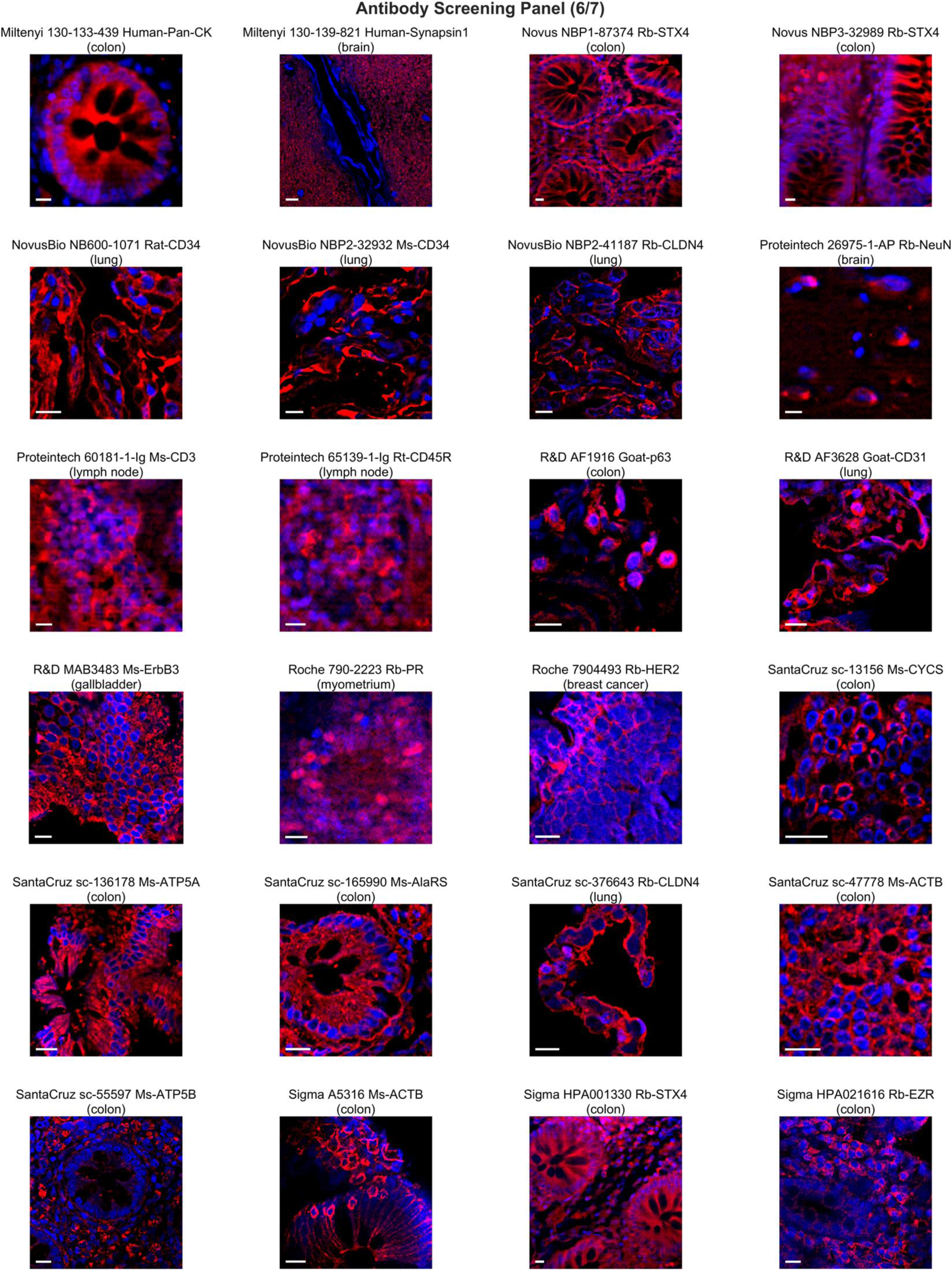
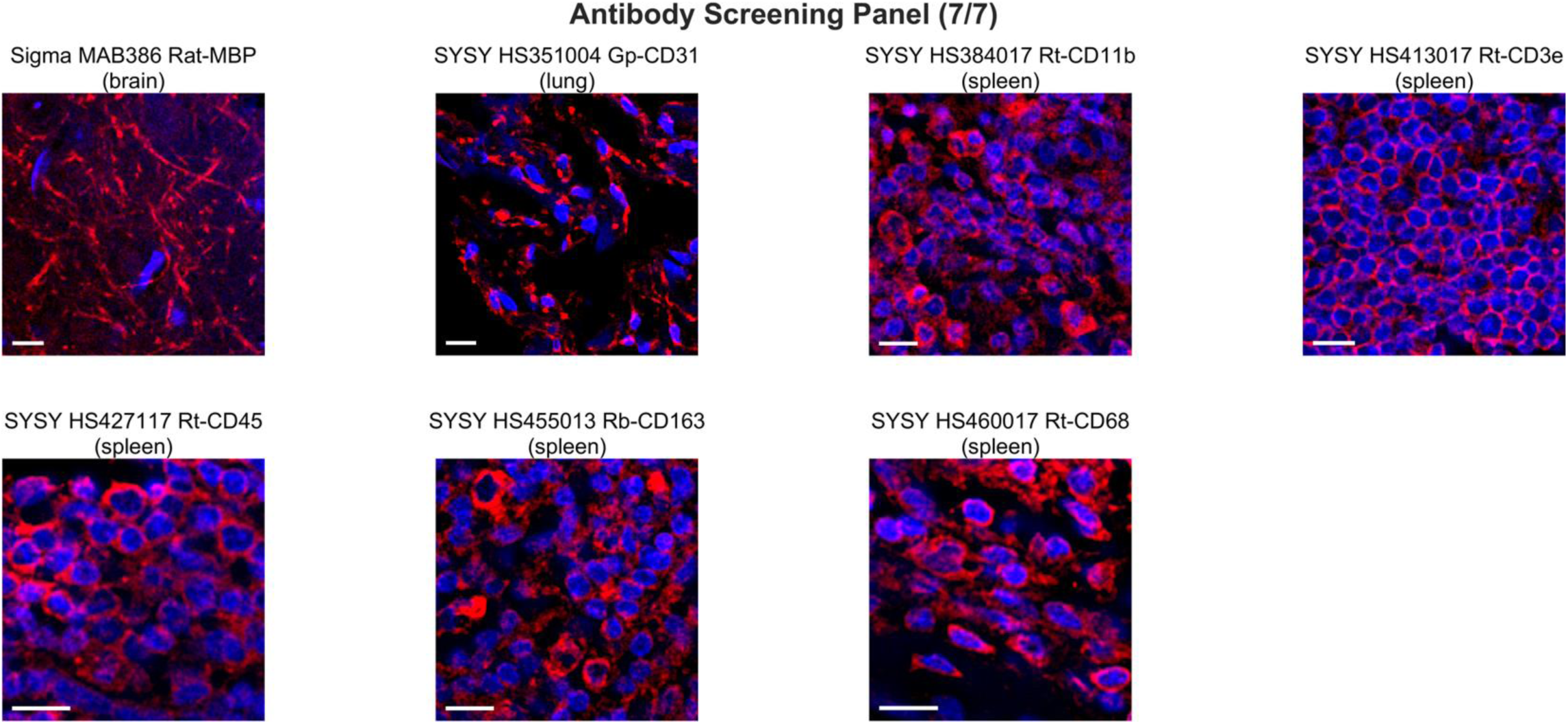
Validated 3D immunostaining results for archived human FFPE blocks after REZCUE. Only a single Z-slice was shown for clarity. Scale bars: 10 μm.

**Supplementary Figure 8.**
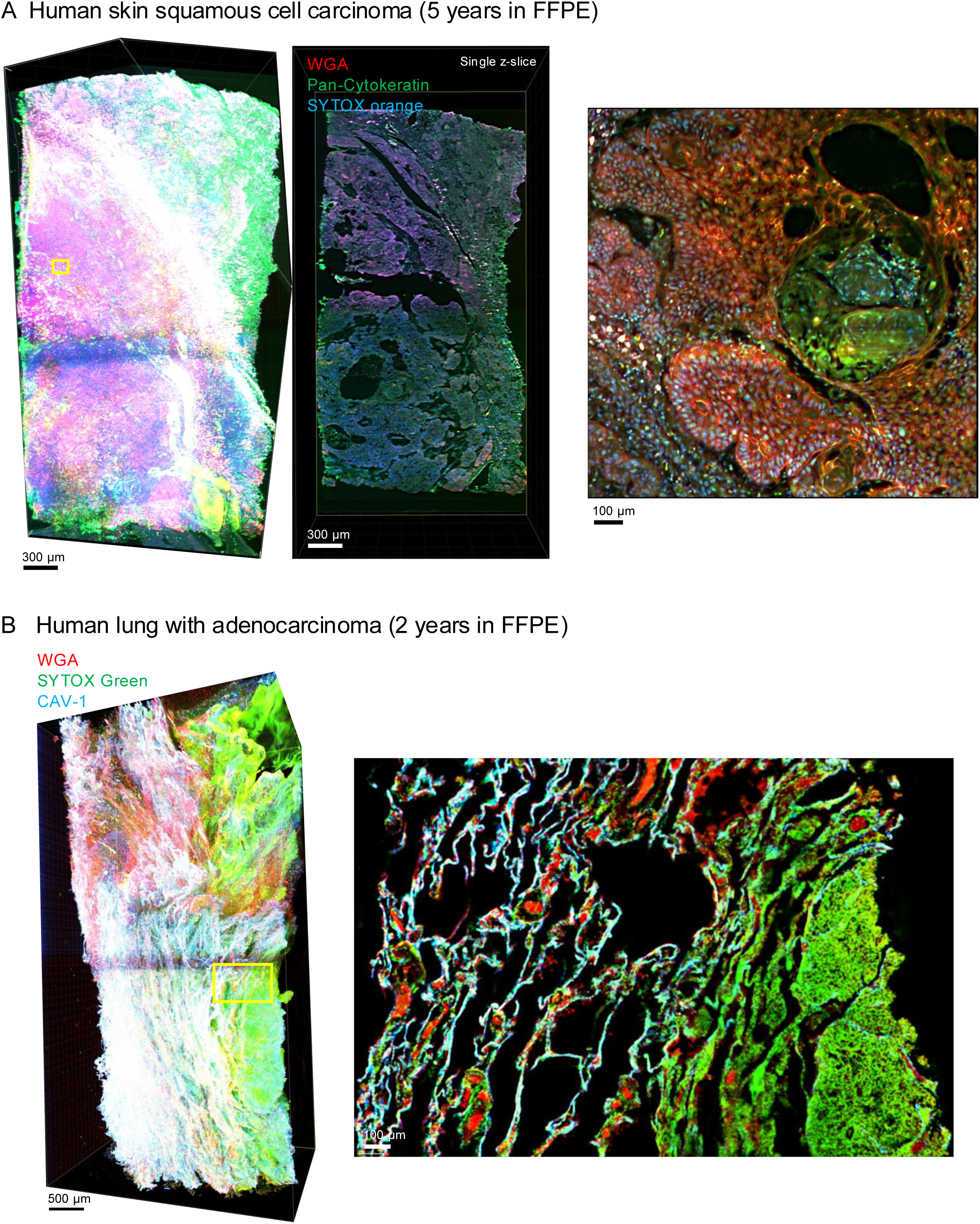
Large-scale 3D imaging of archived FFPE human tissues with REZCUE - Example 1. Scale bars and human organs as indicated. **A.** Overview of a 1.5mm × 5.5 × 3.5 mm human skin squamous cell carcinoma sample (left), a single mid-z-slice (middle), and a zoomed-in view highlighting a keratin pearl (right) corresponding to the yellow-boxed area of the left image. WGA: wheat germ agglutinin. **B.** Overview of a 2.5 mm × 8 mm × 5 mm human lung sample with focal adenocarcinoma (left). A single z-slice zoomed-in view of the yellow-boxed area is shown on the right, revealing the bulk tumor (dense cellularity, as demonstrated by SYTOX green) and adjacent normal alveoli (CAV1-positive structures).

**Supplementary Figure 9.**
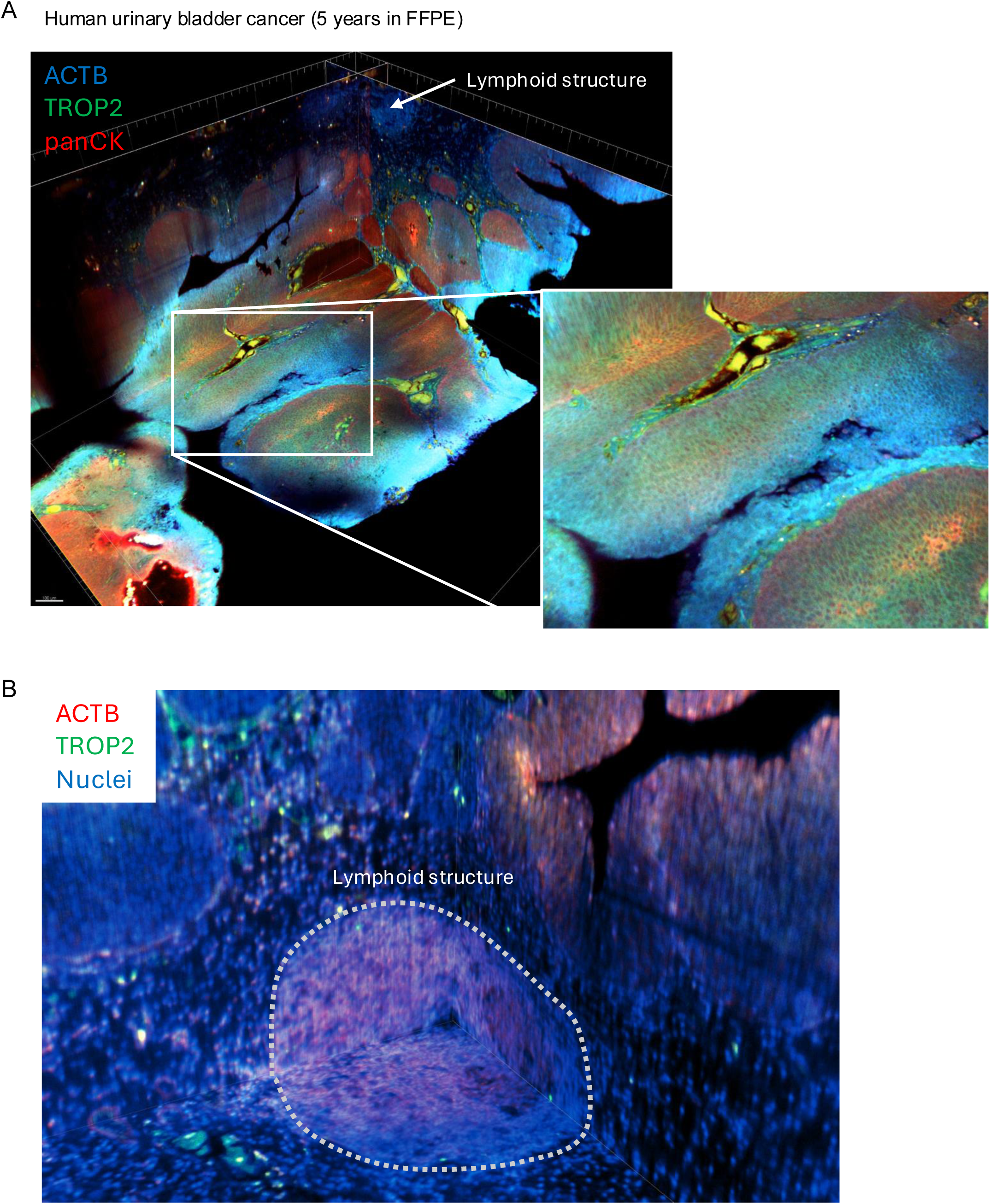
Large-scale 3D imaging of archived FFPE human tissues with REZCUE - Example 2. Scale bars and human organs as indicated. **A**. Triplex imaging of a 3 mm × 4 mm × 2 mm human bladder cancer with orthogonal sectioning view, stained for ACTB (highlighting stroma and immune cells), pan-cytokeratin (panCK, tumor cells and normal epithelium), and TROP2 (an antibody-drug conjugate target). The predominantly membranous expression of TROP2 on tumor cells is visible in the zoomed- in view. **B.** Another orthogonal sectioning view demonstrating lymphoid follicles, which are frequently encountered in bladder cancer tissues.

**Supplementary Figure 10.**
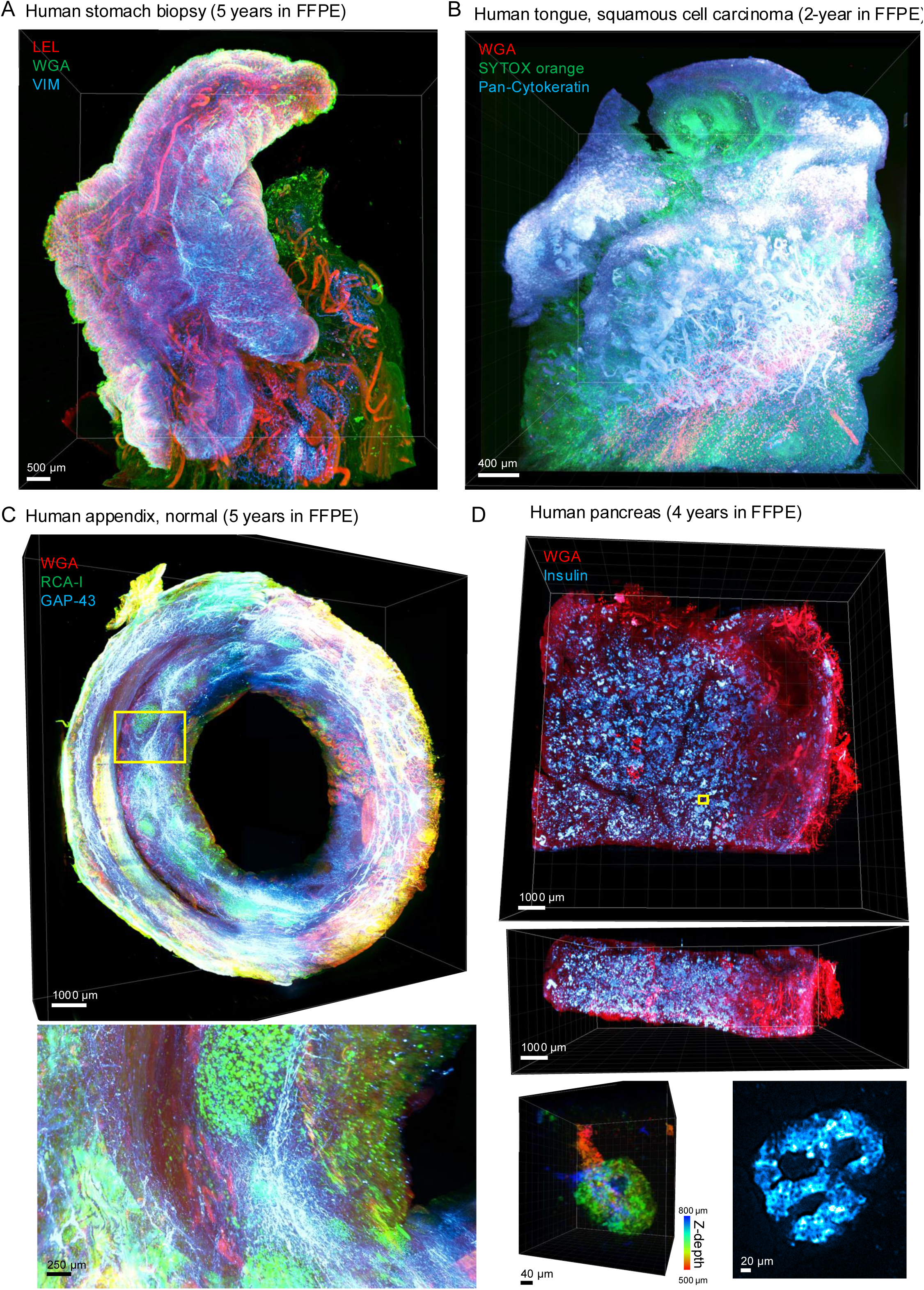
Large-scale 3D imaging of archived FFPE human tissues with REZCUE - Example 3. Scale bars and human organs as indicated. **A**. A 10 mm × 6 mm × 6 mm gastric biopsy tissue. LEL: Lycopersicon esculentum lectin, WGA: wheat germ agglutinin, VIM: vimentin. **B.** A 3.5 mm × 4 mm × 3 mm human tongue squamous cell carcinoma. **C**. A transverse cross-section of a whole human appendix (ø 10 mm × 3 mm) stained with WGA (highlights smooth muscle and epithelium), Ricinus communis agglutinin-I (RCA-I; highlights dendritic cells and macrophages), and an anti-GAP43 antibody (highlights nerves). A zoomed-in view of the yellow-boxed area shows dense nerve plexuses around the lymphoid structures. **D.** An 11 mm × 6 mm × 2.5 mm human pancreas tissue stained with WGA and anti-insulin antibodies with 3D rendering view along the z-axis (upper image) and x-axis (middle image), demonstrating full and homogeneous penetration of the antibody. The lower two images show a 3D rendering of a single islet of Langerhans with color-coded Z-depth and a single z-slice view for morphology for the insulin-labelled channel.

**Supplementary Figure 11.**
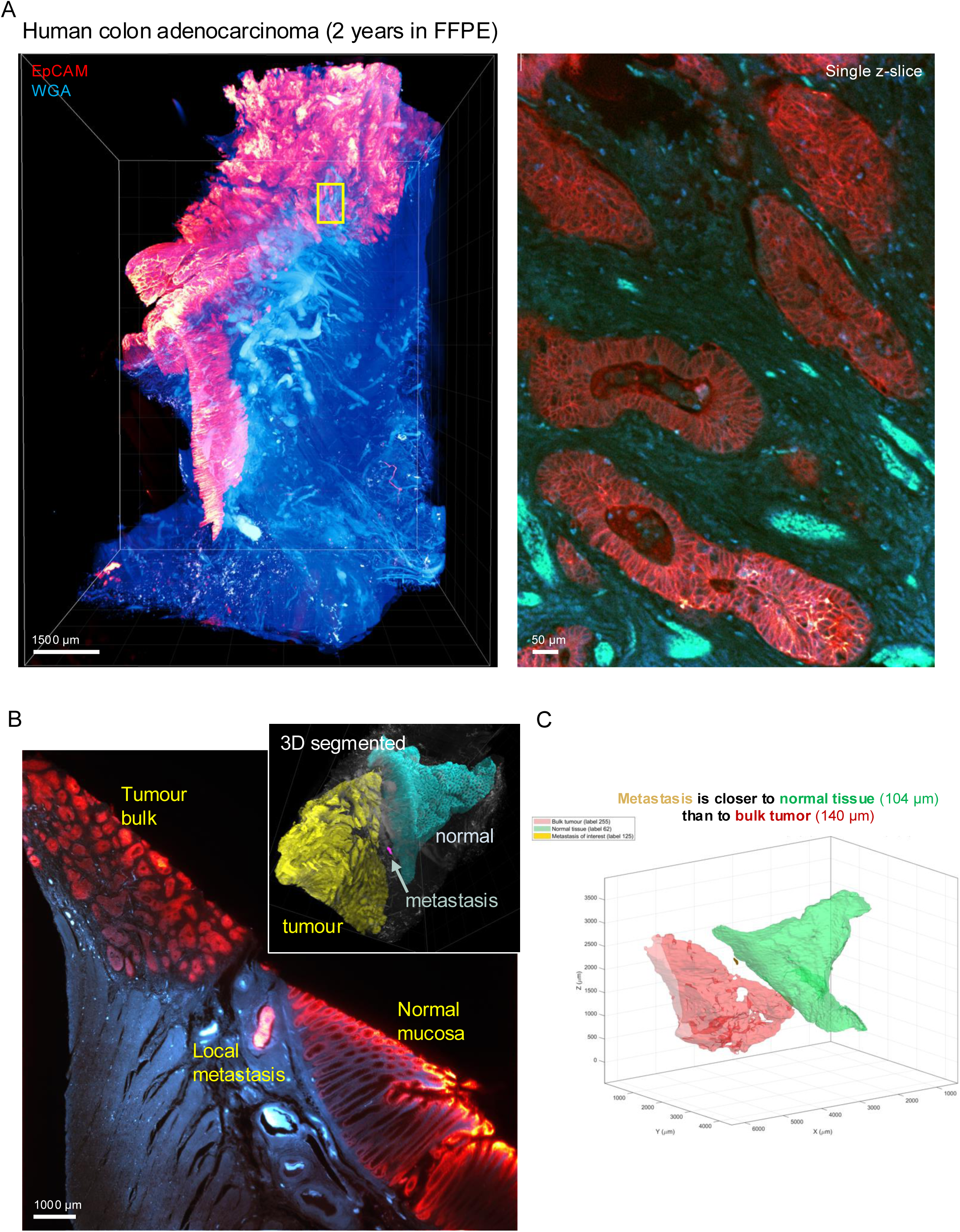
Large-scale 3D imaging of archived FFPE human tissues with REZCUE - Example 4. Scale bars and human organs as indicated. **A.** A 6 mm × 9 mm × 4.5 mm human colon tissue with opposing normal and cancerous epithelia, stained for EpCAM that highlights individual cancer cells. WGA: wheat germ agglutinin, highlights epithelia, smooth muscle, and red blood cells in vessels. Right image: zoomed-in and single z-slice view of the yellow-boxed region for cellular morphology assessment. **B.** A local vascular metastasis was found to have spread relatively far away from the tumor bulk, and instead close to the normal tissue region. The inset shows 3D segmented and color-coded tumor region (in yellow), normal tissue region (in cyan), and the vascular metastasis in magenta. **C.** 3D surface-to-surface distance computation confirms the metastasis is closer to the normal epithelium (104 µm) than to the tumor bulk (140 µm).

**Supplementary Figure 12.**
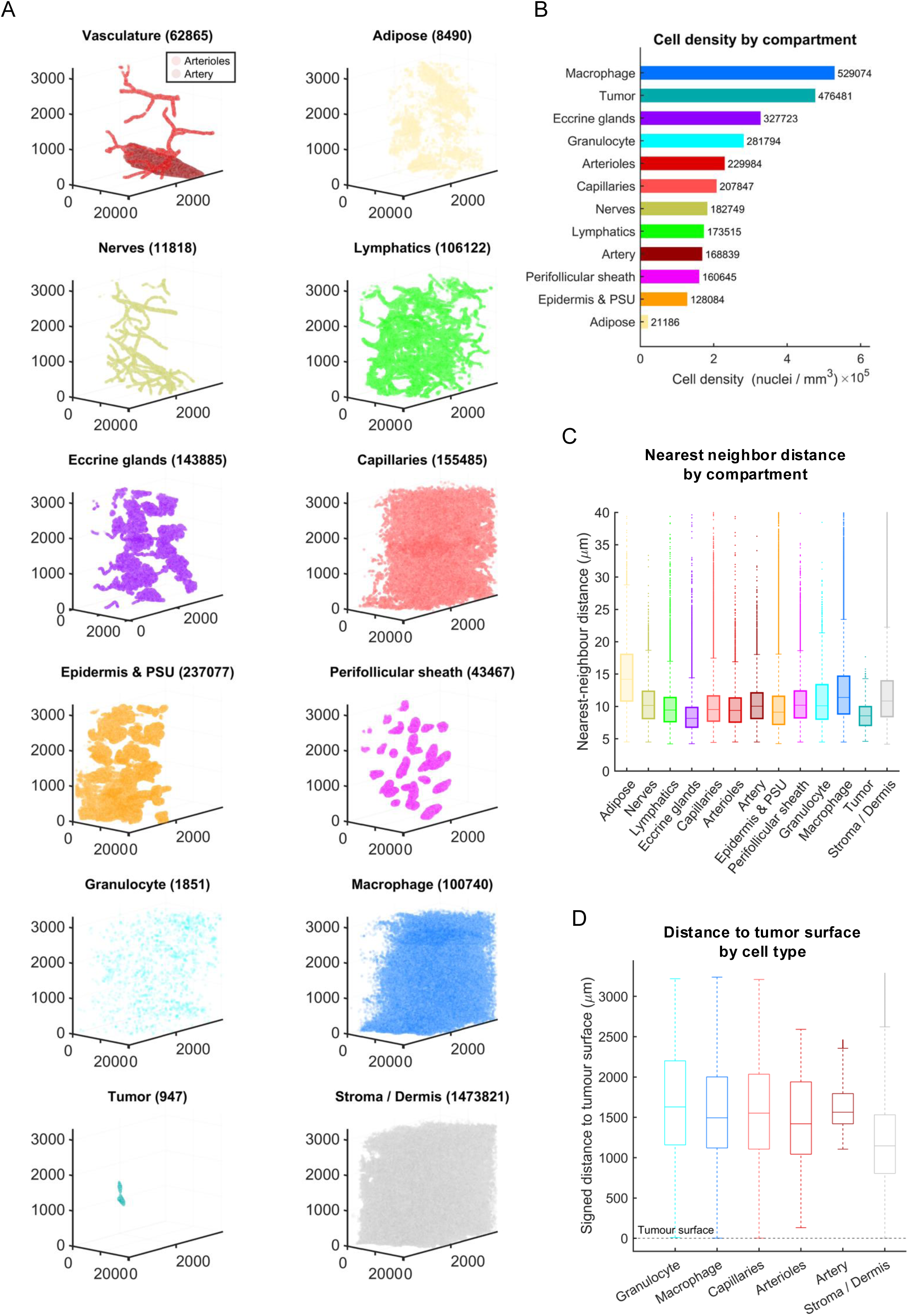
Segmented nuclei and their localized compartments in a REZCUE-treated normal skin tissue adjacent to a sinonasal carcinoma. **A.** Scatter plots of all nuclei classified according to their compartment. Numbers in parenthesis indicate the number of cells found in each compartment. **B.** Density of cells per compartment, which is related to the nuclear density per unit segmented volume, consisting mostly of cytoplasm. Hence the plot roughly reflects the nuclear-to-cytoplasmic volume ratio. **C**. 3D nearest neighbor distance of cells in each compartment. **D**. The closest cells to the tumor surface, classified by cell type.

**Supplementary Figure 13.**
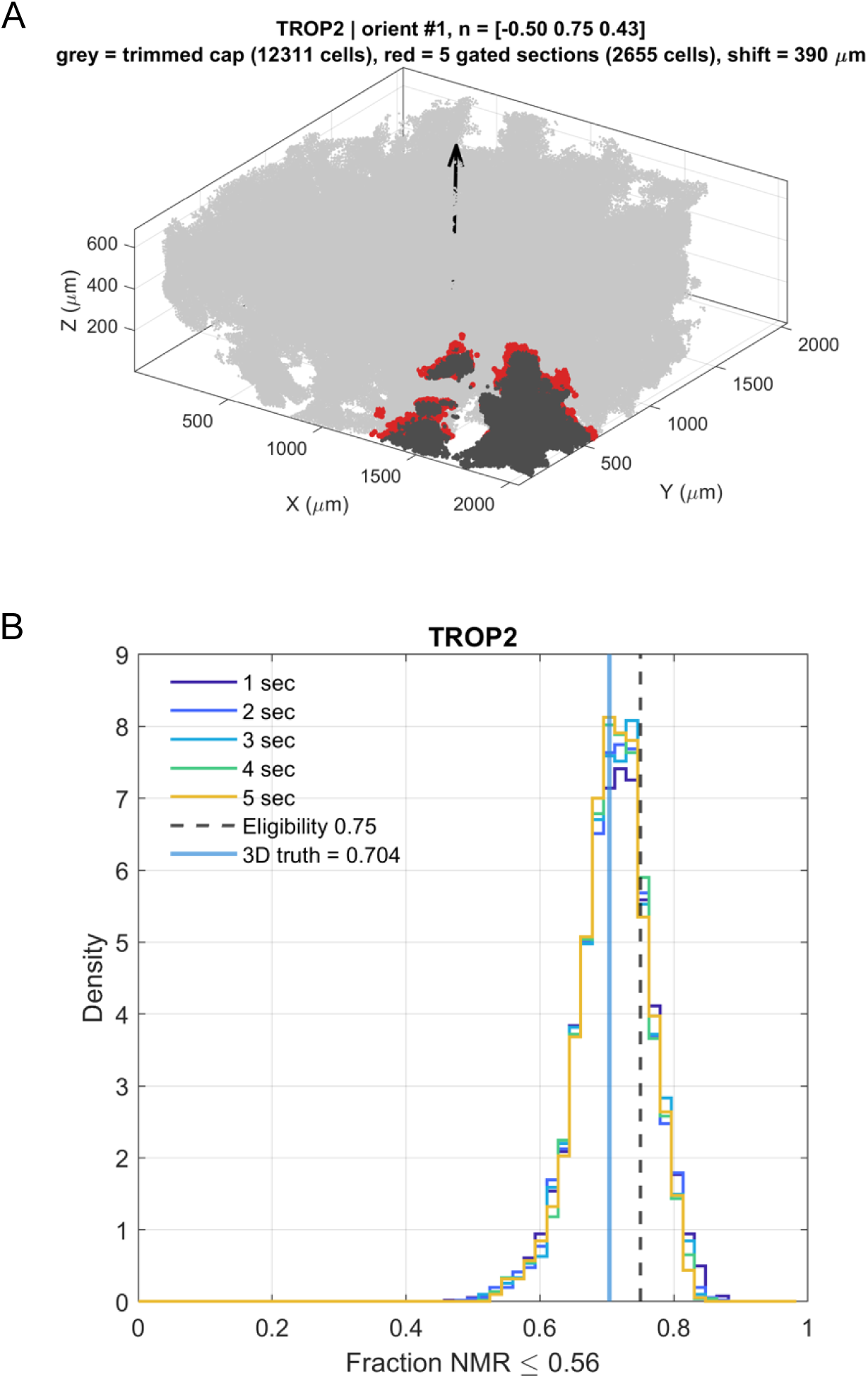
In silico randomly oriented 2D sectioning of lung biopsy and its impact on molecular diagnostic accuracy. **A**. Illustrated example sectioning of the sample for the normal vector [-0.50, 0.75, 0.43]. The section is trimmed (cells indicated in black are removed) until a single section yields a qualifying minimum of 500 cells (resulting on-section cells highlighted in red). **B.** TROP2 2D QCS (a measure of the fraction of cells having NMR ≤ 0.56, x-axis) distribution from 5000 randomly oriented sections. The dashed line indicates 75% of on-section cells with NMR ≤ 0.56, above which the patient will qualify for TROP2 ADC treatment. The 3D ground truth across all 537,553 cells indicates this patient *does not* qualify; hence, the sampled sections above the 0.75 cut-off are false positives. We repeated the simulation, varying the number of qualifying sections sampled from 1 to 5, with little difference in the final result.

**Supplementary Figure 14.**
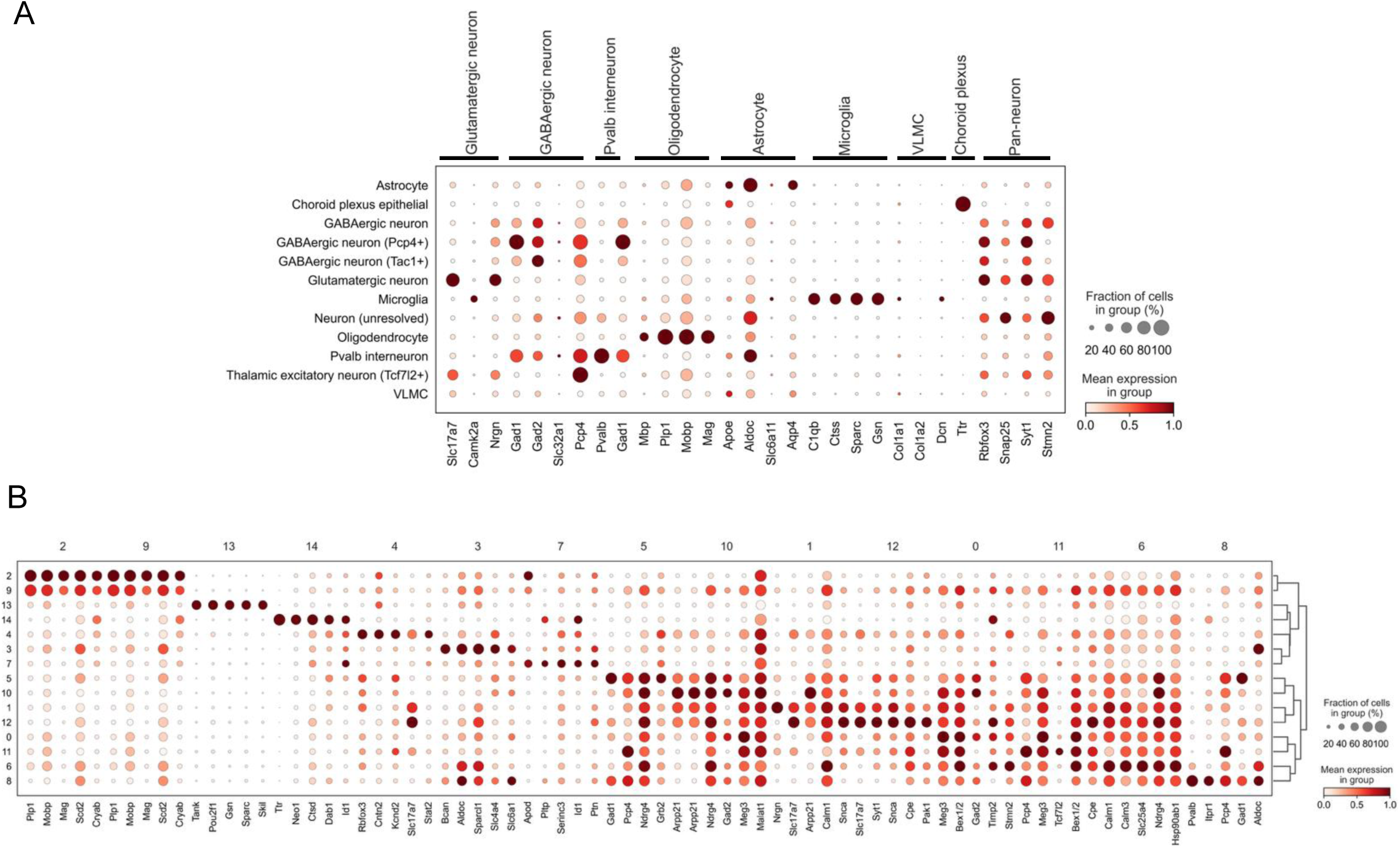
Spatial transcriptomics comparison experiments for REZCUE-treated FFPE tissues versus paired control tissues in mouse brain. **A.** Scatterplot of known neuronal cell type specific markers. **B.** Scatter plot of expression level for these known neuronal cell type specific markers for each cluster.

**Supplementary Figure 15.**
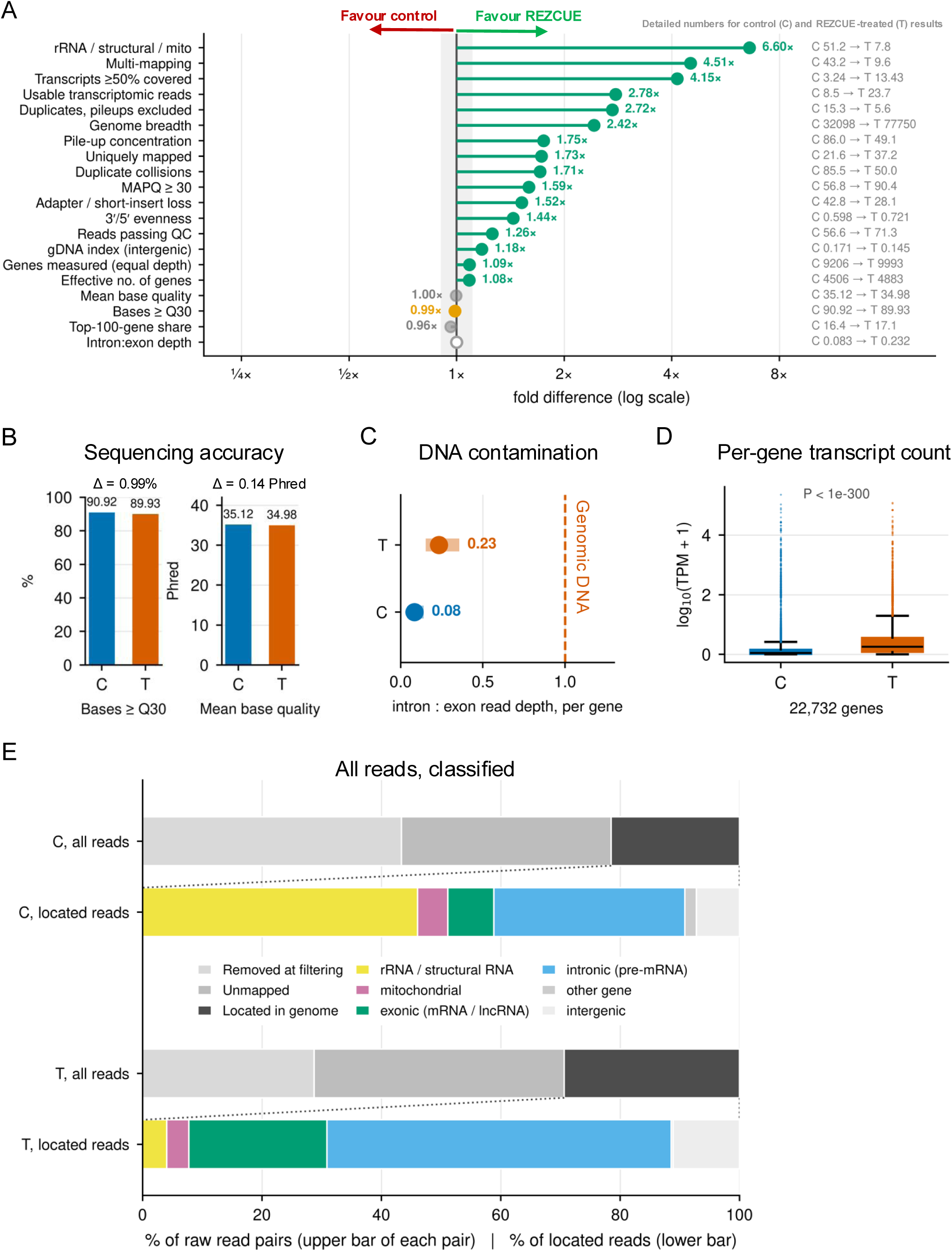
Comparison of bulk RNA sequencing results between REZCUE-treated adult mouse hemibrain and its paired contralateral control. **A.** Forest plot of major metrics comparing the result of bulk RNA sequencing quality control assessment. The specific numerical value of each metric is indicated for control (C) and REZCUE-treated (T) samples in grey text. **B**. Detailed comparison on the two metrics where REZCUE-treated is slightly worse than C, including base quality ≥Q30 and mean base quality. **C**. Assessment of possibility of genomic DNA contamination by intron:exon read depth per gene. Genomic DNA will have a value of 1.0. **D**. Per-gene transcript count (expressed as TPM) for each sample. **E**. Classification of all raw sequencing reads and mapped reads. REZCUE yielded more mRNA and pre-mRNA reads than control.

**Supplementary Figure 16.**
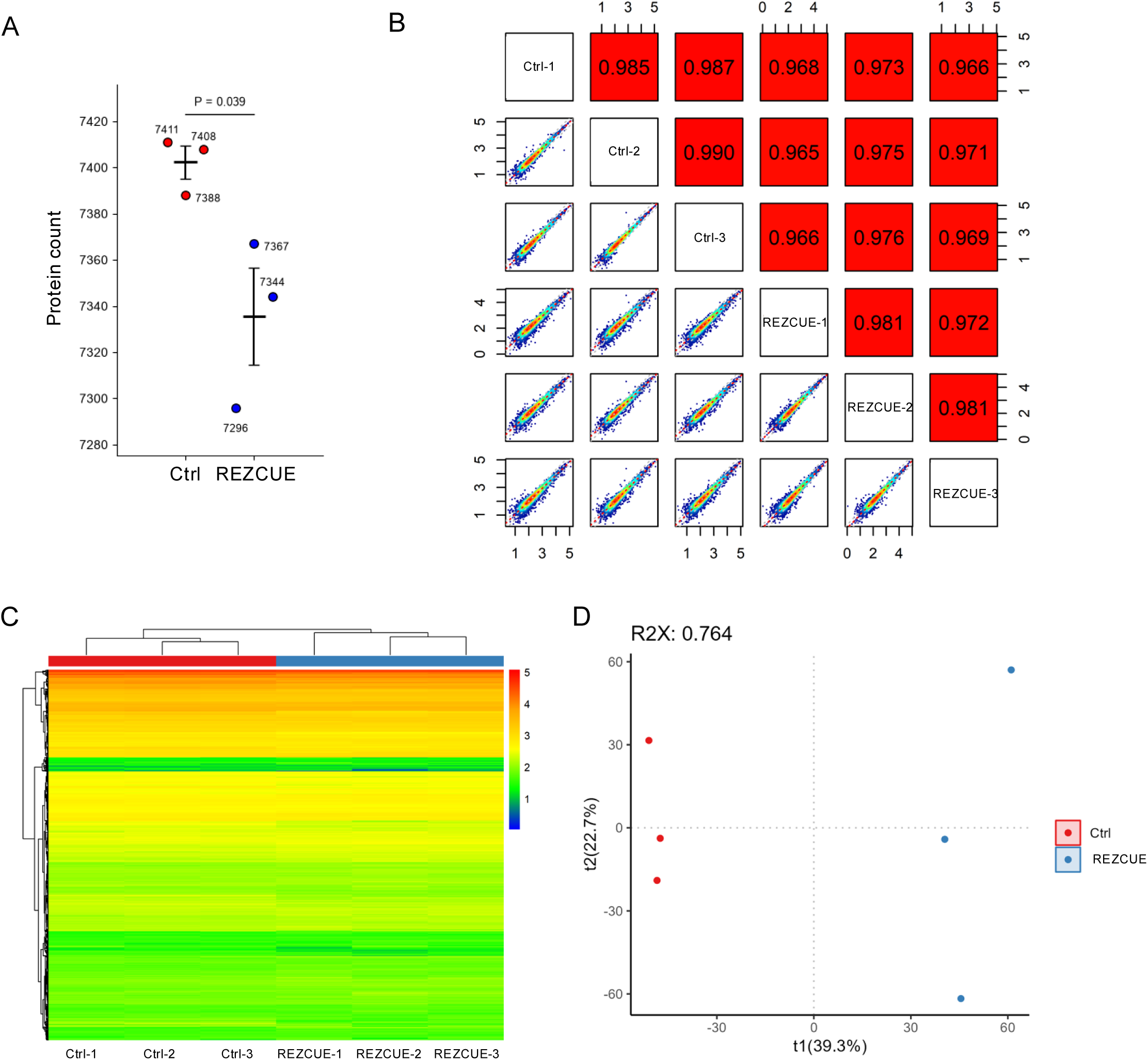
Additional data on post-REZCUE spatial proteomics by laser microdissection and mass spectrometry, compared with a paired non- REZCUE-treated FFPE control. A mouse FFPE hemibrain was REZCUE-treated (labelled as Test) while the other half was not, acting as the control (Ctrl). N=3 cerebellar slices per group were used for comparison, and sections were obtained at similar anatomical depths and orientations. **A.** Total protein count detected per section per group. REZCUE-treated samples have a slight but statistically significant decrease in detected proteins. Bar and error bars: Mean and S.D., *P*-value by 2-sided two-sample t-test. **B.** Paired sample comparison matrix for all detected proteins. The numbers displayed are Pearson correlation coefficients. **C.** Hierarchical clustering of all 6 sections. **D**. Principal components analysis (PCA) and clustering of all 6 sections. R2X: cumulative fraction of variation explained by all the calculated principal components.

**Supplementary Figure 17.**
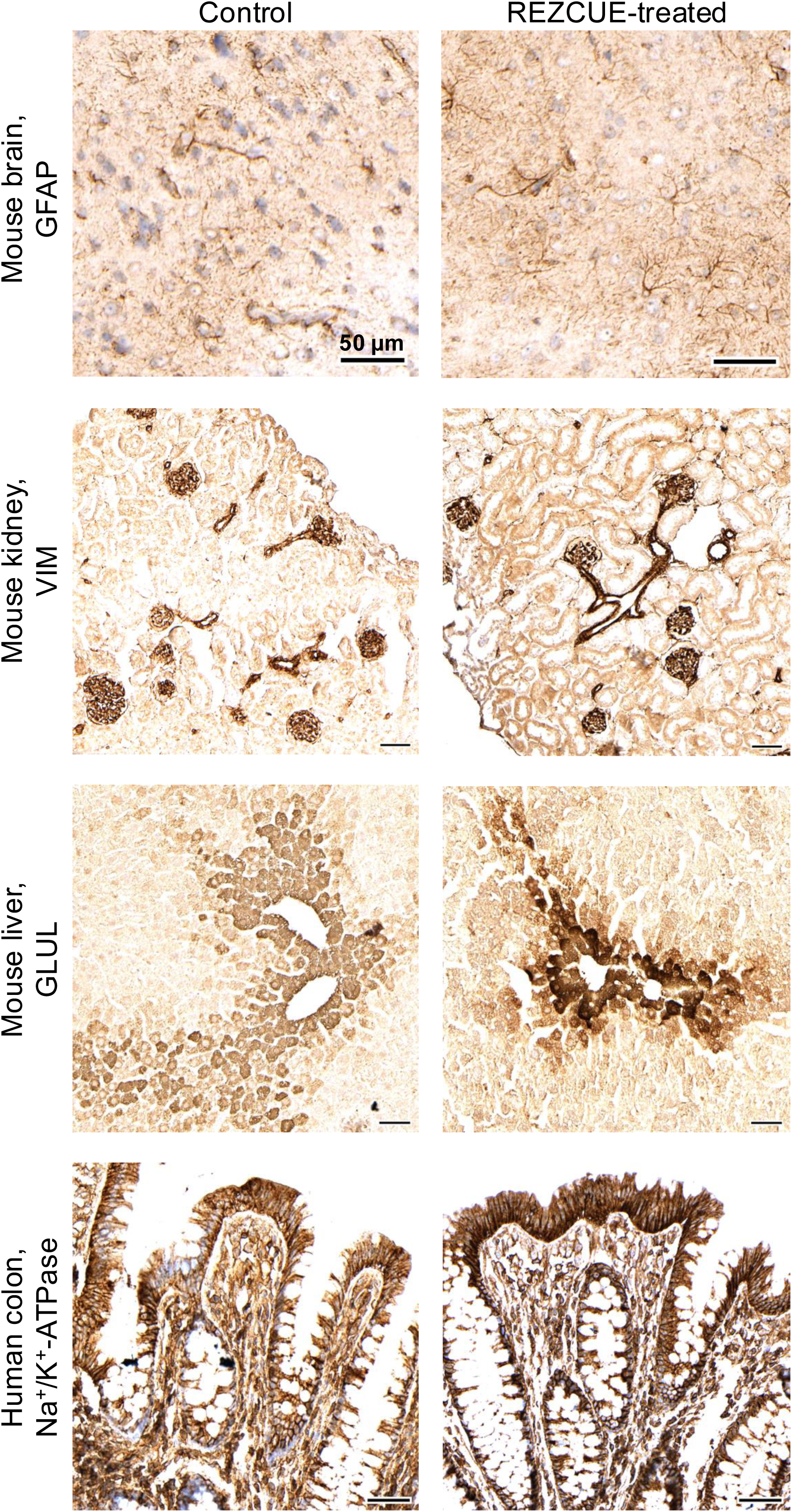
Post-REZCUE 2D immunohistochemistry, compared with matched control. REZCUE–treated FFPE tissues were re-embedded and sectioned along with matched control tissues. The sections were then stained with conventional 2D chromogenic immunohistochemistry.

**Supplementary Figure 18.**
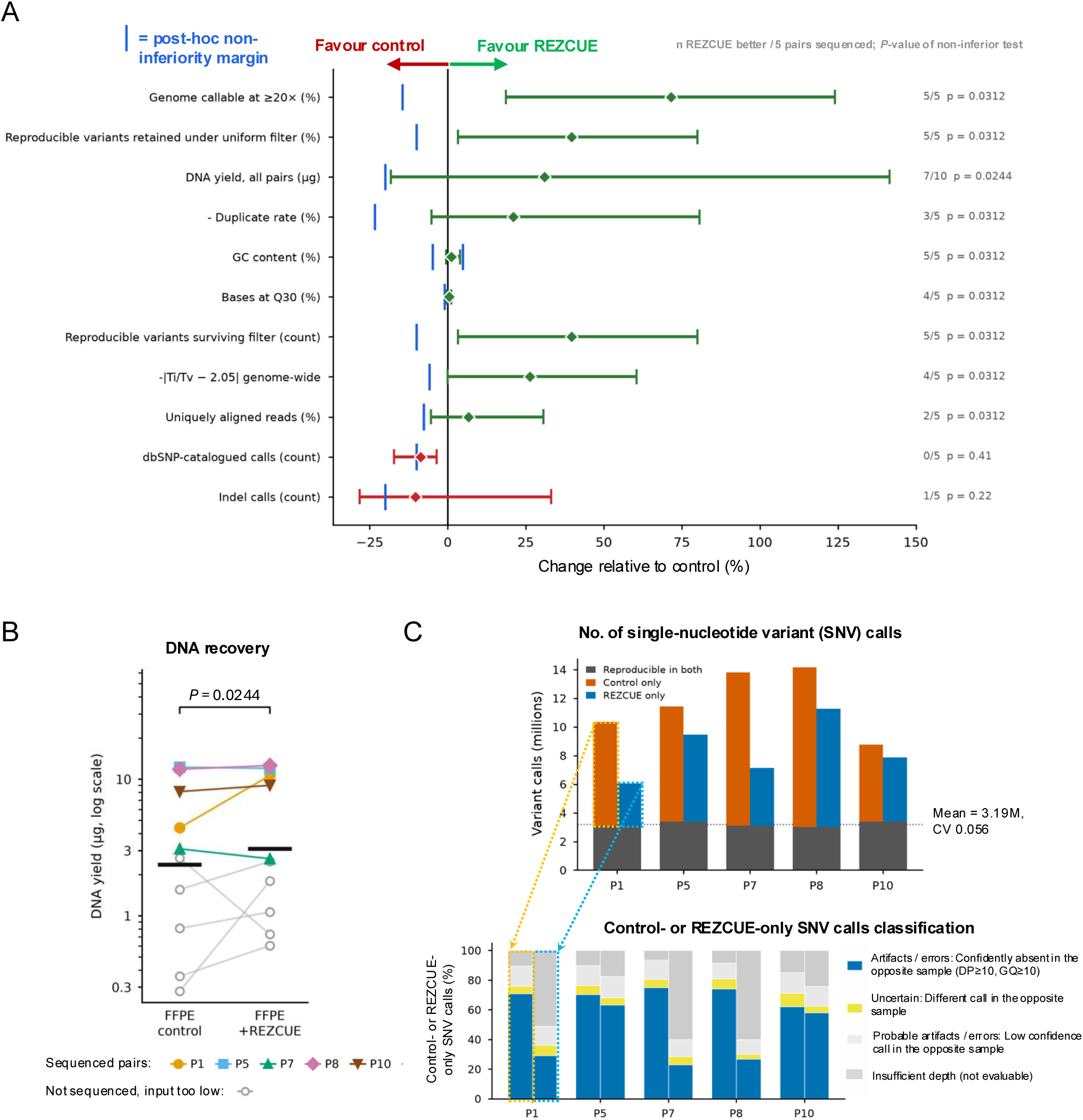
Comparison of whole genome sequencing (WGS) results between REZCUE-treated sample and its paired control. **A.** Forest plot of major metrics showcasing quality control metrics for whole genome sequencing between REZCUE and the control. All *P*-values were computed by a one-sided Wilcoxon signed- rank test to test for non-inferiority from a post-hoc margin of 10% or 5 points. **B**. DNA yields of all 10 sample pairs, only the top 5 pairs (pair 1, 5, 7, 8, 10) yielded sufficient input material and proceeded to sequencing. Middle bar: geometric mean of log(DNA yield in μg). **C.** (Upper) SNVs called in both the REZCUE-treated and control samples of each matched pair (n = 5 pairs) were highly reproducible across pairs (mean 3.19M shared SNVs; coefficient of variation = 5.6%), consistent with published estimates for a typical human genome (3.2–3.8M SNVs). This shared SNV count serves as a stable reference for evaluating the calls unique to each arm — SNVs called only in the control (orange) or only in the REZCUE-treated sample (blue) — which are extracted and classified below. (Lower) A single-arm SNV call was classified as a likely artifact or error when the opposite arm had high-confidence coverage at that position (read depth [DP] ≥ 20; genotype quality [GQ] ≥ 10) but did not call a variant there. REZCUE-treated samples showed a lower fraction of likely artifactual/erroneous SNV calls than their paired controls in all 5 pairs (one-sided Wilcoxon signed-rank test, *p* = 0.031).

**Supplementary Figure 19.**
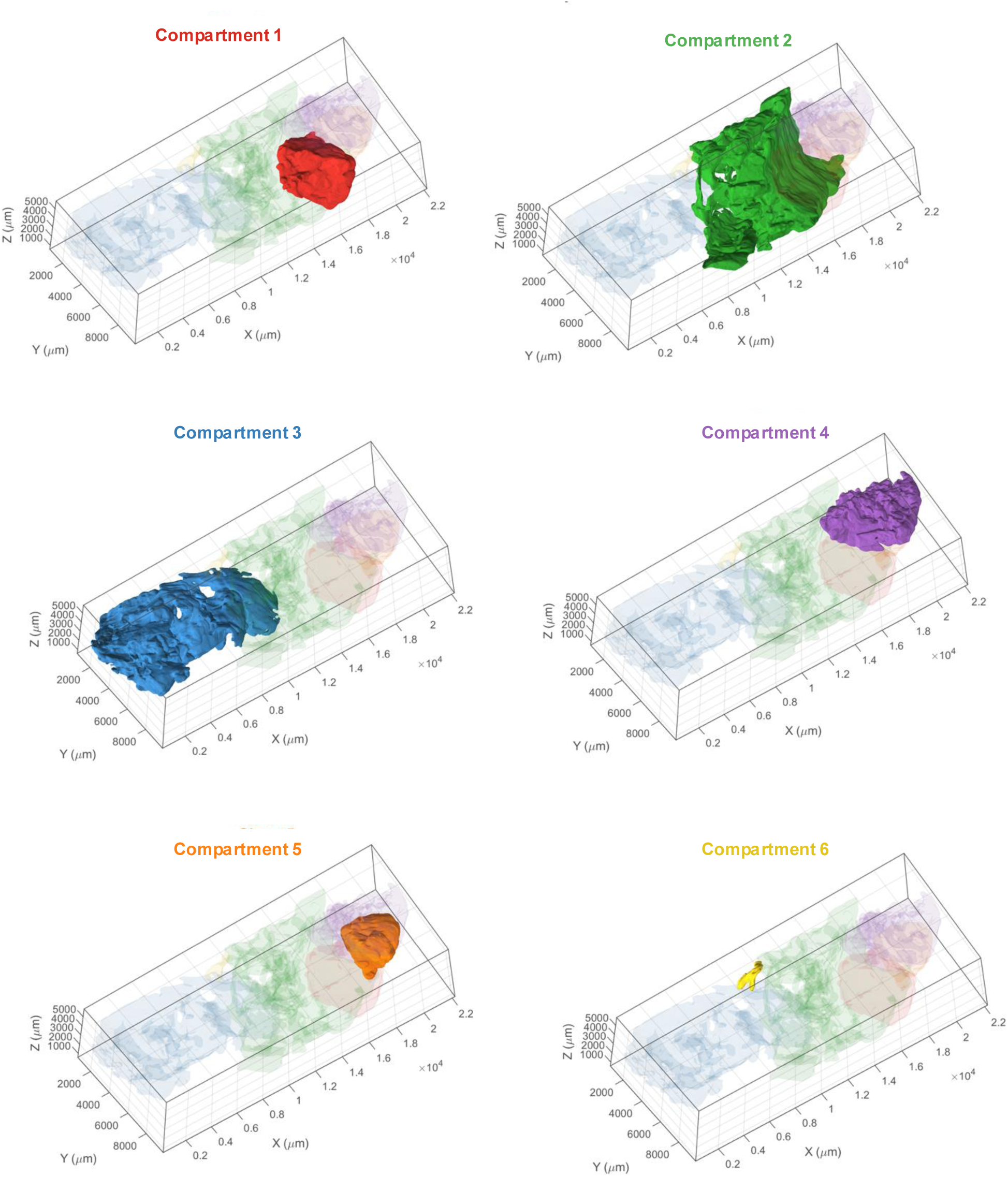
3D-segmented spatially discontinuous tumour compartments in a breast cancer sample. The figure shows surface renderings of each tumor compartment’s boundary.

**Supplementary Figure 20.**
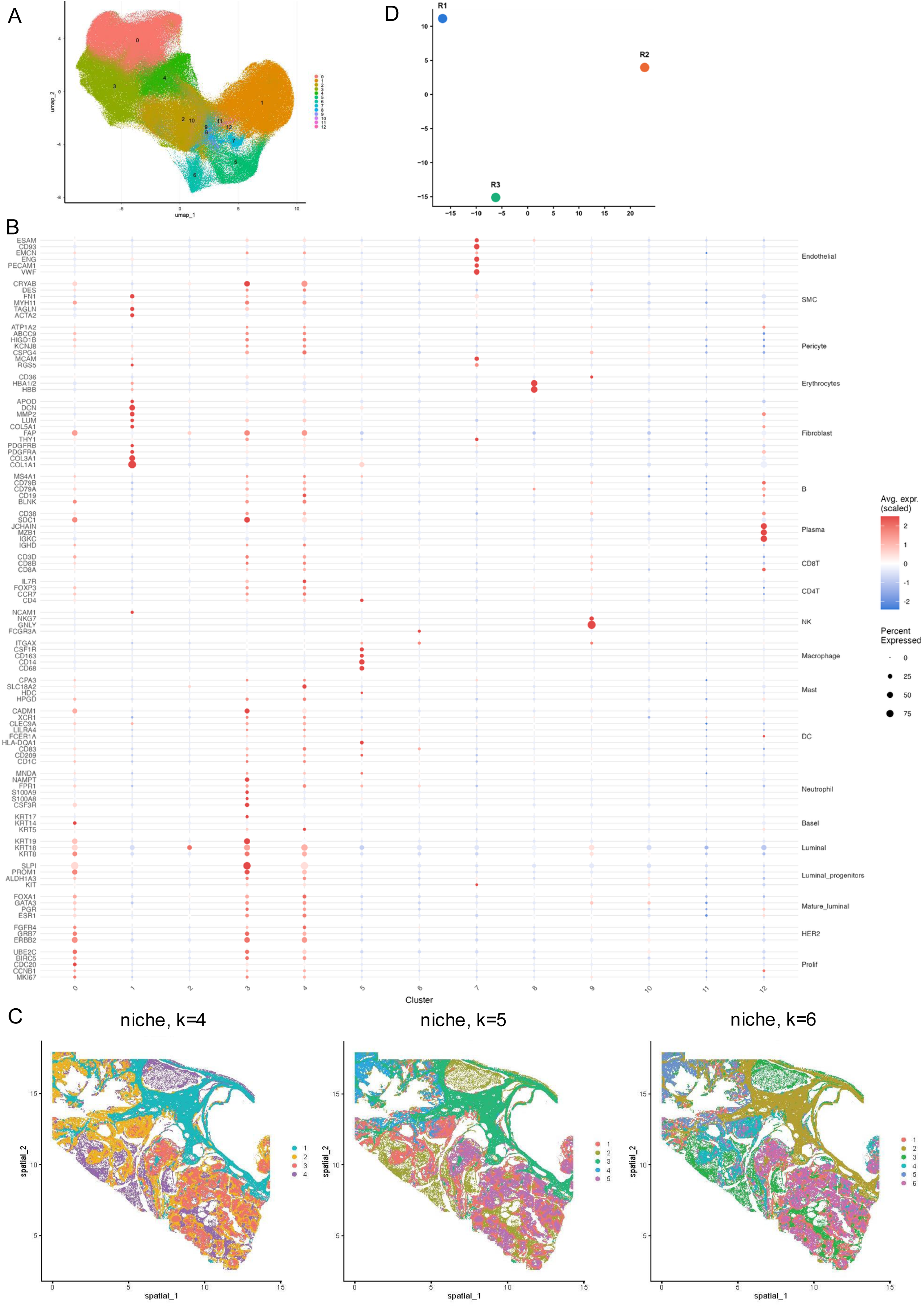
Analysis of breast cancer CosMx spatial transcriptomics data. **A.** UMAP embedding of CosMx spatial transcriptomics data with the original cluster labels **B.** Dot plot of known cell type specific markers for each cluster. **C**. Niche analysis of breast cancer spatial transcriptomics data with varying *k*-values **D.** PCA of region-level proteomic profiles from regions 1, 2, and 3.

**Supplementary Figure 21.**
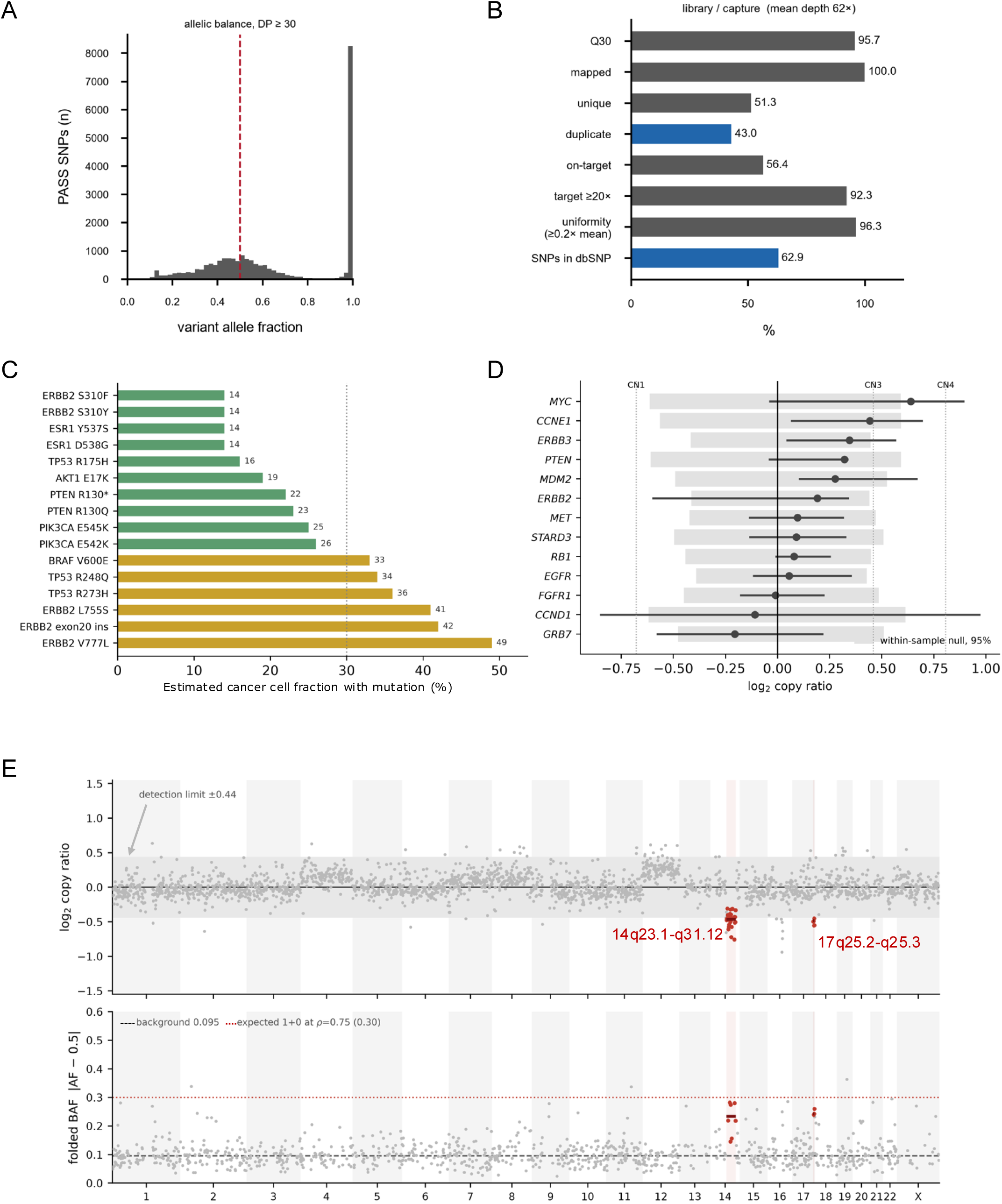
Whole exome sequencing of archived breast cancer sample compartment 1. **A.** Histogram of all SNPs detected as a function of VAF (VAF of 1.0 indicates homozygous SNP, and VAF of 0.5 indicates heterozygous SNPs). **B.** Sequencing quality control metrics breast cancer sample. **C.** Estimated cancer-cell fraction (CCF) containing indicated hotspot mutation given the observed depth with 95% power and ≥3 supporting reads. Hotspots where CCF is ≤30% are colored green. **D.** Log2 copy ratio (point, bootstrap 95% CI) for indicated genes against the 95% interval of a null built from size-matched contiguous windows else in the genome, which absorbs spatially correlated capture bias. No gene clears the null (minimum q = 0.51), and the upper bound for *ERBB2* is CN 3.0. **E.** Upper: GC-corrected log2 copy ratio relative to the on-target median for each chromosome. The grey shading shows the copy-number detection limit established from a within-sample contiguous-window null. Lower: Median folded B-allele frequency for each chromosome with a background of 0.095 (black dashed line). With a histology-estimated tumor purity of 0.7-0.8 (the percent of tumor cells in the biopsy), the expected compartment is 1+0 loss of heterozygosity (LoH) is 0.30 (red dotted line). The discovered LoH regions 17q25.2-25.3 and 14q23.1-q31.12 correspond to a CCF of 76% and 73% (by log2) or 93 and 91% (by folded BAF).

**Method Figure 1.**
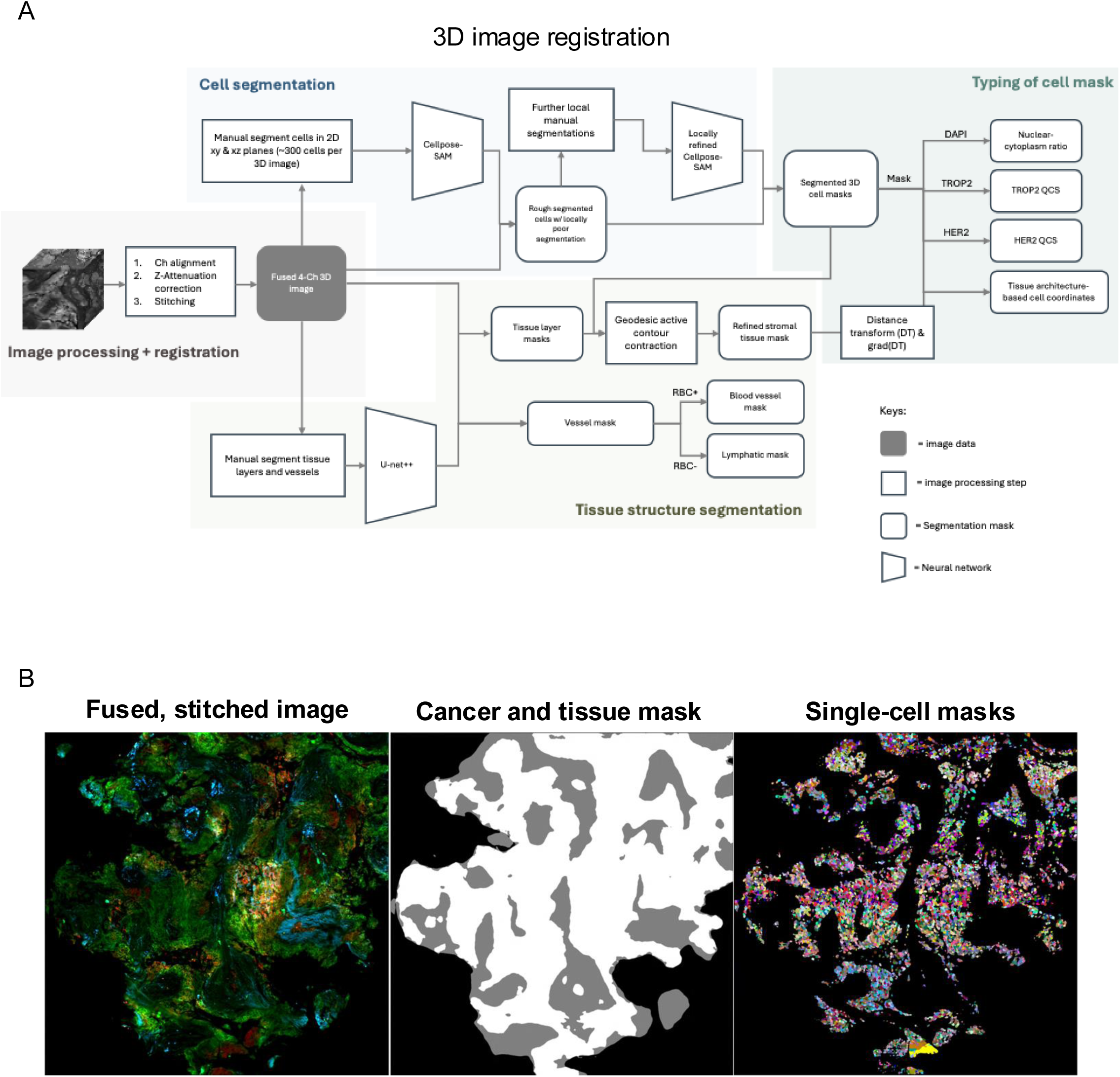
Computational pipeline for processing of lung cancer biopsy specimen images. **A.** Pipeline schematic. **B.** Illustrated processing results for fused and stitched bi-directionally illuminated LSFM image (left), segmented cancer regions (in grey) and tissue stroma (in white) (middle image), and segmented single-cell (color-coded by cell IDs) (right image).

**Method Figure 2.**
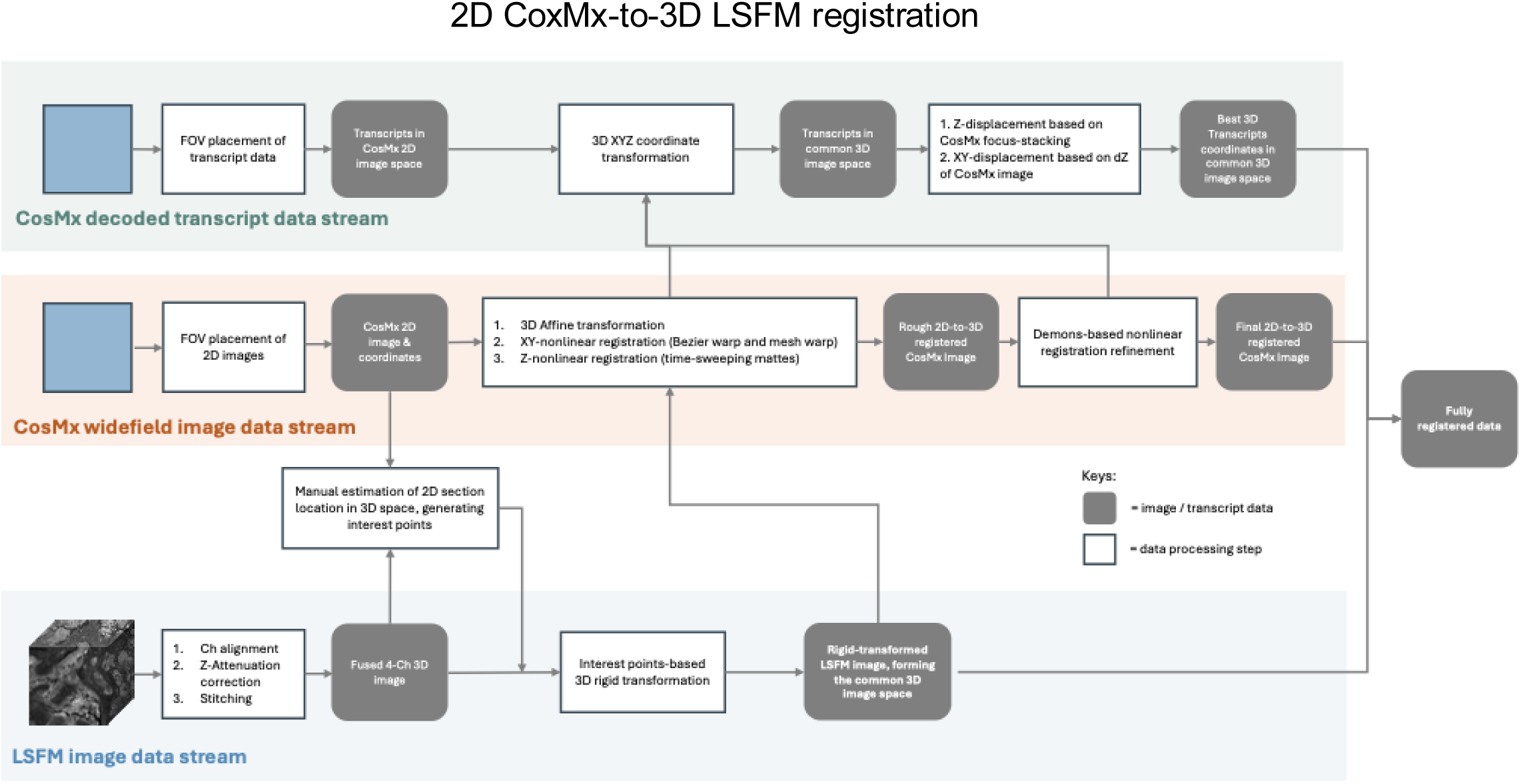
Computational pipeline for 2D CosMx-to-3D LSFM image registration and transcript coordinate transformation. Details were described in Methods.

## Supplementary Tables

**Supplementary Table 1.** Validated REZCUE-compatible antibodies.

**Supplementary Table 2.** RNA sequencing metrics of a REZCUE-treated sample versus its paired control FFPE sample (contralateral hemi-brain of the same mouse).

**Supplementary Table 3.** DNA extraction yields from 10 paired FFPE human lung cancer samples. For each patient ID, the on-block punch biopsies were homogenized mechanically and divided into two equal-weighted portions. One portion was REZCUE-treated and subsequently re-embedded as FFPE tissue; the other was kept at room temperature as a control.

**Supplementary Table 4.** Whole genome sequencing metrics of paired samples from 10 human lung cancer samples from Supplementary Table 3. Abbreviations: C, control; T, REZCUE-treated test sample; Ti/Tv, transition/transversion ratio; SNPs, single nucleotide polymorphisms; InDels, insertions and deletions; CNVs, copy number variations.

**Supplementary Table 5.** miRNAs found in the breast cancer sample that had been archived for 25 years.

## Materials and Methods

### Light-sheet fluorescent microscopy and confocal microscopy

LSFM was performed on a custom-built hybrid benchtop-mainframe mesoSPIM equipped with a zoom body and five lasers as previously described ^54^. Confocal microscopy was performed using a Confocal.NL NL5+ microscope with a 30× UPLSAPO30XS silicone oil-immersion objective (NA 1.05) with 405, 488, 561 and 640 nm lasers, and a Leica SP8 Stellaris confocal microscope with the objectives HC PL APO 10x / 0.40 CS2, HC PL APO 20x / 0.75 CS2, HC PL APO 40x / 1.30 Oil CS2 and HC PL APO 63x / 1.40 OIL CS2 and a white-light laser.

### Chemicals and reagents

Sodium dodecahydro-*closo*-dodecaborate was purchased from Katchem (Cat. no. 257) and used without further purification. 2-Hydroxypropyl-γ-cyclodextrin was obtained from Cyclolab (Cat. no. 38), Cyclodextrin Shop (Cat. no. CDexG-075), or Sigma Aldrich (Cat. no. 779229). Fluorophore-conjugated secondary antibody Fab fragments were purchased from Jackson Immunoresearch.

### Tissues

All experimental procedures were approved by the Animal Research Ethics Committee of the Chinese University of Hong Kong (CUHK) and were performed in accordance with the Guide for the Care and Use of Laboratory Animals. Adult male C57BL/6 mice were utilized. These mice were housed in a temperature-controlled environment (22-23 °C) at <70% relative humidity with a 12-hour light-dark cycle and unrestricted access to a standard mouse diet and water in the Laboratory Animal Service Center of CUHK. Tissues were perfusion formaldehyde-fixed and collected by post-mortem dissection. The research was approved by the CUHK ethics committee (AEEC number 20-287-MIS).

Human tissue samples were obtained from expired clinical blocks slated for disposal. These materials were cleared for discard due to inventory surpluses, extended archival duration, or the demise of the source patients. Ethics approvals were obtained prior to the study from the Joint Chinese University of Hong Kong-New Territories East Cluster Clinical Research Ethics Committee (approval number 2023.407), the Research Ethics Committee (Kowloon Central / Kowloon East) (approval number KC/KE-23-0146/ER-2), and the University of Hong Kong / Hospital Authority Hong Kong West Cluster Institutional Review Board (approval number UW 24-291).

### Tissue paraffin embedding

FFPE tissues from clinical sources were used as received. For tissues not obtained in FFPE format (such as fresh-fixed), the tissues were processed in an Epredia Excelsior AS Tissue Processor. The program involves tissue dehydration in 70% and 90% v/v ethanol in water for 30 minutes each, then in 100% ethanol 3 times for 45 minutes each, and finally in xylene 3 times for 60 minutes. The tissue was then placed in a 65 °C hot bath of histological embedding paraffin wax (Leica Surgiplast Paraplast Plus) for at least 18 hours in a Thermo HistoStar paraffin embedding station before embedding in acetal copolymer tissue cassettes.

### REZCUE processing

In brief, FFPE tissues were first cut out from the cassette, taking care not to crack the dried tissue. The tissue was then immersed in methylcyclopentane, which serves to quickly dissolve the majority of the paraffin, and then incubated in cyclohexane to ensure complete deparaffinization. The tissue was then immersed in 85:15 v/v THF:methanol solution to bridge towards gradient methanol-based rehydration. The rehydrated tissue was then treated in an aqueous solution containing 0.15 M *n*-hexadecyl-*N*,*N*-dimethyl-3-ammonio-1-propanesulfonate (zwittergent SB3-16, CAS number 2281-11-0), 0.25 M sodium dodecahydro-closo-dodecaborate (Na_2_B_12_H_12_), and 0.05 M alpha-cyclodextrin at 55 ॰C for up to 48 hours. The tissue was then washed in water, then in 50% v/v DMSO in water, and then in water. A detailed step-by-step protocol is given in **Supplementary Information**, along with photographs of the tissue after each stage.

### INSIHGT2

INSIHGT2 is an improved version of INSIHGT ^51^, which is a 3D tissue-clearing technique. INSIHGT2 is directly compatible with IgG secondary antibodies in a single unified protocol. Compared to the original version, INSIHGT2 splits the incubation of primary antibodies and secondary Fab fragments or whole IgG secondary antibodies into two separate steps, where both incubations utilize the same in situ host-guest chemistry between [B_12_H_12_]^2-^ and 2-hydroxypropyl-γ-cyclodextrin. This two-stage approach greatly enhances penetration compared to the original one-stage approach. Although the incubation time was slightly longer (fresh formalin-fixed tissue-to-image time of 6 days increased to 9 days), we deemed this adjustment acceptable for better performance. A detailed step-by-step protocol is given in **Supplementary Information**.

### Signal-to-background quantitative comparison

Human FFPE liver samples of various archival years (1, 10, and 30 years) were divided and subjected to xylene + HIER antigen retrieval versus REZCUE antigen retrieval. They were then both stained using INSIHGT2 for GLUL, along with tomato lectin (*L. esculentum* lectin, LEL) and DAPI. After clearing with BABB, confocal images were acquired at 100-120 µm depth from the tissue surface using a 40 × objective, targeting peri-central vein regions where GLUL signals should be present. Hepatocytes were then segmented using a custom-refined Cellpose-SAM model using the LEL channel for hepatocyte membranes, and the 3 cell layers surrounding the central veins were regarded as GLUL+ cells. Cells that are 9 cell-layers away from the central veins were regarded as anatomical GLUL-cells as background controls. The mean voxel intensity for GLUL was then computed for each peri-central hepatocyte and then divided by the mean voxel intensity of the GLUL-cells to obtain the per-cell GLUL signal-to-background ratio (SBR). Specimen-level SBRs were obtained by averaging all per-cell GLUL SBRs. The specimen-level GLUL SBRs were pooled and compared across the two treatment categories as geometric means. A two-tailed, two-sample Student’s *t*-test was used for null hypothesis testing.

### Antibody validation

FFPE samples were screened from the archives and processed through the REZCUE protocol. Tissues known to express the marker or normal human tissues in which the marker is constitutively expressed according to the literature and the Human Protein Atlas were used. The REZCUE-treated sample was stained using the INSIHGT2 protocol with DAPI counterstain and cleared using 1:2 v/v benzyl alcohol:benzyl benzoate (BABB) method. The samples were then imaged at 50-100 μm depths (due to working distance constraints) using a Leica Stellaris 8 Confocal microscope with an HC PL APO 63×/1.40 OIL CS2 (NA 1.40) objective, or a Confocal.NL NL5+ microscope with a 30× UPLSAPO30XS silicone oil-immersion objective (NA 1.05).

### Heat-induced epitope retrieval on intact FFPE tissue blocks

FFPE samples were carefully detached from the embedding cassette and deparaffinized in xylene at room temperature until no visible paraffin wax remains. The tissues were then rehydrated in graded ethanol (100% EtOH 30 min × 3, then sequentially in 90% EtOH, 70% EtOH, 50% EtOH, 30% EtOH, 30 min each), washed in water for 3 × 30 min, and antigen-retrieved in 10mM sodium citrate buffer, pH 6.0 at 95॰C for 1 hour. The tissue was then washed in water for 3 × 30 min and proceeded to INSIHGT2 immunostaining.

### Conventional 2D chromogenic immunohistochemistry

5 μm-thick FFPE sections obtained from REZCUE-treated, re-embedded tissues and its matched controls. The sections were cleared in xylene, rehydrated in graded ethanol, and antigen-retrieved with 10mM sodium citrate buffer at pH 6.0 for 5 minutes. Primary antibodies, horseradish-conjugated secondary antibodies (Jackson Immunoresearch 715-035-151 for mouse and Jackson Immunoresearch 711-035-152 for rabbit) were applied at 1:1000 dilutions sequentially. 1mg/mL 3,3’-diaminobenzidine (DAB) with 0.02% w/v hydrogen peroxide was then applied as a chromogen. The slides were further counterstained with hematoxylin and imaged on a Zeiss Digital Slide Scanner Axioscan 7.

### Bulk RNA sequencing analysis for mouse brain

An adult C57/BL56J mouse was perfused and fixed using ice-cold 4% PFA, and the brain was rapidly dissected, post-fixed in 4% PFA overnight at 4 °C, washed in PBS, dehydrated through graded ethanol series, cleared in xylene, and dissected into two halves that were separately embedded in paraffin wax. One hemisphere was antigen-retrieved with REZCUE and the control FFPE hemibrain was kept at room temperature protected from light. The REZCUE-treated hemibrain was then re-embedded in paraffin wax and sent to BGI for whole transcriptome RNA sequencing (including lncRNA and with rRNA depletion). To evaluate transcript coverage breadth, coordinate-sorted, indexed BAM files for the control and REZCUE-treated libraries were downsampled using *samtools view -s* to match the shallower library’s primary alignment depth based on samtools idxstats counts. Gene models were parsed from the GCF_000001635.27_GRCm39 reference genome, discarding genes shorter than 200 bp or lacking exon annotations; pre-mRNA intervals were defined by the full gene span, and mRNA intervals represented the merged union of overlapping exons. Covered bases (depth ≥ 1) were quantified simultaneously for both libraries using *samtools bedcov -d 1 -j*, ensuring spliced gaps and deletions were excluded from coverage calculations. Percent coverage breadth was calculated for each gene as covered bases divided by interval length, retaining only genes with at least one covered base in either condition. Finally, a paired Wilcoxon signed-rank test was applied to compare control and REZCUE-treated data breadth values across retained genes for each interval type, reporting per-library medians and the probability that the REZCUE-treated sample exceeds the control to describe the gene-level shift between the single-replicate libraries.

### Mass spectrometry-based proteomics analysis of mouse brain

The mouse brain samples were further sectioned to yield three slides containing paired REZCUE-treated and control sections. The sections were then sent to Climb Technology for laser microdissection, protein extraction, trypsin digestion, and mass spectrometry-based proteomics analysis by data-independent acquisition. Briefly, each slide was first stained with H&E. Laser-microdissection was then performed around the cerebellum (chosen for its molecular homogeneity and well-defined anatomy) using the MMI system. The collected tissue regions were then lysed, heat-denatured, sonicated, and digested overnight with trypsin (0.5 μg/μL) at 37 °C in 50 mM TEAB, followed by SDB-RPS desalting and vacuum drying. Dried peptides were resuspended in 0.1% formic acid and separated on a 25 cm Aurora C18 column (75 µm ID, 1.6 µm beads; IonOpticks) at 50 °C using a nano-UHPLC system (Thermo Fisher) coupled to a timsTOF Pro2 (Bruker Daltonics). Separation was performed at 300 nL/min over a ∼31-minute gradient (6–99% acetonitrile in 0.1% formic acid), and data were acquired in dia-PASEF mode (MS range: 100–1700 m/z; ion mobility: 0.6–1.6 Vs/cm²; collision energy: 20–59 eV; 32 windows; capillary voltage: 1600 V). Raw data were processed with Spectronaut 20 (Biognosys) against the UniProt mouse proteome using trypsin digestion with up to 2 missed cleavages, fixed carbamidomethylation of cysteine, variable oxidation of methionine and N-terminal acetylation, and precursor/fragment mass tolerances of 15/20 ppm. Peptide and protein false discovery rate thresholds were both set at < 1%, with a minimum of one unique peptide required per protein identification.

### Whole genome sequencing comparative analyses using human lung samples

Ten formalin-fixed paraffin-embedded (FFPE) blocks of human lung tissues were sampled via a single punch, homogenized, and divided into equal portions. One portion was REZCUE-treated and then re-embedded into paraffin wax, while the untreated portion served as a control and was stored under ambient conditions (15-25 °C, humidity 50-70%, in the dark) all along. These paired samples were then sent to BGI Genomics for DNA extraction and quantification as quality control before proceeding to sequencing. Five pairs selected based on yield exceeding input thresholds in both arms were sequenced. DNA yields for these sequenced pairs ranged from 2.6 to 12.0 µg in the lower-yielding arm, compared to 0.28 to 1.56 µg in the remaining five non-sequenced pairs. REZCUE-treated sample DNA yields were compared against a post-hoc non-inferiority margin of -20% of their paired controls by the exact one-sided Wilcoxon signed-rank test. Libraries were prepared using Short-Insert Library (utilizing KAPA HiFi HotStart DNA Polymerase for PCR amplification), sequenced on the DNBSEQ (specifically paired-end 150 bp, PE150) platform to 15–45× depth per sample, aligned to GRCh38 using BWA-MEM (v0.7.17), and duplicate-marked. Variants were called per sample using GATK HaplotypeCaller (v4.4.0.0) without joint genotyping. Because split portions of each pair were genetically identical, variants shared between halves were classified as reproducible, while private variants were considered irreproducible artifacts. No external truth set or matched normal tissue was available; thus, variants unique to a single arm were not explicitly verified as false positives. We analyzed the retention of reproducible variants after an identical filter applied to both arms, which restricted analysis to sites with 20–50× depth in both arms to control for coverage differences, fraction of the genome callable at ≥20×, and the irreproducible call fraction. Each endpoint was evaluated using an exact one-sided Wilcoxon signed-rank test on paired log-ratios (for multiplicative endpoints) or paired differences (for additive endpoints) against post hoc non-inferiority margins (−10% retention or −5% breadth). At n = 5, the exact minimum one-sided *p* value is 0.0313, which is attainable only when all five pairs surpass the non-inferiority margin; the test effectively reduces to a sign test where magnitudes contribute solely to the interval estimate. The corresponding two-sided minimum *p* value is 0.0625, and superiority was not claimed. Co-primary endpoints (i) and (ii) were combined using the intersection–union principle, requiring no multiplicity adjustment. Confidence intervals represent exact distribution-free bounds at the achieved coverage (93.75% at n = 5). Comprehensive metrics are provided in **Supplementary Table 4**.

### CosMx spatial transcriptomic comparative analysis of mouse brain tissue

#### Tissue processing

An adult C57BL/6 mouse was subjected to perfusion-fixation using ice-cold 4% PFA, and the brain was rapidly dissected, post-fixed in 4% PFA overnight at 4 °C, and washed in PBS. The brain was then bisected, both embedded in paraffin wax and one of the hemispheres was retrieved for REZCUE treatment, while the other half was kept as a control at room temperature (15-25 °C, humidity 50-70%, in the dark), mimicking routine FFPE storage conditions. For the REZCUE-treated hemisphere, the tissue was melted out of the paraffin wax, REZCUE-treated, then stained with the INSIHGT2 method for 3D triplex immunostaining ^51^, then cleared using the BABB method, imaged with a mesoSPIM (v5.1) light-sheet microscope, washed in 100% methanol, cleared with xylene, infiltrated with molten paraffin wax, and re-embedded as a paraffin block. The REZCUE-treated and control hemispheres were then sectioned using a microtome, taking care during mounting to ensure subsequent sectioning will obtain sections of similar depth and regions from both hemispheres. The paired sections were then placed on the same slides and were run on the same microfluidic cell in the Nanostring CosMx single-molecule imaging platform.

#### Quality control

Single-cell spatial gene expression data from mouse brain tissue sections acquired on the CosMx platform were analyzed using Scanpy (v*1.9.6*; ^55^) in Python. Counts from negative control probes included on the CosMx panel were used to estimate background signal as a fraction of total counts per cell. Quality-control metrics, including total transcript counts and unique genes detected per cell, and total counts per FOV, were calculated and inspected across cells and FOVs, including a comparison of the FOV-level transcript-count distribution between REZCUE-processed and control sections. The spatial distribution of total transcript counts per cell was additionally visualized using each cell’s spatial centroid coordinates.

#### Preprocessing, clustering, and batch integration

Low-quality cells (<100 total counts) and genes detected in <400 cells were removed. Raw counts were preserved, and the remaining data were library-size normalized and log-transformed. PCA, a neighborhood graph, and a UMAP embedding were computed on the full dataset, and cells were clustered using the Leiden algorithm. Cluster marker genes were identified using a Wilcoxon rank-sum test. For the REZCUE vs. control comparison, the S1 and S2 sections were subsetted, and technical variation between sections was corrected on the PCA embedding using Harmony ^56^, with tissue-section identity (REZCUE vs. control) specified as the integration covariate. A neighborhood graph, UMAP embedding, and Leiden clustering were recomputed on the Harmony-corrected principal components.

#### Cell-type annotation

Cluster identities were assigned to cell types by cross-referencing the top differentially expressed genes per cluster (Wilcoxon rank-sum test, top 50 genes per cluster) with canonical markers of mouse brain cell types, including *Slc17a7*, *Camk2a*, *Nrgn*, and *Satb2* for glutamatergic neurons, *Gad1*, *Gad2*, *Slc32a1*, and *Pcp4* for GABAergic neurons, *Pvalb* and *Kcnc1* for Pvalb^+^ interneurons, *Mbp*, *Plp1*, *Mobp*, and *Mag* for oligodendrocytes, *Apoe*, *Aldoc*, *Slc6a11*, and *Aqp4* for astrocytes, *C1qb*, *Ctss*, *Sparc*, *Gsn* for microglia, *Col1a1*, *Col1a2*, *Lum*, and *Dcn* for vascular and leptomeningeal cells, and *Ttr*, *Clic6*, *Igfbp2*, and *Krt18* for choroid plexus epithelial cells. This yielded 15 annotated clusters, including neuronal subtypes distinguished by regional or molecular markers (e.g., *Tcf7l2*^+^ thalamic excitatory neurons and *Pcp4*^+^/*Tac1*^+^ GABAergic neuron subtypes). Annotation quality was confirmed by dot plots of the curated canonical marker panel and of the top data-driven cluster markers across annotated cell types.

#### Spatial mapping and compositional analysis

Cell-type identities were mapped onto tissue sections using each cell’s spatial centroid coordinates. To compare cell-type composition between conditions, the fraction of cells assigned to each cell type was calculated within each condition, and the difference in fraction between REZCUE-processed and control samples was computed for each cell type and visualized as a dot plot. This compositional comparison was descriptive in nature; no formal statistical test of differential abundance was applied.

### 3D LSFM image processing: channel alignment, stitching, and illumination fusion

For LSFM, each sample was imaged with dual-sided light sheet illumination to maximize depth coverage with the full specimen volume acquired as a series of overlapping tiled z-stacks collected sequentially channel by channel. This per-channel acquisition strategy introduces inter-channel positional offsets: At imaging resolutions below 1 µm/pixel, the limited mechanical repeatability of long-travel motorized translation stages causes small but significant lateral displacements between channels within each tile. These offsets were corrected by manual channel alignment in Adobe After Effects prior to further processing.

Tile stitching required two additional corrections. Non-uniform Z-step intervals across tiles, caused by stage mechanics, introduced depth-dependent positional inconsistencies, and lateral sample drift during acquisition produced position-dependent XY offsets between tiles. Both effects were corrected in After Effects using keyframe-based parameter adjustment, a technique in which correction values are explicitly specified at discrete Z-positions (keyframes) along the timeline and interpolated between them, allowing spatially varying corrections to be applied continuously across the volume depth. Depth-dependent axial corrections used optical flow–based keyframe interpolation, a pixel-motion estimation method that computes spatially varying displacement fields between frames to achieve smooth, continuous transitions, while lateral (XY) corrections used linear keyframe interpolation.

The stitched volumes from the two opposing illumination directions were then fused. Imperfect cuvette flatness, residual rotations, and inter-stack misalignment result in slightly different spatial orientations of the two illumination-view volumes. To correct this, the central ∼50% overlapping region of both stacks was used to identify corresponding landmark points, from which a 3D affine transformation **T**_aff_(**x**)=**Ax**+**b**, with voxel coordinate **x** ∈ ℝ^3^, linear map **A** ∈ ℝ^3×3^ encoding rotation, scaling, and shear, and translation vector **b** ∈ ℝ^3^, was estimated in MATLAB and applied to one of the two illumination-view stacks. The transformed stack was re-imported into Adobe After Effects, where depth-dependent residual misalignments were further corrected with keyframe-based adjustments to achieve sub-cellular registration between views. The two aligned stacks were blended using a spatially feathered mask centered at their mid-overlap boundary. This is a soft-edge blending approach in which mask influence attenuates linearly near the boundary, equivalent to linearly interpolating intensities across the overlap zone, yielding the final channel-aligned, stitched, and fused 3D LSFM image stack.

### Image processing

No depth-varying nor non-linear intensity transformations were performed on any images in this study. To remove confocal imaging noise, a Gaussian filter of sigma 1 was applied to confocal images when necessary. To remove strip artefacts in the line-scan confocal images, the frequency domain image was obtained by 2D fast Fourier transform (FFT), and a vertical mask centered on the u-axis of the frequency domain image was applied. An inverse 2D FFT was then applied to obtain the artefact-filtered image. All image filtering operations were performed in ImageJ.

### Image segmentation

Segmentation of anatomical structures and cells were performed using multiple tools and custom scripts, as decided by the characteristics of the structures to be segmented. In brief, cells were segmented using a custom retrained Cellpose-SAM model ^35^. Anatomical compartments such as skin appendages, eccrine sweat glands, epidermis were segmented using a custom trained U-Net++ neural network with an encoder depth of 7^57^. Training images were obtained by taking every 20 images along the Z-stack and manually segmented using the 3DSlicer or Ilastik, after which the image and segmentation masks were downsized to 1024 × 1024 pixels and used to train the U-net++ network, implemented in MATLAB ^58,59^. For vascular structures such as capillaries and lymphatic vessels, a coherence enhancing diffusion filter was applied to bridge bright, discontinuous signals into a smooth interconnected component. Additional 3D structure tensor images were also computed to augment the number of channels per slice to help the neural network gain semi-3D information on each slice. The resulting combined multi-channel image set, along with the manual segmentation masks, were then fed into a separate U-net++ network for training. Inferences by the neural network were performed on 2×-downsized images.

### CosMx image preparation for 2D-to-3D registration

The CosMx dataset was collected from a series of FFPE sections cut from a tissue block that had previously been processed through REZCUE, immunostaining, optical clearing, LSFM imaging, clearing reversal, and re-embedding. Sections were arranged on the 20 × 15 mm CosMx scan area following the manufacturer’s slide preparation guidelines. Raw output data from the CosMx run were organized by FOV. A compressed coordinate file recorded the pixel- and micrometer-scale position of each FOV on the slide; the Morphology2D directory contained multichannel fluorescence images for each FOV; and each tile contained decoded transcript coordinates with their associated gene identifiers.

All FOVs were assembled into a unified 2D full-slide coordinate system using a custom MATLAB script, in which each FOV’s image data and transcript coordinates were offset by the corresponding FOV stage position, yielding a continuous mosaic spanning more than 110,000 × 110,000 pixels. Individual tissue sections were then manually cropped from this mosaic, and section-local coordinate systems were defined with the origin at the upper-left corner of each cropped region; transcript coordinates were offset accordingly.

To enable subsequent non-linear coordinate transformation, two reference coordinate grid images were generated for each section: one in which pixel intensity encodes the pixel’s row position, and one encoding the column position, each with uniformly spaced intensity increments. Together, these two images constitute a forward coordinate map of the original section frame, enabling recovery of the spatial transformation applied to any pixel location. A binary crop mask was generated alongside. All three images were stored as separate channels of a single RGB image to allow simultaneous propagation through all downstream transformations. CosMx native cell segmentation outputs were processed identically. All FOV-level segmentation masks were assembled into the full-slide coordinate system, then cropped and coordinate-transformed to their respective section-local frames.

### Aligning 3D LSFM images and CosMx spatial transcriptomics data

To align 3D LSFM images with CosMx spatial transcriptomics, near-voxel-perfect co-registration of 2D tissue sections onto a 3D volume must be achieved in the presence of tissue cracks, sectioning artifacts, non-rigid section deformations, and optical aberrations; these features render fully automated registration intractable. We used Adobe After Effects video compositing and motion graphics software as a platform for human-in-the-loop image alignment and registration.

A central feature exploited throughout this workflow was the dimensional equivalence between the Z-axis of an isotropic 3D LSFM volume and the temporal (frame) axis of the After Effects project timeline. By loading the 3D LSFM volume as a time series, each Z-plane of the volume corresponds to a single video frame. This Z-to-time mapping allows 3D spatial registration problems to be reformulated as temporal alignment operations. This allowed us to take advantage of the GPU-accelerated rendering, efficient memory management, real-time preview, and pre-computed frame caching implemented in After Effects, all of which are essential for interactive, sub-cellular-precision manipulation of large microscopy volumes. One inherent limitation of After Effects is that rotational adjustments in the image plane are not as readily integrated into the interactive pipeline as lateral or axial adjustments. Rotational corrections were therefore performed externally in MATLAB using custom scripts, based on corresponding control points identified interactively within the After Effects interface. A comprehensive description of the After Effects-based workflow will be provided in a dedicated methods article; the complete image analysis pipeline is summarized in the following sections.

#### 2D CosMx section-to-3D LSFM volume registration

Registration of each 2D CosMx tissue section to the 3D LSFM volume was performed in three sequential stages: (1) a global 3D rigid-body alignment of the LSFM volume to the tissue sectioning plane, (2) manual non-linear deformation of the 2D section image in the lateral (XY) plane and along the axial (Z) direction within After Effects, and (3) voxel-level refinement of the lateral registration using the demons algorithm in MATLAB. The full method of 2D-to-3D registration, including a step-by-step video tutorial.

#### Global 3D rigid-body alignment

Each cropped section image was imported into After Effects to determine its approximate Z-position in the LSFM volume and to obtain initial estimates of in-plane rotation, scaling, and translation. Because tissue blocks are rarely embedded in perfect alignment with the LSFM imaging axes, the physical sectioning plane is typically tilted relative to the XY image plane of the LSFM volume: A nominally flat tissue section therefore spans a range of Z-depths within the 3D volume. To correct for this tilt, corresponding landmark points were identified across the range of Z-values spanned by a randomly selected section on the 3D volume, and a 3D rigid-body transformation, **T**_rigid_(**x**)=**Rx** + **t** with rotation matrix **R** ∈ SO(3) and translation vector **t** ∈ ℝ^3^, was fitted to the landmark correspondences in MATLAB and applied to the entire 3D LSFM volume. This reorients the LSFM volume so that its XY-plane is parallel to the tissue sectioning plane. The rigid-transformed LSFM volume defines the global 3D coordinate system to which all subsequent image data and transcript coordinates are referenced.

#### Manual non-linear registration in After Effects

The rigid-transformed LSFM volume was re-imported into After Effects for non-linear registration, which proceeded in two sub-steps: lateral (XY) deformation followed by axial (Z) displacement mapping. For lateral deformation, a suite of local spatial warping effects, specifically CC Bend It, Bézier Warp, and Mesh Warp, each of which deforms an image region according to user-positioned control points, was applied to the 2D section image to match it to the LSFM volume at a target Z-position. To avoid registering the section to an incorrect depth, each control point adjustment was validated by examining match quality across a range of neighboring Z-positions, confirming that the deformation yields the best correspondence at a unique, well-defined depth. Deformations were also constrained to remain smooth and of moderate magnitude, consistent with the expectation that biological tissue deformations are approximately diffeomorphic (i.e., smooth and invertible). No crossing of grid lines was allowed during Mesh Warp application.

For axial displacement mapping, we exploited the Z-to-time dimensional mapping of the After Effects timeline to encode the spatially varying depth of the tissue section. Because different regions of a physically non-flat tissue section lie at different Z-depths in the 3D LSFM volume, registering the full section requires specifying, for each pixel of the 2D section image, the Z-depth at which that pixel best matches the LSFM data. This information was encoded in a Z-displacement map **M** : Ω → [0, 65535], a 16-bit grayscale image defined over the 2D section domain Ω, in which the luminance value at each pixel linearly encodes the local axial position of that tissue region: luminance 65535 (white) corresponds to the Z-position closest to the LSFM camera, and luminance 0 (black) to the farthest position. The Z-displacement map was constructed within After Effects using the Fill effect, which assigns a uniform luminance value to a user-drawn mask region, and then applied to a stack of hand-drawn masks delineating tissue regions with distinct axial positions (referred to in compositing terminology as a matte: a grayscale mask image used to selectively control the visibility of a layer). Edge feathering was applied at each mask boundary, producing smooth, spatially continuous transitions between regions of different Z-displacement, equivalent to applying Gaussian blurring to the displacement magnitude map.

To reconstruct the 3D position of the section from this map within After Effects, the Extract effect was used. The Extract effect is a compositing operation that renders only those pixels of an input layer whose corresponding luminance values in a reference matte fall within a specified range [L_low_, L_high_], masking all other pixels transparent. A JavaScript expression was written to automate the luminance window sweep. The window descends linearly from 65535 (white) at the first frame to 0 (black) at the final frame. Because each frame corresponds to a distinct Z-plane, this sweep progressively reveals each tissue region at its encoded Z-depth. At any given frame, only the tissue region whose matte luminance matches the current window is rendered, effectively placing that region at the correct axial position in the 3D volume. The temporal span of the layer was matched to the Z-span of the section in the LSFM volume. This architecture allows the XY deformation, Z-displacement magnitudes, and spatial boundaries of each Z-displacement region to be adjusted interactively at any time, with all changes immediately visible through After Effects’ real-time GPU-accelerated rendering pipeline.

To account for the finite physical thickness of each tissue section (∼6.4 µm) relative to the 1 µm axial sampling interval of the LSFM acquisition, a small luminance boundary softness parameter was applied to the Extract effect. This broadens the luminance window slightly, allowing each tissue region to contribute to a narrow range of Z-positions rather than a single plane, equivalent to extending the effective depth-of-field of the section during compositing, and physically consistent with the capacity of a 6.4 µm-thick section to match LSFM data across several adjacent Z-planes through slight defocusing.

#### Voxel-level refinement with the demons algorithm

The Z-displacement maps and XY-deformed section images were exported from After Effects and imported into MATLAB for voxel-level refinement. For each Z-level defined by the displacement map, the corresponding portion of the XY-deformed section image was obtained by multiplying the deformed image by the binary Z-level mask derived from the displacement map at that depth, and then registered to the corresponding Z-plane of the rigid-transformed LSFM volume using the MATLAB demons algorithm, an iterative non-rigid image registration method that treats intensity differences between a moving and fixed image as forces driving a dense voxel-wise displacement field toward convergence. The resulting displacement field was saved for each Z-level. This procedure was repeated across all Z-levels spanned by each section and for all sections, yielding a complete, voxel-accurate non-rigid registration of each 2D CosMx section to the 3D LSFM volume.

#### Transcript coordinate transformation

Following image registration, the spatial coordinates of all decoded transcripts were subjected to the same sequence of transformations as the image data, ensuring precise spatial correspondence between transcript positions and the registered images for subsequent single-cell spatial analysis. To propagate the non-linear transformations to point coordinates rather than image pixels, the reference coordinate grid images generated during CosMx data preparation served as fiducial trackers. The two-channel coordinate map, in which pixel intensities encode the original row and column positions, was passed through the identical After Effects processing pipeline. The MATLAB demons algorithm was applied to the corresponding section image, with two critical constraints: Nearest-neighbor interpolation was used throughout, and all anti-aliasing was disabled, as anti-aliasing would blend adjacent encoded coordinate values and corrupt the mapping. By comparing the pre- and post-transformation coordinate maps pixel-by-pixel, an interpolated forward transformation function was derived for each section, mapping any 2D position (*u*, *v*) in the original section coordinate system to its transformed counterpart.

The axial coordinate of each transcript was assigned by combining three contributions: its native within-section Z-position as measured by CosMx through Z-focus stacking, the nominal Z-depth of the section in the global LSFM coordinate system, and the local Z-displacement encoded in the matte at the transcript’s lateral position. Finally, to account for the lateral parallax introduced by spatially varying Z-displacement, regions with non-zero displacement gradients were geometrically shifted laterally relative to a planar reference. The spatial gradient of the Z-displacement map was computed, and the resulting lateral correction vector was applied to each transcript’s (*u*, *v*) coordinates. The outcome is a complete table of 3D transcript coordinates registered to the global LSFM coordinate system with sub-cellular precision in all three spatial dimensions, achieving fully 3D-registered spatial transcriptomics.

### CosMx spatial transcriptomic analysis of human breast tumor tissue

#### Data processing, dimensionality reduction, and clustering

Single-cell spatial gene expression data for the breast tumor specimen CA2 were acquired on the CosMx Spatial Molecular Imager platform and processed into cell-by-gene count matrices prior to the analyses described here. SCTransform-normalized and scaled expression data were imported into Seurat (v5.4.0 ^60^ ) as a Seurat object. PCA was performed, and a shared nearest-neighbor graph was constructed using the first 30 principal components (*FindNeighbors*, reduction = “pca”, dims = 1:30). Cells were clustered using the Louvain algorithm implemented in *FindClusters* across a range of resolutions. A resolution of 0.2, yielding 13 clusters, was selected for downstream cell-type annotation. 2D embeddings for visualization were generated by uniform manifold approximation and projection (dims = 1:30).

#### Cell-type annotation

Cluster marker genes were identified with *FindAllMarkers* (Wilcoxon rank-sum test, only.pos = TRUE, min.pct = 0.25). The top five markers per cluster, ranked by average log2 fold change, were considered together with a curated panel of canonical lineage markers spanning stromal (endothelial, smooth muscle/pericyte, fibroblast/cancer-associated fibroblast), immune (B, plasma, CD8^+^ T, CD4^+^ T, NK, macrophage, myeloid, mast, dendritic, neutrophil), and epithelial/tumor (basal, luminal, luminal progenitor, mature luminal, HER2, proliferative) lineages to guide cluster annotation. Mean scaled expression and the percentage of cells expressing each marker within each cluster were visualized as a dot plot (*DotPlot*, grouped by cluster, faceted by marker category). On the basis of these marker profiles, each of the 13 clusters was manually assigned to a cell type or tumor subtype (Tumor Luminal-Basal, Tumor Luminal A, Tumor Hypoxic, Tumor Luminal B-HER2^+^, cancer-associated fibroblasts, macrophage, myeloid, endothelial, erythrocyte, NK, plasma, or non-specific/low-quality).

#### Spatial visualization of cell types, compartments, and niches

Cell-type identities were visualized in UMAP space and mapped onto the tissue section using the x/y centroid coordinates extracted from the CosMx field-of-view image object. For spatial cell-type maps, points were rendered in a fixed layering order (background/non-specific cells first, followed by stromal and immune populations, and tumor subtypes last) to improve visibility of rarer tumor populations against the dense background. Compartment assignments and tissue niche assignments were mapped onto tissue coordinates and visualized as categorical spatial scatter plots.

#### Cell-type composition across compartments and niches

To quantify how cell-type composition varied across compartments and niches, cross-tabulations of cell type by compartment and cell type by niche were constructed, and proportions were computed within each compartment or niche. For each cell type, the range of variation across compartments was calculated and used to rank cell types by the magnitude of compositional change.

#### Identification of compartment-divergent genes

To identify genes whose expression diverged most strongly among malignant compartments, tumor cells assigned to the three primary compartments were extracted, and normalized expression values were retrieved as a sparse matrix. For each gene, mean normalized expression was calculated within each compartment and log2-transformed. For each gene, the “driver compartment” was defined as the compartment with the highest log2-transformed mean expression, and the magnitude of divergence was defined as the difference between the maximum and minimum log2-transformed mean expression across the three compartments. The five genes with the greatest log2(fold change) range within each driver compartment were selected as compartment-divergent marker genes.

### Mass spectrometry data analysis for human breast tumor tissue

#### Data processing and principal component analysis

Normalized protein intensities from three microdissected regions of the same tumor specimen (regions 1, 2, and 3) were imported, and proteins with non-missing (detected) values in at least one region were retained. Intensities were log2-transformed within each region. For PCA and pairwise distance calculations, the subset of proteins detected in all three regions was used. Log2 intensities were standardized (z-scored) within each region across all proteins, and PCA (n_components = 2) was applied to the region-by-protein z-scored matrix to summarize the overall proteomic relationships among the three regions. Pearson correlation coefficients and pairwise Euclidean distances between regions were also computed on the z-scored data.

#### Protein annotation

Protein identifiers were annotated with gene symbols, gene names, and full protein names by querying the UniProtKB REST API (rest.uniprot.org/uniprotkb/search); the primary gene symbol for each protein was taken as the first token of the returned gene-names field.

#### Trajectory-based identification of gradient proteins

To identify proteins whose abundance changed progressively along a defined spatial trajectory through the three regions, we applied a stepwise, trajectory-based statistic. For a given ordering of the three regions (e.g., 1 → 2 → 3), each protein’s log2 fold change over each of the two sequential “legs” of the trajectory (leg 1: region 2 vs. region 1; leg 2: region 3 vs. region 2) was computed, and each leg’s fold-change distribution was independently z-scored across all proteins to derive an empirical null distribution. A protein was considered to change monotonically along the trajectory if both legs changed in the same direction (matching sign). For monotonic proteins, an intersection–union test statistic was defined as the smaller of the two legs’ absolute z-scores (z_min_), which was converted to a two-sided empirical p-value using the standard normal cumulative distribution function [p = 2 × (1 − Φ(z_min_))]; non-monotonic proteins were assigned z_min_ = 0 (p = 1). Proteins were called trajectory “gradient” proteins (increasing or decreasing) if they were monotonic, showed an endpoint-to-endpoint log2 fold change ≥ 1 in absolute value, and had z_min_ ≥ 2 (i.e., both legs individually deviating from the null by ≥2 standard deviations in the same direction). This analysis was performed independently for each of three possible region orderings (1 → 2 → 3, 3 → 2 → 1, and 1 → 3 → 2) to evaluate all distinct trajectory hypotheses through the three regions. Results were visualized as volcano-style plots (endpoint-to-endpoint log2 fold change vs. −log10[p], capped at 20 for visualization, with capped points indicated separately), with gene symbols labeled for called gradient proteins using a custom label-repulsion algorithm to minimize label overlap.

### Small RNA sequencing analysis of human breast tumor tissue

Two punch biopsies were obtained over the compartment 1 region on the breast tumor tissue FFPE block. RNA from the biopsies was extracted and sent to BGI for small RNA sequencing. The two biopsies (designated as A and B) yielded 32.0M reads and 32.5M reads, with 12.7M and 11.2M uniquely attributable reads, respectively. To identify the miRNAs, reads were collapsed to distinct 15-60 nt sequences and matched to miRBase v22 mature tags under a one-credit-per-read rule: where each read was assigned to its single longest exact or 3’-extended match using an anchor of 18-34 nt, so no read contributes to more than one feature. Putative miRNAs were then filtered to exclude miRBase-documented structural RNAs (miR-1246 (RNU2-1/U2 snRNA), miR-1274a/b, miR-1280, miR-720, miR-1308 (tRNA fragments), miR-886 (nc886/VTRNA2-1), miR-664a/b (SNORA36), miR-1291 (SNORA34), miR-4488, miR-4516, miR-12136). Putative miRNAs with a modal tag matching an rRNA or tRNA, or with tags sharing with an Rfam family whose Rfam type is not “Gene; miRNA” were also excluded. Rfam accessions typed as miRNA precursor families were retained (RF00027, let-7, MIPF0000002; RF00736, mir-320, MIPF0000163); accessions typed as structural were rejected, including RF01684 (mascRNA/menRNA, the tRNA-like RNase-P product of MALAT1), RF01960 (18S rRNA), RF02543 (28S rRNA), RF00023 (tmRNA) and RF00005 (tRNA). Indistinguishable mature sequences were reported as single groups. Abundances are counts per million attributable reads; detection floors are 0.79 CPM (biopsy A) and 0.89 CPM (biopsy B). The miRNAs were then classified by counting the number of literature on human breast cancer in support of their roles being oncogenic or tumor suppressive, performed by looking up on the OncomiRDB (oncomirdb.v-1.1-20131217) and miRCancer databases (version June 2020). miRNAs with no data or ties are marked as ambiguous.

### Whole exome sequencing analysis of human breast tumor tissue

A third punch biopsy was obtained over the compartment 1 region after the initial two punch biopsies for small RNA sequencing. DNA from the punch biopsy was extracted, exome-captured, and sequenced by BGI. The sequences were aligned to GRCh38 (full analysis set with decoys) with BWA-MEM, duplicate-marked, and base-quality recalibrated. Somatic point mutations were called with Mutect2 against an allele-frequency-only germline resource and filtered with FilterMutectCalls including a read-orientation prior, which removed 651,384 of 1,002,222 candidates, consistent with formalin-induced cytosine deamination. Hotspot negatives were converted to excluded cancer-cell fractions by binomial power (≥3 supporting reads, 95%). Target depth was bimodal (off-target mode 6.5×, on-target mode 57.3×), so all copy-number analysis was restricted to capture bins exceeding the between-mode minimum (21.1×; 300,730 bins, 59.9 Mb) and expressed as log2 ratios to the on-target median (56.0×). GC bias was removed by subtracting the median log2 of 40 GC quantiles (per-bin |log2| median absolute deviation 0.412 → 0.395). Lacking a matched normal and a process-matched panel of normals, significance was assessed against a within-sample null: for each gene or segment, the distribution of median log2 across 2,000–4,000 size-matched contiguous bin-runs sampled from other chromosomes, which absorbs spatially correlated capture bias. Empirical *p* values were Benjamini–Hochberg adjusted (gene level) or reported one-sided (segment level). The copy number was derived as 2 + 2(2^log2^ − 1)/ρ with ρ = 0.75.

Allelic imbalance was assessed from tumor-only heterozygous SNVs with depth ≥30× and ≥3 reads supporting each allele. Because no matched normal was available, genotypes were tumor-derived and imbalance was summarized as folded B-allele frequency, ∣AF−0.5∣. Sites were restricted to main chromosomes and anomalously dense 1-Mb windows were masked using a median + 5 scaled-MAD rule, removing 4,919 of 26,292 sites (18.7%); chromosome X was excluded from the scan. Segments were discovered de novo by chromosome arm using a rolling median over 51 consecutive informative sites. The screening threshold was estimated empirically as the 99.5th percentile of medians from size-matched contiguous runs sampled from the informative-site pool. Candidate segments were excluded and the background and threshold were re-estimated iteratively to convergence; the final background-folded BAF was 0.0952 and the threshold was 0.1981 after three passes. Adjacent runs separated by ≤20 sites and ≤2 Mb were merged, boundaries were refined by local CUSUM change-point analysis, and retained segments required ≥25 sites and ≥1 Mb. Segment significance was assessed against 4,000 size-matched contiguous-run null samples, with Benjamini– Hochberg correction, and genome-wide family-wise error was estimated from 500 block-bootstrap replicates using 10-site blocks. Depth support was evaluated independently from GC-corrected on-target log2 ratios using contiguous genomic null windows.

## Data availability

The processed data is available via Dryad (DOI: 10.5061/dryad.2ngf1vj56) and the raw data may be made available upon request to the corresponding author (H.M.L).

## Code availability

The analytical code can be found in https://github.com/heiminglai/REZCUE

## Statistic limitations

For whole genome sequencing comparison, with five pairs the exact Wilcoxon signed-rank test, and hence significance thresholds as well as effect size cannot be confirmed.

## Acknowledgement

We would like to express our deepest gratitude to the tissue donors and their family for their generosity and wisdom in supporting scientific studies. We thank Ka Wai Chan and Cathy Shuk Ling Chan for administrative support to the project. Part of the work was completed during JJXL’s affiliation with the Department of Anatomical and Cellular Pathology, CUHK, he can be reached currently at his listed affiliation. Figure 5A was partly created with BioRender.com.

## Author Contributions

Conceived the idea: HML, Developed REZCUE: LCZ, HML, Investigation: WCSC, JJXL, Performed the experiments: ZL, BPSP, BTYW, HML, Data analysis: EPLT, YT, HML, Administration: HML, Funding acquisition: YT, HML, Supervision: YT, HML. Writing: YT, HML.

## Supplementary Information

### Step-by-step REZCUE and INSIHGT2 protocol

#### Chemicals required

1. Doubly-distilled H_2_O (ddH_2_O)
2. Methylcyclopentane
3. Cyclohexane
4. Tetrahydrofuran (with or without stabilizers)
5. DMSO
6. Methanol
7. BABB (1:2 v/v benzyl alcohol:benzyl benzoate)
8. Sodium dodecahydro-*closo*-dodecaborate
9. Zwittergent SB3-16
10. α-cyclodextrin
11. Sulfobutyl-β-cyclodextrin
12. 2-Hydroxypropyl-γ-cyclodextrin

#### Apparatuses required

1. Shaking incubator capable of reaching temperatures of 55-60 °C. Water/metal bead-immersion shaking baths are preferred over air incubators for better heat equilibration.
2. 100-mL GL45 laboratory glass bottles (blue capped)
3. 50-mL conical tubes, made of polypropylene

##### Suitable containers for REZCUE and INSIHGT2

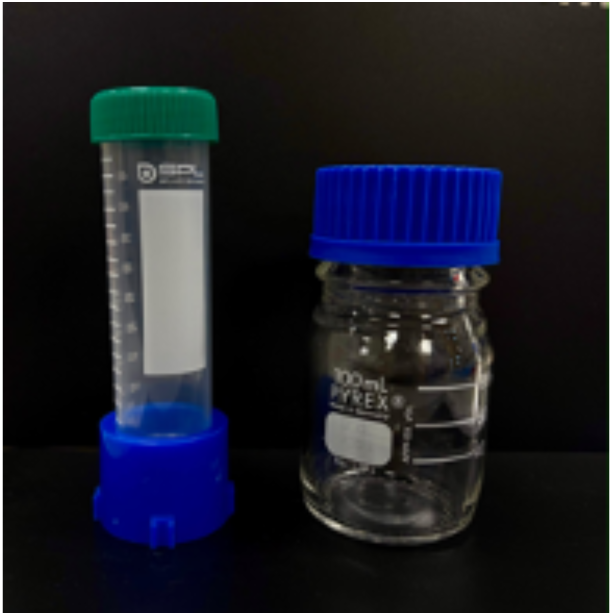

#### Stock solutions to be prepared

1. REZCUE stock A: 2.5 M sodium dodecahydro-*closo*-dodecaborate in ddH_2_O
2. REZCUE stock B: 0.3 M zwittergent SB3-16 in ddH_2_O
3. REZCUE stock C: 0.2 M α-cyclodextrin in ddH_2_O

- You may heat the solutions to 55 °C until all solid dissolves.
- These solutions are indefinitely stable at room temperature provided that there is no microbial contamination.
4. PBSN: 1x PBS with 0.02% w/v sodium azide

- Sodium azide is highly toxic, follow local institutional guidelines for safety. Do not acidify PBSN with strong acids as toxic hydrazoic acid fumes will be emitted.
5. 2x INSIHGT2 stock A: 0.5 M sodium dodecahydro-*closo*-dodecaborate in PBSN
6. 1x INSIHGT2 stock B: 0.25 M 2-hydroxypropyl-γ-cyclodextrin in 1x PBSN
7. 1x INSIHGT2 stock C: 0.1 M sulfobutyl-β-cyclodextrin in 1x PBSN

### REZCUE

This protocol assumes a tissue block of 5 mm x 3 cm x 2 cm. For smaller blocks, we recommend the same durations of incubations and volumes of solutions be used as a trial run. Scaling down reagent concentrations and volumes can be done after successful validation of the antibodies and tissue type according to this general protocol.

** Incubate at 20–25 °C unless otherwise specified.

** Use fresh solution for each repeat. Keep the specimen fully always submerged and agitate on an orbital shaker at 100 rpm, and place the tube horizontally to maximize fluid flow unless otherwise specified.

#### A.#Deparaffinization

1. Remove excess paraffin with a scalpel without cracking the tissue. Place the tissue into a 100-mL GL45 laboratory glass bottle.

2. Incubate in 25 mL methylcyclopentane (4 × 1 hour) to dissolve most of the paraffin.

3. Replace with 50 mL cyclohexane and incubate for 6–18 hours.

4. Replace with 50 mL fresh cyclohexane and incubate for 1 hour.

#### B.#Rehydration

5. Replace with 50 mL 85:15 v/v THF:methanol and incubate for 6–18 hours.

6. Transfer the tissue to a new 50-mL conical tube. Rehydrate sequentially in 25 mL 100% methanol (3 × 30 min), 25 mL 50% v/v methanol in H₂O (2 × 30 min), and 25 mL ddH₂O (2 × 30 min). Shake the tube horizontally.

#### C.#Aqueous antigen retrieval

7. Pre-warm aliquots of REZCUE stocks A-C and ddH₂O to 55 °C.

1. For each tissue, add the following components in the order shown (total 10 mL per tissue):
2. 1.5 mL ddH₂O
3. 5 mL REZCUE stock A
4. 1 mL REZCUE stock B
5. 2.5 mL REZCUE stock C

- Upon cooling, the mixture may form a white precipitate. If this happens, warm the closed tube to 55 °C until the precipitate redissolves completely; this may take up to 1 hour.

##### REZCUE aqueous mix at 55 °C (left) and room temperature (right)

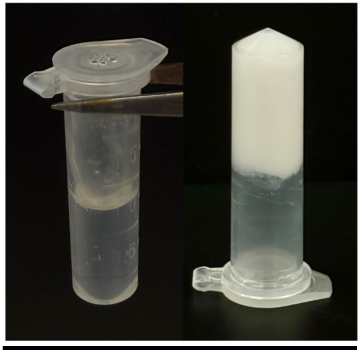

9. Agitate the tube upright at 55 °C for at least 12 hours, but not longer than 48 hours in total.

- Pre-warm at least 50 mL ddH₂O alongside each tissue for the two 25 mL washes in Step 10.
- If precipitates appear, increase the temperature to 60 °C for no more than 2 hours total. Once dissolved, return to 55 °C.
- Air incubators may equilibrate slowly. Use a water or metal-bead bath for better temperature control.

10. Wash the tissue in 25 mL ddH₂O at 55 °C (2 × 1 hour).

11. Wash the tissue in 25 mL 50% v/v DMSO in H₂O at room temperature (1 × 2 hours).

12. Wash the tissue in 25 mL ddH₂O (2 × 30 min). The tissue may proceed INSIHGT2 as with formalin-fixed tissue or be stored in 1× PBS with 0.02% w/v sodium azide at 4 °C for up to 1 year.

### INSIHGT2

This protocol assumes a 1 cm^3^ tissue block or an adult whole mouse brain. For smaller tissues, we recommend using the same durations of incubations and volumes of solutions be used as a trial run. Scaling down reagent concentrations and volumes can be done after successful validation of the antibodies and tissue type according to this general protocol.

**Incubate at 20–25 °C unless otherwise specified.

**Use fresh solution for each repeat. Keep the specimen fully always submerged and agitate on an orbital shaker at 100 rpm, and place the tube horizontally to maximize fluid flow unless otherwise specified.

#### A.#Tissue delipidation

1. Place the tissue in a 50-mL conical tube.

2. Dehydrate in 25 mL 50% v/v methanol in H₂O 2 × 30 min, then in 25 mL 100% methanol 3 × 30 min.

3. Delipidate in 25 mL 2:1 v/v dichloromethane:methanol for 6–18 hours in a fume hood.

4. Rehydrate and wash sequentially in 25 mL 100% methanol 3 × 30 min, 50% v/v methanol in H₂O 2 × 30 min, and PBSN 2 × 30 min.

5. Pre-incubate in 2 mL 1x INSIHGT2 stock A at 37 °C with the tube upright for 6– 18 hours.

- Prepare the 1× stock A by mixing equal volumes of 2x INSIHGT2 stock A and PBSN.

#### B1. Primary antibody dispersal assisted by superchaotrope

6. Discard the preincubation solution and add 1 mL 2x INSIHGT2 stock A.

7. Add 10 μg of each primary antibody or lectin per 1 cm³ of tissue or per whole adult mouse brain as a starting amount. Directly conjugated primary antibodies and lectins may be included.

- Reduce the amount only after validation for the individual probe and specimen type.
- You can add multiple species-compatible primary antibodies for 3D multiplexed immunohistochemistry. (Optional nuclear stain:) In a separate tube, mix 3 μL of 1 mM nuclear-dye stock with 3 μL 1x INSIHGT2 stock C; add the entire mixture to the staining solution.

8. Add PBSN or 2x INSIHGT2 stock A to reach a final 1x INSIHGT2 stock A concentration in the staining solution. To do so, let *V* (in mL) equal the combined volume added in Steps 7–8:

- If *V* < 1 mL, add (1 − *V*) mL PBSN.
- If *V* > 1 mL, add (*V* − 1) mL 2x INSIHGT2 stock A.

9. If necessary, add a minimal volume of 1x INSIHGT2 stock A with the same concentration of probes to fully submerge the tissue.

10. Incubate and agitate the tube upright at 20–25 °C for at least 4 days in the dark.

- If penetration is suboptimal, extending the incubation time to up to 14 days may help.

#### B2.#*In situ* host-guest chemistry - 1st stage

11. Incubate in 5 mL 1x INSIHGT2 stock B at 20–25 °C for 6–18 hours while agitating the tube upright.

12. Wash in 25 mL PBSN (3 × 2 hours), shake the tube horizontally.

- If all primary antibodies and lectins are directly fluorophore-conjugated, proceed directly to Step 21.

#### C1.#Secondary antibody dispersal assisted by superchaotrope

13. Pre-incubate in 2 mL 1x INSIHGT2 stock A at 20–25 °C for 6–18 hours while agitating the tube upright.

14. Discard the preincubation solution and add 1 mL 2x INSIHGT2 stock A.

15. Add secondary reagents based on the mass of each unconjugated primary antibody used in INSIHGT2 protocol Step 7.

- Add 1 μg of secondary IgG or Fab fragment per 1 μg of the corresponding primary antibody.
- Add 0.4 μg of nanobody per 1 μg of the corresponding primary antibody.

16. Add PBSN, 2x INSIHGT2 stock A, or 1x INSIHGT2 stock A, using the dilution and coverage rules in INSIHGT2 protocol Steps 9–10.

17. Incubate and agitate the tube upright at 20–25°C for 2–3 days in the dark.

#### B2.#*In situ* host-guest chemistry - 2nd stage

18. Incubate in 5 mL 1x INSIHGT2 stock B at 20–25 °C for 6–18 hours while agitating the tube upright.

19. Wash in 25 mL PBSN (3 × 2 hours), shake the tube horizontally.

20. Dehydrate in 25 mL 50% v/v methanol in H₂O 2 × 30 min, then in 25 mL 100% methanol 3 × 30 min.

21. Clear in 20 mL BABB consisting of benzyl alcohol:benzyl benzoate at 1:2 v/v. Replace with 20 mL fresh clearing solution after 3 hours and continue for at least 6 additional hours, or until the tissue is optically clear.

- BABB is highly recommended for human tissues, especially for tissues retrieved from FFPE.

### Tissue recovery for downstream omics studies – as FFPE tissue block

1. Place the BABB-cleared tissue into a 100-mL GL45 laboratory glass bottle.
2. Wash the BABB-cleared tissue in 25 mL 100% methanol for 12-18 hours or overnight.
3. Wash sequentially in 25 mL 100% methanol for 2 × 30 min, then in 25 mL 100% ethanol for 3 × 30 min, then in 25 mL xylene for 2 × 30 min, then in fresh 25 mL xylene for 12-18 hours or overnight.
4. Put the tissue into an acetal copolymer tissue cassette. Permeate the tissue in a 60 °C hot bath of paraffin wax in a tissue preparation station for 12-18 hours or overnight.
5. Proceed to usual paraffin wax embedding. The subsequently obtained FFPE block can be used for sectioning and downstream omics assays.

### Tissue recovery for downstream omics studies – as rehydrated tissue volume

1. Place the BABB-cleared tissue into a 50-mL conical tube.
2. Wash the BABB-cleared tissue in 25 mL 100% methanol for 12-18 hours or overnight.
3. Wash sequentially in 25 mL 100% methanol for 2 × 30 min, then in 25 mL 50% v/v methanol in H₂O for 3 × 30 min, then in water for 2 × 30 min, and finally in PBSN for 2 × 30 min.
4. The tissue can be re-used as a formalin-fixed tissue or placed in a tissue processor for automated dehydration and paraffin embedding.

#### Tissue physical appearance at different stages of REZCUE and INSIHGT2 protocol

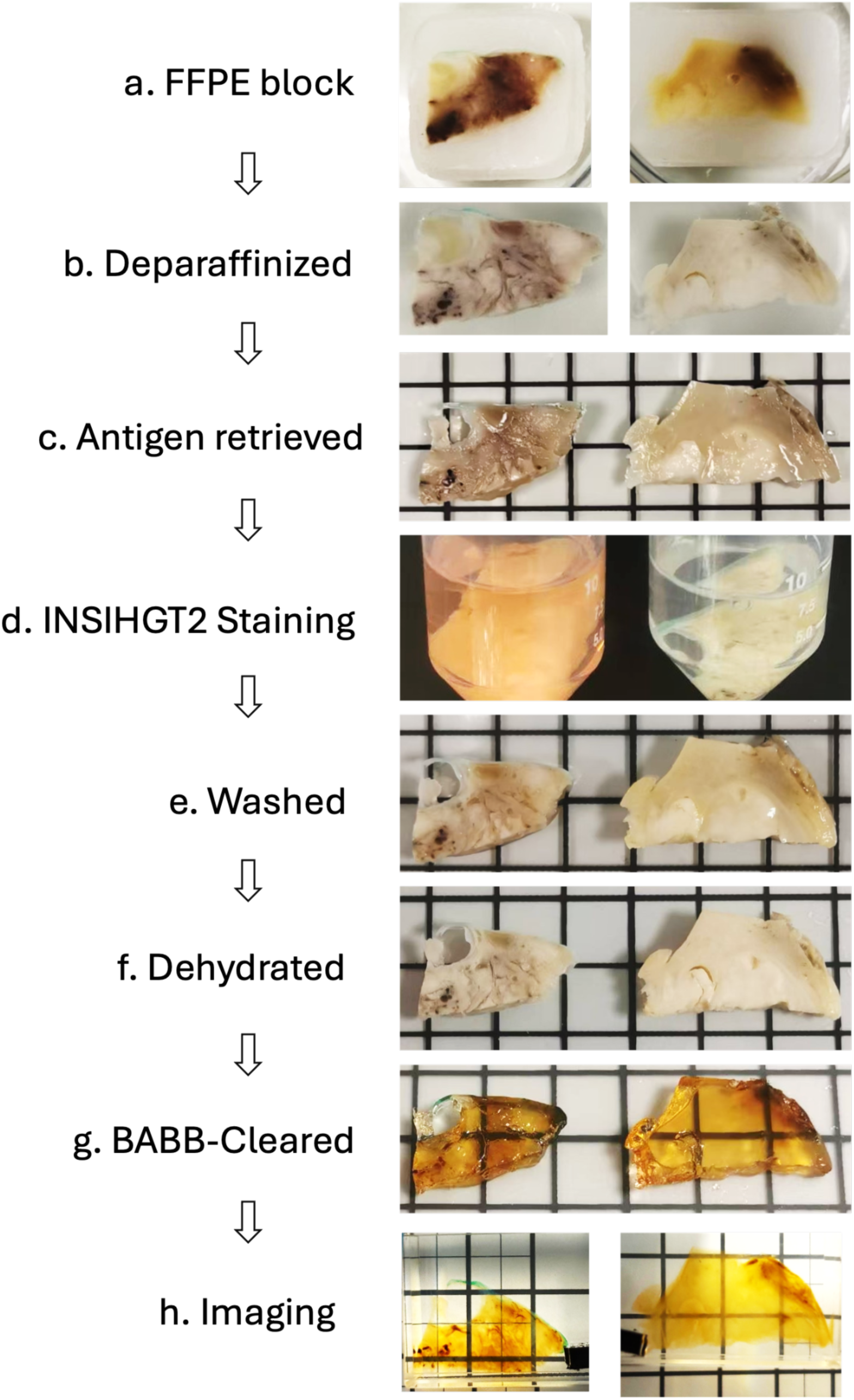

## Supplementary notes

### Note 1. Achieving antigen retrieval as a global phase change in tissue

The completeness of deparaffinization is critical as any residual paraffin wax pocket will create a local hydrophobic environment, jeopardizing the next stage of protein reconfiguration in the aqueous environment. Paraffin residues create hydrophobic pockets that expose hydrophobic residues in adjacent proteins, which, in turn, promote the exposure of hydrophobic residues to these neighboring proteins. If proteins are assumed to have a binary hydrophobic-exposed or hydrophilic-exposed state, this is akin to an Ising model with an underlying random scalar hydrophobicity field that describes the environmental influence on the protein’s folding propensity in situ.

We identified three major problems with 3D antigen retrieval: (1) the incomplete deparaffinization of tissues, (2) the reprecipitation of wax during tissue rehydration, and (3) the inefficient re-exposure of hydrophilic epitopes. Notably, we found that xylene is a poor extracting solvent for paraffin wax. Intriguingly, *n*-hexane also performed poorly, even though it has a small molecular weight with near-ideal enthalpic interaction with paraffin wax molecules. This can be explained by the relationship between molecular geometry and dissolution entropy, as the elongated *n*-hexane molecules will preferably align with the wax molecules for optimal packing, trading orientational entropy for translational entropy. Based on such theoretical considerations, a smaller isomer with similar intermolecular interaction strength would be beneficial. Indeed, a switch to cyclohexane led to fast and efficient paraffin wax dissolution (**Supplementary Note 2**), with methylcyclopentane dissolving paraffin wax twice as fast as cyclohexane but with slightly reduced total wax solubility. The smaller cyclopentene dissolved the paraffin wax even more rapidly, but its low boiling point and relatively high cost render it a less optimal choice. Thus, we employ a fast wax dissolution step in methylcyclopentane, followed by a cyclohexane incubation step in which paraffin wax is most soluble.

After wax dissolution, the tissue is transitioned through increasingly hydrophilic solvent blends for rehydration, and it is crucial to avoid wax reprecipitation within the dense tissue matrix, as the tissue will be incubated in aqueous environment till the end of immunostaining. We thus tested for a water-miscible transition solvent blend that would minimize wax precipitation when mixed with wax-saturated cyclohexane. After extensive optimization, we arrived at an 85/15 v/v mixture of tetrahydrofuran and methanol. This solvent mixture dissolves a considerable amount of paraffin wax and is phase-stable when mixed with cyclohexane or water, which are properties ideal for a transitioning solvent washing step. The tissue can then be rehydrated in graded methanol to achieve complete rehydration.

We next sought to develop the best epitope recovery method to enable downstream 3D multiplexed immunohistochemistry. We tested organic phase and aqueous phase epitope re-exposure conditions, conducted before and after graded methanol rehydration, respectively (**Supplementary Figure 2**). Most failed. Serendipitously, we observed in our previous development of INSIHGT ^51^ that a zwitterionic detergent-based artificial chaperone was able to improve immunostaining quality (**Supplementary Figure 2**). As chaotropes also facilitate the solubilization of proteins, we tested a mixture of an alkyl sulfobetaine zwitterionic detergent and the superchaotrope [B_12_H_12_]^2-^. This mixture becomes semi-solid when left at room temperature, which is reversible through heating and supramolecular chemical means by adding γ-cyclodextrin. FFPE tissues incubated in this solution at 55 °C for more than 2 hours showed completely recovered antibody- and lectin-based staining quality, which is indistinguishable from non-FFPE controls (**Supplementary Figure 5C**).

### Note 2. Entropic considerations in paraffin wax dissolution

Paraffin wax is a heterogeneous mixture of mostly saturated aliphatic hydrocarbons with minimal branched chains. The drastically improved performance of cyclohexane compared to *n*-hexane and xylene can be understood through entropic gains of the paraffin wax molecules ^61^. Due to the hydrocarbon nature of the solute and the solvent, it is also reasonable to assume the enthalpic contributions, mostly due to dispersion forces, are similar across all cases. The internal energies of the system also did not change. In this case, dissolution happens spontaneously when the Helmholtz free energy change Δ*F* is negative. ΔF is related to the entropy changes Δ*S* through the following equation:

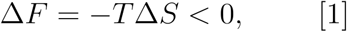

where the total entropy change is decomposed into translational and orientational entropies of the molecules is

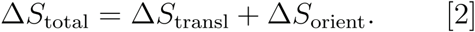

Where Δ*S*_transl_ is the translational entropy and Δ*S*_orient_ is the orientational entropy. *We* can consider paraffin wax as a mixture of long rod-shaped molecules in a confined, ordered state, in semi-crystalline form (solid nematic phase), arranged parallel to each other along their long axes. When the paraffin wax molecules go from the solid nematic phase to full dissolution in with isotropic arrangement, the maximal translational entropic gain for the wax molecules is

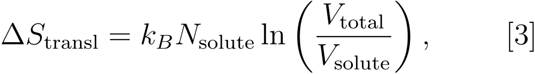

Where *k_B_* is Boltzmann’s constant, *N* is the number of solute molecules, and *V* is the volume. The orientational entropic gain is

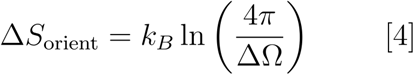

where the orientationally confined rod-shaped molecules go from a solid angle ΔΩ to a full 4π steradian.

Solvent molecules of different geometries can be introduced into the system by formulating them as shorter rod-shaped molecules of diameter *d* and length *l*. Consider hexane, cyclohexane, and benzene (as a 6-carbon surrogate to xylene). In isotropic fluid, the excluded volumes of each molecule are simply their largest diameter sweeping out a spherical volume of π*D*^3^/6.

In the case of hexane and benzene molecules, where *d* ≠ *l*, the optimal packing of hexane and benzene molecules is nematic ^62–64^ and stacked ^62–64^, respectively. However, this sacrifices the orientational degree of freedom around the wax molecules, leading to a reduction in 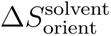 trading for maximal 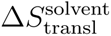. Alternatively, it can also be viewed that the dissolution of wax molecules with random orientation would disrupt the nematic or stacking arrangement of hexane or benzene, respectively. This results in an effective excluded volume 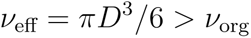, where *D* is the molecule’s largest diameter and *v* is the molecular volume. Since *N* = *V/v*, from equation [3], the Δ*S* term will also decrease. If the overall system volume increases as a result, it will introduce a *P_atm_*Δ*V* term into the Helmholtz free energy in equation [1] for work done against the atmospheric pressure. At isobaric conditions, a positive Δ*H* term would be introduced to the free energy change, due to decrease in local cohesive dispersion forces. Overall, these effects mean that the near-spherical cyclohexane molecules are the best solvent for paraffin wax. There are many options for spheroid-shaped solvent molecules with molecular weights similar to cyclohexane. We carried out paraffin wax dissolution experiments to determine the best solvent, constrained by (1) a boiling point of >55 °C, (2) chemical stability, and (3) minimal health hazards. We tested 6-carbon and 5-carbon solvents and determined that the optimal solvents for dissolution of paraffin are:

cyclopentene > methylcyclopentane ∼ cyclohexene ∼ cyclohexa-1,4-diene > cyclohexa-1,3-diene > cyclohexane > neohexane > n-hexane

From this trend, it is apparent that the wax dissolution kinetics and stability are thus influenced by a combination of solvent molecular weight, wax packing disruption due to the 3D shape of the solvent molecules, and solvent molecule surface area for van der Waals interaction with wax molecules. These solvent factors, in turn, govern the solvent diffusivity, dissolution entropies, and dissolution enthalpies, respectively. We ultimately chose methylcyclopentane and cyclohexane for their stability, safety, and performance.

